# Genetic, Clinical Underpinnings of Brain Change Along Two Neuroanatomical Dimensions of Clinically-defined Alzheimer’s Disease

**DOI:** 10.1101/2022.09.16.508329

**Authors:** Junhao Wen, Zhijian Yang, Ilya M. Nasrallah, Yuhan Cui, Guray Erus, Dhivya Srinivasan, Ahmed Abdulkadir, Elizabeth Mamourian, Ioanna Skampardoni, Gyujoon Hwang, Ashish Singh, Mark Bergman, Jingxuan Bao, Erdem Varol, Zhen Zhou, Aleix Boquet-Pujadas, Jiong Chen, Arthur W. Toga, Andrew J. Saykin, Timothy J. Hohman, Paul M. Thompson, Sylvia Villeneuve, Randy Gollub, Aristeidis Sotiras, Katharina Wittfeld, Hans J. Grabe, Duygu Tosun, Murat Bilgel, Yang An, Daniel S. Marcus, Pamela LaMontagne, Tammie L. Benzinger, Susan R. Heckbert, Thomas R. Austin, Lenore J. Launer, Mark Espeland, Colin L Masters, Paul Maruff, Jurgen Fripp, Sterling C. Johnson, John C. Morris, Marilyn S. Albert, R. Nick Bryan, Susan M. Resnick, Luigi Ferrucci, Yong Fan, Mohamad Habes, David Wolk, Li Shen, Haochang Shou, Christos Davatzikos, iSTAGING, the AI4AD, and the ADSP phenotypic harmonization consortia, the BLSA, the PREVENT-AD, and the ADNI studies

## Abstract

Alzheimer’s disease (AD) is associated with heterogeneous atrophy patterns. We employed a semi-supervised clustering technique known as Surreal-GAN, through which we identified two dominant dimensions of brain atrophy in symptomatic mild cognitive impairment (MCI) and AD patients: the “diffuse-AD” (R1) dimension shows widespread brain atrophy, and the “MTL-AD” (R2) dimension displays focal medial temporal lobe (MTL) atrophy. Critically, only R2 was associated with widely known sporadic AD genetic risk factors (e.g., *APOE ε4*) in MCI and AD patients at baseline. We then independently detected the presence of the two dimensions in the early stages by deploying the trained model in the general population and two cognitively unimpaired cohorts of asymptomatic participants. In the general population, genome-wide association studies found 77 genes unrelated to *APOE* differentially associated with R1 and R2. Functional analyses revealed that these genes were overrepresented in differentially expressed gene sets in organs beyond the brain (R1 and R2), including the heart (R1) and the pituitary gland, muscle, and kidney (R2). These genes were enriched in biological pathways implicated in dendritic cells (R2), macrophage functions (R1), and cancer (R1 and R2). Several of them were “druggable genes” for cancer (R1), inflammation (R1), cardiovascular diseases (R1), and diseases of the nervous system (R2). The longitudinal progression showed that *APOE ε4*, amyloid, and tau were associated with R2 at early asymptomatic stages, but this longitudinal association occurs only at late symptomatic stages in R1. Our findings deepen our understanding of the multifaceted pathogenesis of AD beyond the brain. In early asymptomatic stages, the two dimensions are associated with diverse pathological mechanisms, including cardiovascular diseases, inflammation, and hormonal dysfunction – driven by genes different from *APOE* – which may collectively contribute to the early pathogenesis of AD.

## Introduction

Alzheimer’s disease (AD) is the most common cause of dementia in older adults and remains incurable despite many pharmacotherapeutic clinical trials, including anti-amyloid drugs^1,2^ and anti-tau drugs.^3^ This is largely due to the complexity and multifaceted nature of the underlying neuropathological processes leading to dementia. The research community has embraced several mechanistic hypotheses to elucidate AD pathogenesis.^4–6^ Among these, the amyloid hypothesis has been dominant over the past decades and has proposed a dynamic biomarker chain: extracellular beta-amyloid (Aβ) triggers a cascade that leads to subsequent intracellular neurofibrillary tangles, including hyperphosphorylated tau protein (tau and p-tau), neurodegeneration, including medial temporal lobe atrophy, and cognitive decline.^7,8^ However, the amyloid hypothesis has been reexamined and revised due to substantial evidence which questions its current form^8–10^. While amyloid remains critical to AD development, the amyloid cascade model has been continually refined as other biological factors are discovered to influence the pathway from its accumulation to cell death.

Cardiovascular dysfunction has been widely associated with an increased risk for AD.^11^ There is also growing evidence that inflammatory^10–12^ and neuroendocrine processes^5,13^ influence pathways of amyloid accumulation and neuronal death. The inflammation hypothesis claims that microglia and astrocytes release pro-inflammatory cytokines as drivers, byproducts, or beneficial responses associated with AD progression and severity.^12–14^ The neuroendocrine hypothesis, first introduced in the context of aging^15^, has been extended to AD^16^, where it proposes that neurohormones secreted by the pituitary and other essential endocrine glands can affect the central nervous system (CNS), which subsequently contribute to developing AD. For example, Xiong and colleagues^17^ recently found that blocking the action of follicle-stimulating hormone in mice abrogates the AD-like phenotype (e.g., cognitive decline) by inhibiting the neuronal C/EBPβ–δ-secretase pathway. These findings emphasize the need to further elucidate early brain and body changes well before they lead to irreversible clinical progression.^18^

Recent advances in artificial intelligence (AI), especially deep learning (DL), applied to magnetic resonance imaging (MRI), showed great promise in biomedical applications^19,20^. DL models discover complex non-linear relationships between phenotypic and genetic features and clinical outcomes, thereby providing informative imaging-derived endophenotypes^21^. In particular, AI has been applied to MRI to disentangle the neuroanatomical heterogeneity of AD with categorical disease subtypes.^22–26^ The genetic underpinnings^27,28^ of this neuroanatomical heterogeneity in AD are also complex and heterogeneous. The most recent large-scale genome-wide association study^27^ (GWAS: 111,326 AD vs. 677,633 controls) has identified 75 genomic loci, including *APOE* genes, associated with AD. However, such case-control group comparisons conceal genetic factors that might contribute differentially to different dimensions of AD-related brain change. More importantly, the genetic variants that contribute to the initiation and early progression of brain change in younger and asymptomatic individuals are poorly understood.

In this study, we utilize a novel semi-supervised deep learning approach, Surreal-GAN, to characterize the neuroanatomical heterogeneity of the disease. Unlike our previous model, Smile-GAN^22^, which categorized subtypes, Surreal-GAN generates multiple continuous dimensional scores, simultaneously accounting for spatial and temporal disease heterogeneity, similar to what was accomplished in a previous unsupervised clustering model known as Sustain. These multi-dimensional scores reflect the co-expression level of respective brain atrophy dimensions in the same patient. Refer to the method (**Surreal-GAN for disease heterogeneity**) and **Supplementary eMethod 1** for methodological details of Surreal-GAN, comparisons to other subtyping methods, and strengths of semi-supervised representation learning. We hypothesized that genetic variants, potentially unrelated to *APOE* genes, contribute to early manifestations of multiple dimensions of brain atrophy in early asymptomatic stages. We first trained the Surreal-GAN model to define the AD dimensions to test this hypothesis in the late symptomatic stages. We then examined their expression back to early asymptomatic stages. In our previous study,^29^ we derived two neuroanatomical dimensions (R1 and R2) by applying Surreal-GAN to the MCI/AD participants (target population) and cognitively unimpaired (CU) participants (reference population) from the Alzheimer’s Disease Neuroimaging Initiative study (ADNI^30^). Herein, we applied the trained model to three asymptomatic populations and one symptomatic population: the *general population* (*N*=39,575; age: 64.12±7.54 years) from the UK Biobank (UKBB^31^) excluding demented individuals; the *cognitively unimpaired population* (*N*=1658; age: 65.75±10.90 years) from ADNI and the Baltimore Longitudinal Study of Aging study (BLSA^32^); the *cognitively unimpaired population with a family risk* (*N*=343; age: 63.63±5.05 years) from the Pre-symptomatic Evaluation of Experimental or Novel Treatments for Alzheimer’s Disease (PREVENT-AD^33^); the *MCI/AD population* (*N*=1534; age: 73.45±7.69 years) from ADNI and BLSA. Refer to the method (**Study design and populations**) and **Table 1** for details of the definition of these populations.

## Materials and methods

### Study design and populations

The current study consists of four main populations (**Table 1**), which were jointly consolidated by the iSTAGING^34^ and the AI4AD consortia (http://ai4ad.org/): the iSTAGING consortium consolidated all imaging and clinical data; imputed genotyping data were downloaded from UKBB; the AI4AD consortium consolidated the whole-genome sequencing (WGS) data for the ADNI study. The definition of each population and inclusion criteria are detailed in **Supplementary eMethod 2**.

### Image preprocessing

All T1w-weighted MR images were first corrected for magnetic field intensity inhomogeneity.^35^ A deep learning-based skull stripping algorithm was applied for the removal of extra-cranial material. In total, 145 anatomical regions of interest (ROIs) were generated in gray matter (GM, 119 ROIs), white matter (WM, 20 ROIs), and ventricles (6 ROIs) using a multi-atlas label fusion method^36^ (**Supplementary eMethod 3**). The 119 ROIs were statistically harmonized by an extensively validated approach, i.e., ComBat-GAM^37^, using the entire imaging data of iSTAGING. **Supplementary eFigure 1** demonstrates the normality check for the MUSE ROI (right accumbent area) before and after statistical harmonization, illustrating that our statistical harmonization enhanced the normality of ROIs across various studies.

### Surreal-GAN for disease heterogeneity

Surreal-GAN^29^ dissects underlying disease-related heterogeneity via a deep representation learning approach under the principle of semi-supervised learning.^22,23^ At a high level, its most fundamental novelty is that it provides a continuous representation of the presence of multiple, non-exclusive abnormal brain patterns in each individual, rather than clustering individuals into one of many clusters, i.e. disease subtypes. More specifically, several methodological advancements were considered compared to its predecessor, Smile-GAN^22^. First, Surreal-GAN is to model neuroanatomical heterogeneity by considering both spatial and temporal (i.e., disease severity) variation using only cross-sectional MRI data. Secondly, Surreal-GAN disentangles the neuroanatomical heterogeneity of AD by enabling patients to simultaneously exhibit multiple distinct imaging patterns (i.e., high scores for expressing all these patterns), resulting in high-dimensional scores across multiple dimensions. Lastly, in contrast to prior probability-based clustering methods like Smile-GAN, Surreal-GAN operates without the constraint that all dimensional scores must sum to 1. This allows for a more normal distribution of dimensional scores suited for GWAS (**Supplementary eFigure 2)**. Further methodological details are elaborated upon in **Supplementary eMethod 1**.

Alternative clustering techniques, such as Sustain^24^ and Bayesian latent^38^ methods, are available for deciphering the neuroanatomical heterogeneity in AD^22–26,39^. Surreal-GAN distinguishes itself from these approaches based on fundamental methodological distinctions, such as its utilization of semi-supervised deep learning compared to the unsupervised approach of others^40^. Additionally, Surreal-GAN generates continuous dimensions associated with distinct phenotypic outcomes, allowing the simultaneous co-expression of multiple patterns instead of categorizing patients into a single dominant subtype or stage, as seen in other methods. Notably, the two dimensions, R1 and R2, displayed correlations with the four subtypes generated by Smile-GAN, particularly R2 exhibited a correlation with P3 (reflecting medial temporal lobe atrophy), and both R1 and R2 displayed correlations with P4 (representing global atrophy), as depicted in **Supplementary eFigure 3**.

### Brain and clinical variable associations

We performed brain-wide associations for the 119 GM ROIs. For baseline brain-wide associations, linear regression models were fitted with R1 and R2 dimensions as independent variables, with each ROI as the dependent variable, controlling for age, sex, intracranial volume (ICV), and/or diagnosis as covariates.

We performed a two-step linear regression for longitudinal brain-wide associations. First, we derived the individual-level age change rate using a linear mixed-effects model. To this end, we included a participant-specific random slope for age and random intercept; age and sex were treated as fixed effects. Secondly, the same linear regression model as in baseline brain associations was fitted with the age change rate as the independent variable.

We also performed clinical variable association for all clinical variables and neuropsychological testing available for each population, using the same model in the baseline brain-wide associations. Bonferroni correction of 119 GM ROIs was performed to adjust for the multiple comparisons.

### Genetic analyses

Genetic analyses were performed for the WGS data from ADNI and the imputed genotype data from UKBB. Our quality check protocol is detailed in **Supplementary eMethod 4**. This resulted in 1487 participants and 24,194,338 SNPs in ADNI WGS data. For UKBB, we limited our analysis to European ancestry participants, resulting in 33,541 participants and 8,469,833 SNPs.

Using UKBB data, we first estimated the SNP-based heritability using GCTA-GREML,^41^ controlling for confounders of age (at imaging), age-squared, sex, age-sex interaction, age-squared-sex interaction, ICV, and the first 40 genetic principal components, following a previous pioneer study.^42^ In GWAS, we performed a linear regression for each neuroanatomical dimension and included the same covariates as in the heritability estimates. We adopted the genome-wide P-value threshold (5 × 10^−8^) in all GWAS. The annotation of genomic loci (displayed by its top lead SNP) and gene mappings, prioritized gene set enrichment, and tissue specificity analyses were performed using FUMA (https://fuma.ctglab.nl/, version: v1.3.8) (**Supplementary eMethod 5** and **6**). A two-step procedure (**Supplementary eMethod 7**) was performed to determine if an annotated genomic locus or gene was associated with AD-related clinical traits. We calculated the polygenic risk scores (PRS)^43^ using both ADNI and UKBB genetic data (**Supplementary eMethod 8**). Finally, we constructed a target-drug-disease network for these genes associated with R1 and R2 to identify these “druggable genes” (**Supplementary eMethod 9**).

## Results

### Two dominant dimensions of brain atrophy found in MCI and AD

In MCI/AD patients, the “diffuse-AD” dimension (R1) showed widespread brain atrophy without an exclusive focus on the medial temporal lobe (**Fig. 1A** and **Supplementary eTable 1** for P-values and effect sizes). In contrast, the “MTL-AD” dimension (R2) displayed more focal medial temporal lobe atrophy, prominent in the bilateral parahippocampal gyrus, hippocampus, and entorhinal cortex (**Fig. 1A**). All results, including P-values and effect sizes (Pearson’s correlation coefficient *r*), are presented in **Supplementary eTable 1**. The atrophy patterns of the two dimensions defined in the symptomatic MCI/AD population (**Fig. 1A**) were present in the asymptomatic populations, albeit with a smaller magnitude of *r*. (**Supplementary eTable 1**, **4**, and **8**). We presented the age distribution (**Supplementary eFigure 4A**), as well as the expression of R1 and R2, along with the population-level difference for the four populations in **Supplementary eFigure 4B-D**.

**Figure 1:**
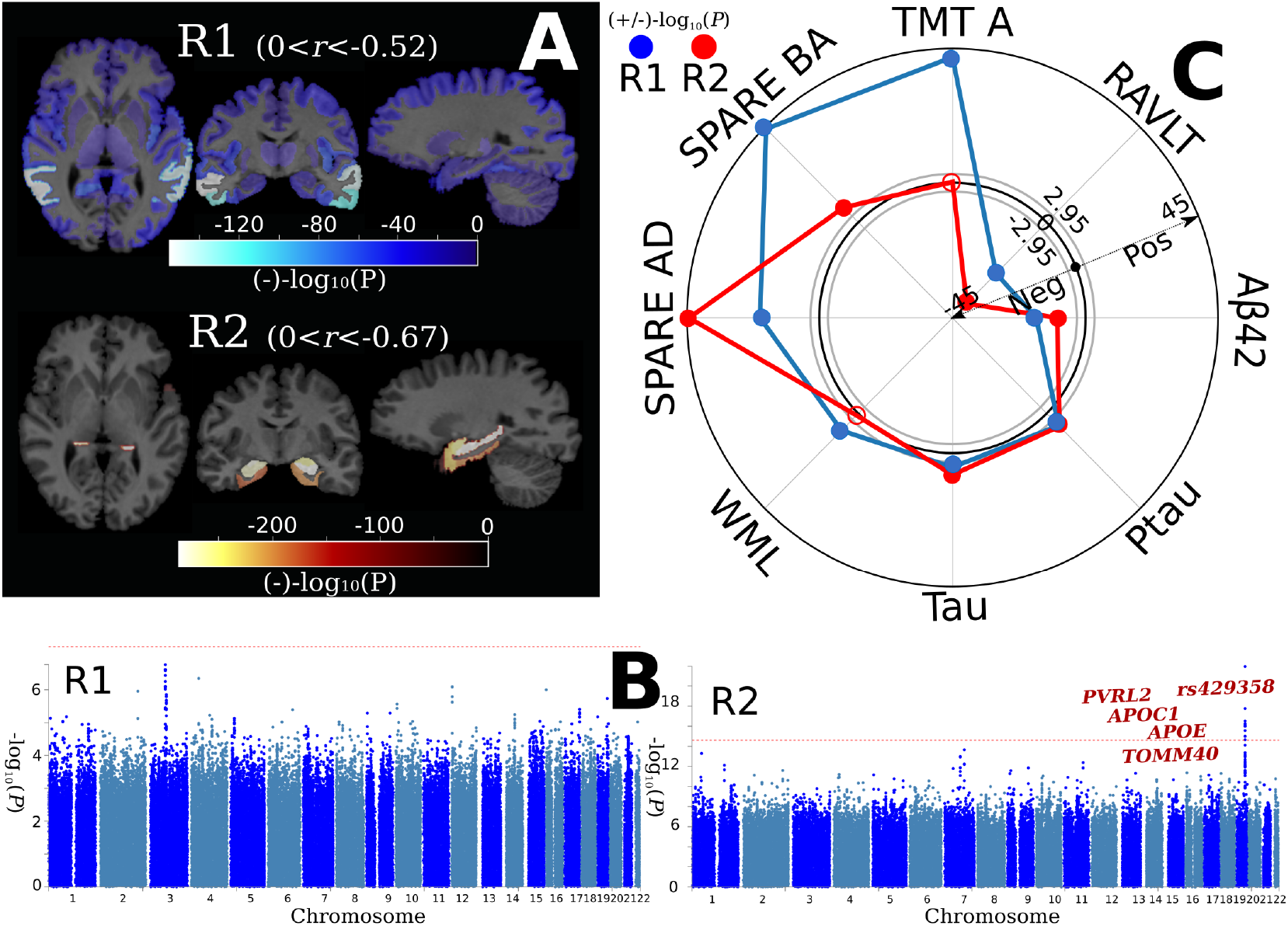
The manifestation of the R1 and R2 dimensions of brain atrophy in the MCI/AD population. **A)** Brain association studies reveal two dominant brain atrophy dimensions. A linear regression model was fit to the 119 GM ROIs at baseline for the R1 and R2 dimensions. The –log_10_(P-value) of each significant ROI (Bonferroni correction for the number of 119 ROIs: – log_10_(P-value) > 3.38) is shown. A negative value denotes brain atrophy with a negative coefficient in the linear regression model. P-value and effect sizes (*r*, Pearson’s correlation coefficient) are presented in **Supplementary eTable 1**. The range of *r* for each dimension is also shown. Of note, the sample size (*N*) for R1 and R2 is the same for each ROI. **B)** Genome-wide association studies demonstrate that the R2, but not R1, dimension is associated with variants related to *APOE* genes (genome-wide P-value threshold with the red line: –log_10_(P-value) > 7.30). We associated each common variant with R1 and R2 using the whole-genome sequencing data from ADNI. Gene annotations were performed via positional, expression quantitative trait loci, and chromatin interaction mappings using FUMA.^53^ We then manually queried whether they were previously associated with AD-related traits in the GWAS Catalog.^50^ Red-colored loci/genes indicate variants associated with AD-related traits in previous literature. **C)** Clinical association studies show that the R2 dimension is associated to a larger extent with AD-specific biomarkers, including SPARE-AD^44^, an imaging surrogate to AD atrophy patterns, and *APOE ε4,* the well-established risk allele in sporadic AD. The R1 dimension is associated to a larger extent with aging (e.g., SPARE-BA,^47^ an imaging surrogate for brain aging) and vascular-related biomarkers (e.g., WML, white matter lesion). The same linear regression model was used to associate the R1 and R2 dimensions with the 45 clinical variables, including cognitive scores, modifiable risk factors, CSF biomarkers, disease/condition labels, demographic variables, and imaging-derived phenotypes. The radar plot shows representative clinical variables; results for all 45 clinical variables are presented in **Supplementary eTable 3**. The SPARE-AD and SPARE-BA scores are rescaled for visualization purposes. The gray-colored circle lines indicate the P-value threshold in both directions (Bonferroni correction for the 45 variables: –log_10_(P-value) > 2.95). A positive/negative –log_10_(P-value) value indicates a positive/negative correlation (beta). The transparent dots represent the associations that do not pass the Bonferroni correction; the blue-colored dots and red-colored dots indicate significant associations for the R1 and R2 dimensions, respectively.

At baseline, the R1-dominant group had 25.72 % AD patients (*N*=222 out of 863); the R2-dominant group consisted of 30.10% AD patients (202 out of 671). Within a 7-year follow-up period, MCI participants from both the R1-dominant and R2-dominant groups progressed to AD, with the R2-dominant group exhibiting a higher proportion of AD patients (40% vs. 25%) (**Supplementary eFigure 5 A-B**); the two dominant dimensions developed independently throughout the 7-year follow-up period (**Supplementary eFigure 5 C-D**).

### *APOE* genes are associated with R2 but not with R1 in the MCI/AD population

In GWAS, the R2 dimension, but not R1, was associated with well-established AD genomic loci (*rs429358*, chromosome: 19, 45411941; minor allele: C, P-value: 1.05 × 10^−11^) and genes (*APOE*, *PVRL2*, *TOMM40*, and *APOC1*) (**Fig. 1B**). The details of the identified genomic locus and annotated genes are presented in **Supplementary eTable 2**. The PRS of AD showed a slightly stronger positive association with the R2 dimension [*r*=0.11, −log_10_(P-value)=3.14] than with the R1 dimension [*r*=0.09, −log_10_(P-value)=2.31, **Supplementary eFigure 6**]. The QQ plots of the baseline GWAS are presented in **Supplementary eFigure 7**.

### Clinical profiles of the R1 and R2 dimensions in the MCI/AD population

Clinical association studies correlated the two dimensions with 45 clinical variables and biomarkers. Compared to the R1 dimension, R2 showed associations, to a larger extent than R1, with SPARE-AD and the Rey Auditory Verbal Learning Test (RAVLT). SPARE-AD quantifies the presence of a typical imaging signature of AD-related brain atrophy, which has been previously shown to predict clinical progression in both CU and MCI individuals.^44^ RAVLT measures episodic memory, a reliable neuropsychological phenotype in AD, which is also correlated with medial temporal lobe atrophy.^45,46^ The R1 dimension was associated to a greater extent with 1) SPARE-BA, which captures the individualized expression of advanced brain age from MRI^47^; 2) white matter lesions (WML), which are commonly associated with vascular risk factors and cognitive decline^48^, and 3) whole-brain uptake of 18F-fluorodeoxyglucose (FDG) PET, which is a biomarker of brain metabolic function and atrophy. Both dimensions were positively associated with cerebrospinal fluid (CSF) levels of tau and p-tau and negatively associated with the CSF level of Aβ42^49^ (**Fig. 1C**), as well as the whole-brain standardized uptake value ratio of 18F-AV-45 PET (**Supplementary eTable 3**). Results for all 45 clinical variables, including cognitive scores, modifiable risk factors, CSF biomarkers, disease/condition labels, demographic variables, and imaging-derived phenotypes, are presented in **Supplementary eTable 3** for P-values and effect sizes (i.e., beta coefficients).

### Clinical profiles of the R1 and R2 dimensions in the general population

Brain association studies confirmed the presence of the two atrophy patterns in the general population (**Fig. 2A** and **Supplementary eTable 4** for P-values and effect sizes). In clinical association studies, the R1 dimension was significantly associated, to a larger extent than R2, with cardiovascular (e.g., triglycerides) and diabetes factors (e.g., Hba1c and glucose), executive function (TMT-B), intelligence, physical measures (e.g., diastolic blood pressure), SPARE-BA [–log_10_(P-value) = 236.89 for R1 and −46.35 for R2] and WML [–log_10_(P-value) = 120.24 for R1 and 2.06 for R2]. In contrast, the R2 dimension was more significantly associated with SPARE-AD [–log_10_(P-value) = 136.01 for R1 and 250.41 for R2] and prospective memory (**Fig. 2B**). Results for all 61 clinical variables, including cardiovascular factors, diabetic blood markers, social demographics, lifestyle, physical measures, cognitive scores, and imaging-derived phenotypes, are presented in **Supplementary eTable 5** for P-values and effect sizes.

**Figure 2:**
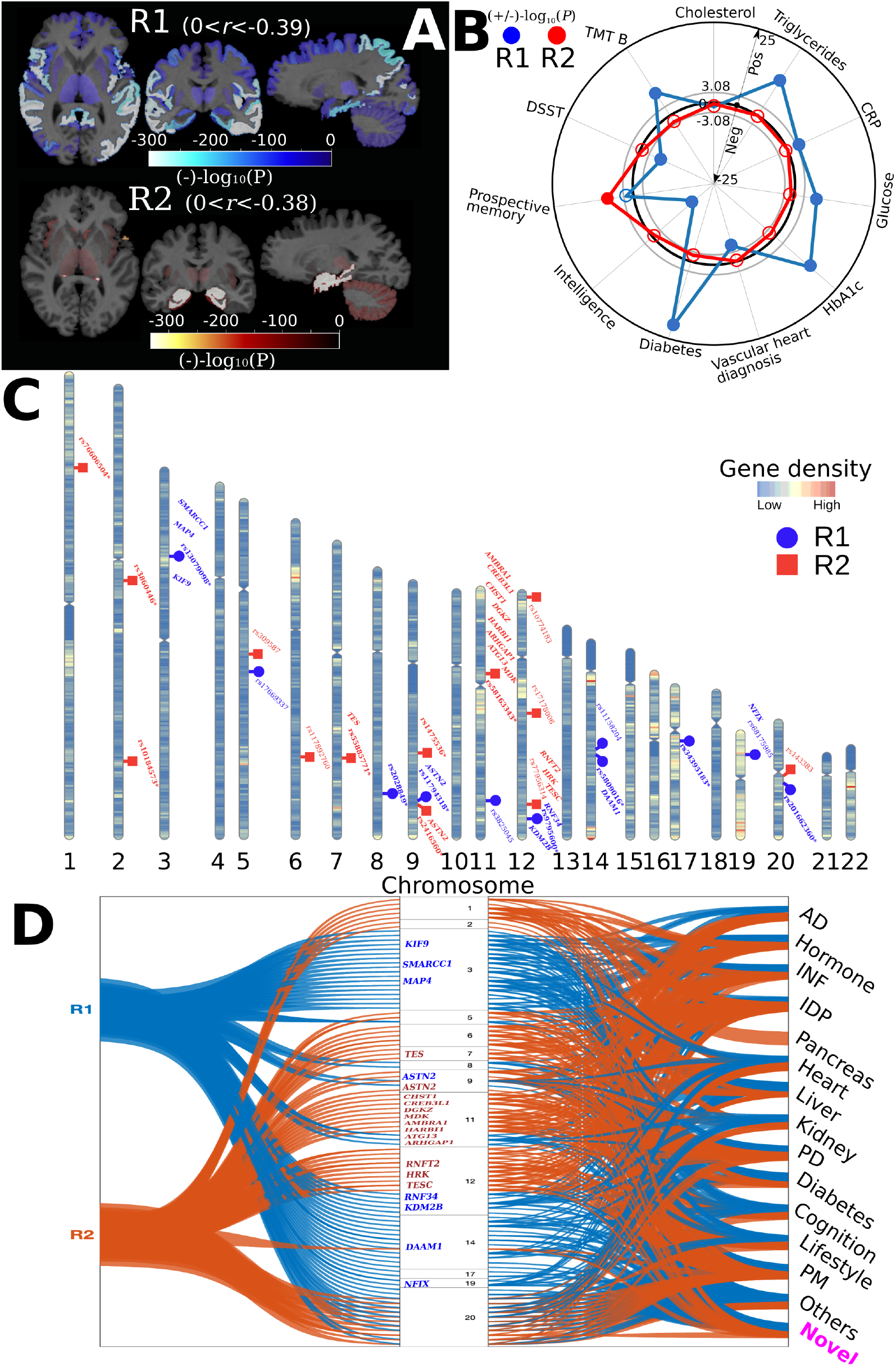
The expression of the R1 and R2 dimensions in the general population. **A)** Brain association studies confirm the presence of the two dimensions in the general population: the R1 dimension shows widespread brain atrophy, whereas the R2 dimension displays focal medial temporal lobe atrophy. P-value and effect sizes (*r*, Pearson’s correlation coefficient) are presented in **Supplementary eTable 4**. The range of *r* for each dimension is also shown. Of note, the sample size (*N*) for R1 and R2 is the same for each ROI. **B)** Clinical association studies further show that the R2 dimension is associated with prospective memory, and the R1 dimension is associated with several cognitive dysfunctions, cardiovascular risk factors (e.g., triglycerides), and diabetes (e.g., HbA1c). The same linear regression models were used to associate the R1 and R2 dimensions with the 61 clinical variables, including cardiovascular factors, diabetic blood markers, social demographics, lifestyle, physical measures, cognitive scores, and imaging-derived phenotypes. The radar plot shows representative clinical variables; all other results are presented in **Supplementary eTable 5**. The gray circle lines indicate the P-value threshold in both directions (Bonferroni correction for the 61 variables: –log_10_(P-value) > 3.08). A positive/negative –log_10_(P-value) value indicates a positive/negative correlation (beta). Transparent dots represent the associations that do not pass the Bonferroni correction; the blue-colored dots and red-colored dots indicate significant associations for the R1 and R2 dimensions, respectively. **C)** Genome-wide association studies demonstrate that the R2 dimension is associated to a larger extent with genomic loci and genes previously associated with AD-related traits in the literature (genome-wide P-value threshold with the red line: –log_10_(P-value) > 7.30). Each genomic locus is represented by its top lead SNP. The R1 dimension identified 8 (blue-colored in bold) out of the 49 mapped genes associated with AD-related traits. The R2 dimension identified 13 (red-colored in bold) out of 40 mapped genes associated with AD-related traits. Gene annotations were performed via positional, expression quantitative trait loci, and chromatin interaction mappings using FUMA (**Supplementary eTable 6** for all mapped genes).^53^ The genomic loci and mapped genes were manually queried in the GWAS Catalog^50^ to determine whether they were previously associated with AD (novel or not). * denotes that the genomic locus is novel. **D)** Besides AD-related traits, the genes and genomic loci in the two dimensions were also associated with other clinical traits, including inflammation, neurohormones, and imaging-derived phenotypes, in the literature from the GWAS Catalog.^50^ The flowchart first maps the genomic loci and genes (left) identified in the two dimensions onto the human genome (middle). It then links these variants to any clinical traits identified in previous literature from the GWAS Catalog (right). In the middle of the human genome, we show chromosomes 1 to 22 (above to below); the blue and red-colored genes are AD-related for the R1 and R2 dimensions, respectively. The black-colored genes (Fig. C) are not annotated. INF: inflammation; PD: psychiatric disorder; PM: physical measure; Novel (pink-colored in bold, corresponding to the novel loci/genes in Fig C) indicates that the locus or gene was not associated with any traits in the literature.

### Twenty-four genomic loci and seventy-seven genes unrelated to *APOE* are associated with the R1 and R2 dimensions in the general population

GWAS identified 24 genomic loci, 14 of which are novel (not previously associated with any traits in GWAS Catalog), and 77 positionally and functionally mapped genes unrelated to *APOE* associated with R1 or R2. In particular, the R1 dimension was significantly associated with 11 genomic loci and 49 genes. Eight genes (blue-colored genes in **Fig. 2D**) were previously associated with AD-related traits; 12 novel loci/genes have not been previously associated with any clinical traits. The R2 dimension was significantly associated with 13 genomic loci and 40 annotated genes. 13 genes (red-colored genes in **Fig. 2D**) were associated with AD-related traits; 8 loci/genes were novel (**Fig. 2C**, **Supplementary eTable 6**). These genomic loci and genes were also associated with many clinical traits in the literature from the GWAS Catalog.^50^ These included hormones (e.g., sex hormone-binding globulin measurement vs. *CCKN2C*), inflammatory factors (e.g., macrophage inflammatory protein 1b measurement vs. *CDC25A*), imaging-derived phenotypes (e.g., cerebellar volume measurement from MRIs vs. *DMRTA2*), and psychiatric disorders (e.g., unipolar depression vs. *ASTN2*) (**Fig. 2D**). Details of the GWAS Catalog results are presented in **Supplementary eFile 1**. The Manhatton and QQ plots of the baseline GWAS are presented in **Supplementary eFigure 8**. The LDSC^51^ interpcept of the two GWASs was close to 1, indicating no substantial genomic inflation (R1=1.0032±0.0084; R2=1.023±0.0084). Furthermore, our main GWASs using European ancestry were robust in three sensitivity check analyses: split-sample, sex-stratified, and mixed-effect^52^ linear model analyses. Detailed results are presented in **Supplementary eText 1** and **Supplementary eFile2-4**.

The two dimensions were significantly heritable in the general population based on the SNP-based heritability estimates (R1: *h*^2^ = 0.49 ± 0.02; R2: *h*^2^ = 0.55 ± 0.02). The PRS of AD showed a marginally positive association with the R2 dimension [−log_10_(P-value)=1.42], but not with the R1 dimension [−log_10_(P-value)=0.47<1.31] in this population.

### Genes associated with the R1 and R2 dimensions are overrepresented in organs beyond the brain in the general population

Tissue specificity analyses test whether the mapped genes are overrepresented in differentially expressed gene sets (DEG) in one organ/tissue compared to all other organs/tissues using different gene expression data^53^. The genes associated with the R1 dimension were overrepresented in caudate, hippocampus, putamen, amygdala, substantia nigra, liver, heart, and pancreas; the genes associated with the R2 dimension were overrepresented in caudate, hippocampus, putamen, amygdala, anterior cingulate, pituitary, liver, muscle, kidney, and pancreas (**Fig. 3A** and **Supplementary eFigure 9**). Genes in DEG over-expressed in the heart were only associated with R1, while those in DEG over-expressed in the pituitary gland, muscle, and kidney were unique in R2. The expression values of every single gene for all tissues are presented in **Supplementary eFigure 10**.

**Figure 3:**
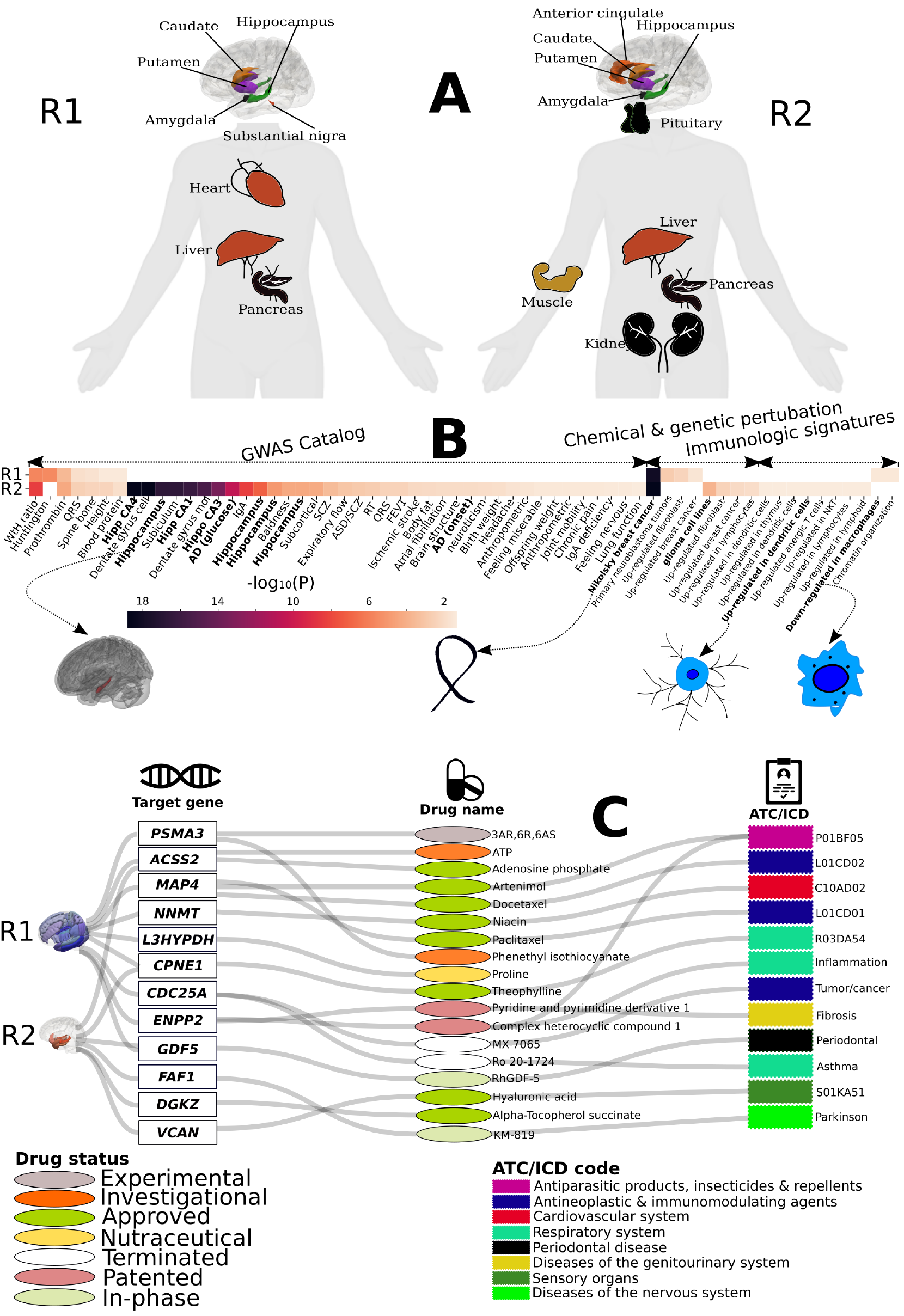
Tissue specificity and biological pathway enrichment analysis of the R1 and R2 dimensions in the general population. **A)** Tissue specificity analyses show that genes associated with the two dimensions of neurodegeneration are overrepresented in organs/tissues beyond the human brain (R1 and R2). The unique overrepresentation of genes in differentially expressed gene sets (DEG) in the heart (R1) and the pituitary gland, muscle, and kidney (R2) may imply the involvement of inflammation^12,71,72^ and neurohormone dysfunction^15–17^, respectively. The *GENE2FUNC*^53^ pipeline from FUMA was performed to examine the overrepresentation of prioritized genes (Fig. 2C) in pre-defined DEGs (up-regulated, down-regulated, and both-side DEGs) from different gene expression data. The input genes (Fig. 2C) were tested against each DEG using the hypergeometric test. We present only the organs/tissues that passed the Bonferroni correction for multiple comparisons. **B)** Gene set enrichment analysis shows that genes associated with the two dimensions are enriched in different biological pathways. For example, genes associated with the R1 dimension are implicated in down-regulated macrophage functions, which have been shown to be associated with inflammation.^13^ In contrast, the R2 dimension is enriched in AD hallmarks (e.g., hippocampus), AD-related gene sets, and the pathway involved in dendritic cells, which may regulate amyloid-β-specific T-cell entry into the brain.^55^ Both dimensions are enriched in gene sets involved in cancer, which may indicate overlapped genetic underpinnings between AD and cancer.^54^ The *GENE2FUNC*^53^ pipeline from FUMA was performed to examine the enrichment of prioritized genes (Fig. 2C) in pre-defined gene sets. Hypergeometric tests were performed to test whether the input genes were overrepresented in any pre-defined gene sets. Gene sets were obtained from different sources, including MsigDB^93^ and GWAS Catalog.^50^ We show the significant results from gene sets defined in the GWAS Catalog, curated gene sets, and immunologic signature gene sets. All results are shown in **Supplementary eTable 7. C)** The target-drug-disease network for R1 and R2-associated genes provides great potential for drug discovery and repurposing. R1-annotated “druggable genes” were developed for cardiovascular diseases, various cancers, and inflammation, whereas R2-annotated “druggable genes” were developed for diseases of the nervous system (e.g., Parkinson’s disease). For the target-drug-disease network, the 5^th^ level of the Anatomical Therapeutic Chemical (ATC) code is displayed for the DrugBank database,^94^ and the disease name defined by the International Classification of Diseases (ICD-11) code is showed for the Therapeutic Target Database.^95^

### Genes associated with the R1 and R2 dimensions are enriched in key biological pathways in the general population

Genes associated with the two dimensions were enriched in different biological pathways. Genes associated with the two dimensions were implicated in several types of cancer, including up-regulation of fibroblast, breast cancer, and neuroblastoma tumors (**Fig. 3B**), which indicate a certain extent of genetic overlaps and shared pathways that may explain the intriguing inverse relationship between AD and cancer.^54^ Genes associated with the R1 dimension were implicated in pathways involved in the down-regulation of macrophages (**Fig. 3B**), which are involved in the initiation and progression of various inflammatory processes, including neuroinflammation and AD.^13^ Inflammation is also known to be associated with vascular compromise and dysfunction. This further concurs with the stronger cardiovascular profile of R1, especially with increased WML and predominant SPARE-BA increases. Genes associated with the R2 dimensions were enriched in pathways involved in AD onset, hippocampus-related brain volumes, and dendritic cells (**Fig. 3B**). In particular, dendritic cells may regulate amyloid-β-specific T-cell entry into the brain,^55^ as well as the inflammatory status of the brain.^56^ The gene set enrichment analysis results are presented in **Supplementary eTable 7**.

### Genes associated with the R1 and R2 dimensions show potential for drug discovery and repurposing

We queried whether these 77 genes associated with R1 and R2 are “druggable genes” from the constructed target-drug-disease network – the target genes express proteins to bind drug-like molecules, and the drug is at any stage of the clinical trial. For the 49 R1-annotated genes, 9 genes were targets for 15 drugs and drug-like molecules, treating various cancer, inflammation, and cardiovascular dysfunctions. For the 40 R2-annotated genes, 6 genes were targets for 7 drugs developed for diseases of the nervous system, such as Parkinson’s (**Fig. 3C**). The pharmacological mechanisms targeted by these identified drugs are largely related to the pathogenesis of AD in previous literature. For example, FDA-approved Niacin [R1; target gene: *NNMT*; Anatomical Therapeutic Chemical (ATC) code: C10AD02] is a B vitamin used to treat various deficiencies and diseases in the cardiovascular system, including myocardial infarctions,^57^ hyperlipidemia,^58^ and coronary artery disease.^59^ Interestingly, a recent study^60^ showed that Niacin detained AD progression in a 5xFAD mice model. The niacin receptor HCAR2 modulates microglial response to amyloid deposition, ultimately alleviating neuronal loss and cognitive decline. Other drugs for potential drug repurposing of AD are the FDA-approved Docetaxel (R1; target gene: *MAP4*; ATC: L01CD02) and Paclitaxel (R1; target gene: *MAP4*; ATC: L01CD01), which both target various cancers, including breast cancer and metastatic prostate cancer. The intriguing inverse relationship between AD and cancer has long been established, but the underlying shared etiology remains unclear.^54,61^ One hypothesis was that microtubule-associated protein tau – a pathological biomarker of AD – was associated with resistance to Docetaxel in certain cancer treatments.^62^ In addition, Docetaxel impacted the blood-brain barrier function of breast cancer brain metastases.^63^ Another drug called KM-819 (R2; target gene: *FAF1*) is currently in Phase 1 for a clinical trial of Parkinson’s disease,^64^ which aims to suppress α-synuclein-induced mitochondrial dysfunction,^65^ consistent with the mitochondrial hypothesis^66^ of AD. To sum up, R1 and R2 show distinct landscapes of the “druggable genome”^67^ on drug discovery and repurposing^68^ for future clinical translation.

### The longitudinal rate of change in the R2 dimension, but not R1, is marginally associated with the *APOE ε4* allele, tau in cognitively unimpaired individuals

Using cognitively unimpaired participants from ADNI and BLSA, longitudinal brain association studies showed that the rate of change in the R1 dimension was associated with the change of brain volume in widespread brain regions. In contrast, the rate of change in the R2 dimension was associated with the change of brain volume in the focal medial temporal lobe (**Fig. 4A** and **Supplementary eTable 8** for P-values and effect sizes). This further indicates that the two dominant patterns discovered cross-sectionally also progress in consistent directions longitudinally. The two dimensions were not associated with CSF biomarkers (Aβ42, tau, and p-tau) and the *APOE ε4* allele (*rs429358*) at baseline [–log_10_(P-value) < 1.31)]. The rate of change of the R2 dimension, but not R1, was marginally [nominal threshold: –log_10_(P-value) > 1.31] associated with the *APOE ε4* allele, the CSF level of tau, and p-tau (**Fig. 4B** and **Supplementary eTable 9** for P-values and effect sizes), but they did not survive the Bonferroni correction [–log_10_(P-value) = 2.95]. The longitudinal rate of change of both dimensions was negatively associated [–log_10_(P-value) > 2.95] with the total CSF level of Aβ42.

**Figure 4:**
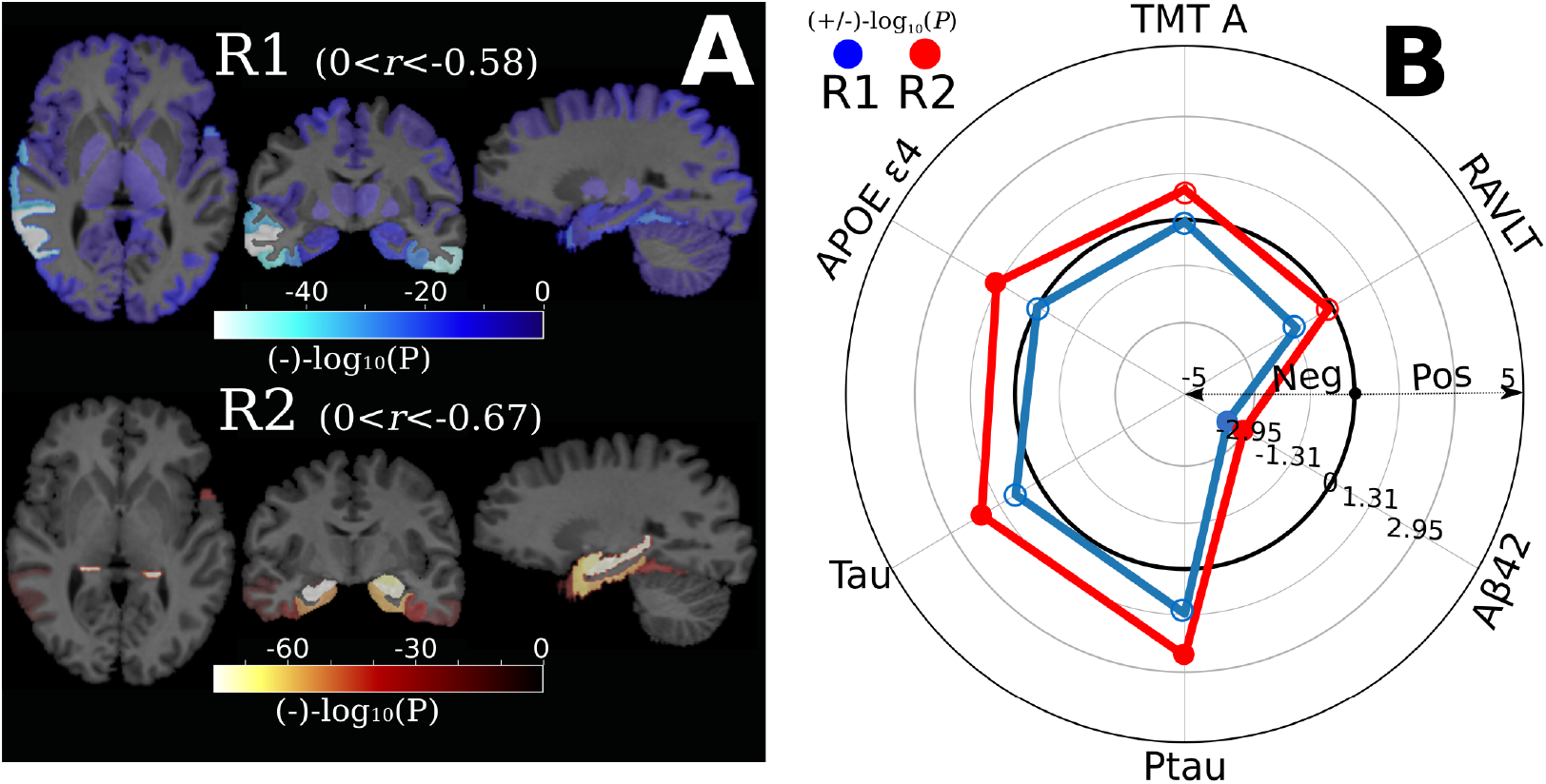
The longitudinal rate of change in R1 and R2 in the cognitively unimpaired population. **A)** Longitudinal brain association studies show that the R1 dimension exhibits longitudinal brain volume decrease in widespread brain regions, whereas the R2 dimension displays longitudinal brain volume decrease in the focal medial temporal lobe. We first derived the rate of change of the 119 GM ROIs and the R1 and R2 dimensions using a linear mixed effect model; a linear regression model was then fit to the rate of change of the ROIs, R1, and R2 to derive the beta coefficient value of each ROI. A negative value denotes longitudinal brain changes with a negative coefficient of the rate of change in the linear regression model. P-value and effect sizes (*r*, Pearson’s correlation coefficient) are presented in **Supplementary eTable 8**. The range of *r* for each dimension is also shown. Of note, the sample size (*N*) for R1 and R2 is the same for each ROI. **B)** The rate of change, not the baseline measurement, in the two dimensions is negatively associated with the CSF level of Aβ42 (Bonferroni correction for the 45 variables: –log_10_(P-value) > 2.95). The rate of change in the R2 dimension, not the R1 dimension, was marginally (–log_10_(P-value) > 1.31) associated with the CSF level of tau and p-tau, and *APOE ε4*. All other clinical associations are presented in **Supplementary eTable 9**. The gray-colored circle lines indicate different P-value thresholds in both directions (Bonferroni correction for the 45 variables: –log_10_(P-value) > 2.95 and the nominal P-value threshold: –log_10_(P-value) > 1.31). A positive/negative –log_10_(P-value) value indicates a positive/negative correlation (beta). Transparent dots represent the associations that do not pass the nominal P-value threshold [log_10_(P-value) = 1.31]; the blue-colored dots and red-colored dots indicate significant associations [log_10_(P-value) > 1.31] for the R1 and R2 dimensions, respectively.

We tested these associations using cognitively unimpaired individuals with a high risk of AD based on their family history from the PREVENT-AD cohort. Similarly, at baseline, the two dimensions were not associated with CSF biomarkers or the *APOE ε4* allele (*rs429358*). The longitudinal rate of change in the R2 dimension, but not R1, was marginally [nominal threshold: –log_10_(P-value) > 1.31] associated with the *APOE ε4* allele [–log_10_(P-value) = 1.92], the CSF level of tau [–log_10_(P-value) = 1.65], and p-tau [–log_10_(P-value) = 1.66].

Longitudinal brain association studies also confirmed the longitudinal progression of the two dimensions in the MCI/AD population (**Supplementary eFigure 11A**). The rates of change in the two dimensions were both associated with *APOE ε4* [–log_10_(P-value) = 12.54 for R1 and 9.05 for R2] in GWAS (**Supplementary eFigure 11B**), and related to CSF levels of tau [–log_10_(P-value) = 16.47 for R1 and 9.73 for R2], p-tau [–log_10_(P-value) = 19.13 for R1 and 10.81 for R2], and Aβ42 [–log_10_(P-value) = 13.64 for R1 and 13.55 for R2] (**Supplementary eFigure 11C).**

## Discussion

The current study leveraged a deep semi-supervised representation learning method to establish two predominant dimensions in the symptomatic MCI/AD population, which were independently found to be expressed, to a lesser degree, in three asymptomatic populations. In particular, the R1 dimension represented a “diffuse-AD” atrophy pattern: varying degrees of brain atrophy throughout the entire brain. In contrast, the R2 dimension showed an “MTL-AD” atrophy pattern: brain atrophy predominantly concentrated in the medial temporal lobe (**Fig. 1A**). Importantly, only R2 was found to be significantly associated with genetic variants of the *APOE* genes in MCI/AD patients. Furthermore, our study examined early manifestations of the R1 and R2 dimensions in asymptomatic populations with varying levels of AD risks and their associations with genetics, amyloid plaques and tau tangles, biological pathways, and body organs. We identified that 24 genomic loci, 14 of which are novel, and 77 annotated genes contribute to early manifestations of the two dimensions. Functional analyses showed that genes unrelated to *APOE* were overrepresented in DEG sets in organs beyond the brain (R1 and R2), including the heart (R1) and the pituitary gland (R2), and enriched in several biological pathways involved in dendritic cells (R2), macrophage functions (R1), and cancer (R1 and R2). Several of these genes were “druggable genes” for cancer (R1), inflammation (R1), cardiovascular diseases (R1), and diseases of the nervous system (R2). Longitudinal findings in the cognitively unimpaired populations showed that the rate of change of the R2 dimension, but not R1, was marginally associated with the *APOE ε4* allele, the CSF level of tau, and Aβ42 (R1 and R2). Our findings suggested that diverse pathologic processes, including cardiovascular risk factors, neurohormone dysfunction, and inflammation, might occur in the early asymptomatic stages, supporting and expanding the current amyloid cascade.^7,8^

AD has been regarded as a CNS disorder. However, increasing evidence has indicated that the origins or facilitators of the pathogenesis of AD might involve processes outside the brain.^6^ For example, recent findings revealed that gut microbiota disturbances might influence the brain through the immune and endocrine system and the bacteria-derived metabolites.^69,70^ Our findings support the view that multiple pathological processes might contribute to early AD pathogenesis and identify non-*APOE* genes in the two dimensions overrepresented in tissues beyond the brain (e.g., the heart, pituitary gland, muscle, and kidney). Pathological processes may be involved in different cells, molecular functions, and biological pathways, exaggerating amyloid plaque and tau tangle accumulation and leading to the downstream manifestation of neurodegeneration and cognitive decline.

The genetic and clinical underpinnings of the R1 dimension support inflammation, as well as cardiovascular diseases, as a core pathology contributing to AD.^12,71,72^ Genes associated with the R1 dimension were previously associated with various inflammation-related clinical traits (**Fig. 2D**), and enriched in biological pathways involved in immunological response (e.g., up-regulation in macrophages^73^, **Fig. 3B**). In addition, genes in this dimension were overrepresented in DEG sets in the heart (**Fig. 3A**). Previous literature indicated that inflammation is likely an early step that initiates the amyloidogenic pathway – the expression of inflammatory cytokines leads to the production of β-amyloid plaques.^13^ Several markers of inflammation are also present in serum and CSF before any indications of Aβ or tau tangles.^74^ For example, clusterin, a glycoprotein involved in many processes and conditions (e.g., inflammation, proliferation, and AD) induced by tumor necrosis factor (TNF), was present ten years earlier than Aβ deposition.^75^ In addition, the R1 dimension was also strongly associated with cardiovascular and diabetes biomarkers (**Fig. 2B**). Inflammatory processes have been critical, well-established risk factors for compromised cardiovascular function,^76^ such as coronary artery disease and the breakdown of the blood-brain barrier. Our results corroborated the close relationships between AD, cardiovascular diseases, and inflammation.

The genetic and clinical underpinnings of the R2 dimension support that neuroendocrine dysfunction might be an early event contributing to the pathogenesis of AD.^16,17^ Genes in the R2 dimension were previously associated with different hormone and pancreas-related traits from GWAS Catalog (**Fig. 2D**); they were also overrepresented in DEG in the pituitary and pancreas glands, muscle and kidney (**Fig. 3A**), which are master glands or key organs in the endocrine system.^77^ Previous literature suggested that neuroendocrine dysfunction might contribute to AD development by secreting neurohormonal analogs and affecting CNS function.^16^ For example, luteinizing hormone-releasing hormone and follicle-stimulating hormone in serum or neurons were associated with the accumulation of Aβ plaques in the brain.^17,78^.^79^ However, early experimental studies on antagonists of Luteinizing hormone-releasing hormone and growth hormone-releasing hormone in animal models of AD have shown promising but not entirely convincing evidence.^16^ Taken together, neurodegeneration in the R2 dimension represents an AD-specific phenotype that might be driven by hormonal dysfunction, leading to rapid accumulation of amyloid plaques, and was potentially accelerated by the *APOE ε4* allele – the rate of change in R2, but not R1, was associated with the *APOE ε4* allele in cognitively unimpaired individuals (**Fig. 4B**).

The hypothesized implications above of the R1 and R2 dimensions on inflammation, cardiovascular functions, and neuroendocrine dysfunctions are not mutually exclusive and may collectively contribute to AD pathogenesis. It has been shown that dysregulation of the hypothalamic-pituitary-gonadal axis is associated with dyotic signaling, modulating the expression of TNF and related cytokines in systemic inflammation, and the induction of downstream neurodegenerative cascades within the brain.^80,81^ These studies hypothesized that the neuroendocrine dysfunction and the inflammation mechanism might be the upstream and downstream neuropathological processes along the disease course of AD.^16^ That is, the loss of sex steroids and the elevation of gonadotropins might lead to a higher level of inflammatory factors in the brain. Finally, other competing hypotheses may also play a role in developing AD in early asymptomatic stages, including the mitochondrial hypothesis,^66^ the metabolic hypothesis,^82^ and the tau hypothesis.^3^

The NIA-AA framework^83^ claims that AD is a continuum in which AD pathogenesis is initiated in early asymptomatic cognitively unimpaired stages and progresses to amyloid-positive and tau-positive (A+T+) in late symptomatic stages.^83^ Our findings are consistent with this framework and elucidate the cross-sectional and longitudinal associations of the two dimensions with genetic and clinical markers from early asymptomatic to late symptomatic stages. In early asymptomatic stages, the rates of change in the two dimensions are both associated with amyloid. However, only the R2 dimension, not R1, is marginally associated with the *APOE ε4* allele and the CSF level of tau (**Fig. 4B**). In contrast, in late symptomatic stages, the rates of change in the two dimensions are both associated with the *APOE ε4* allele, CSF levels of tau, p-tau, and amyloid (**Supplementary eFigure 11**). Our findings suggest that comorbidities or normal aging in R1 may alter the rate or trajectory of neurodegeneration at early asymptomatic stages, but *APOE*-related genes might play a more pronounced role in the acceleration and progression during late symptomatic stages for both dimensions (**Fig. 5**).

**Figure 5:**
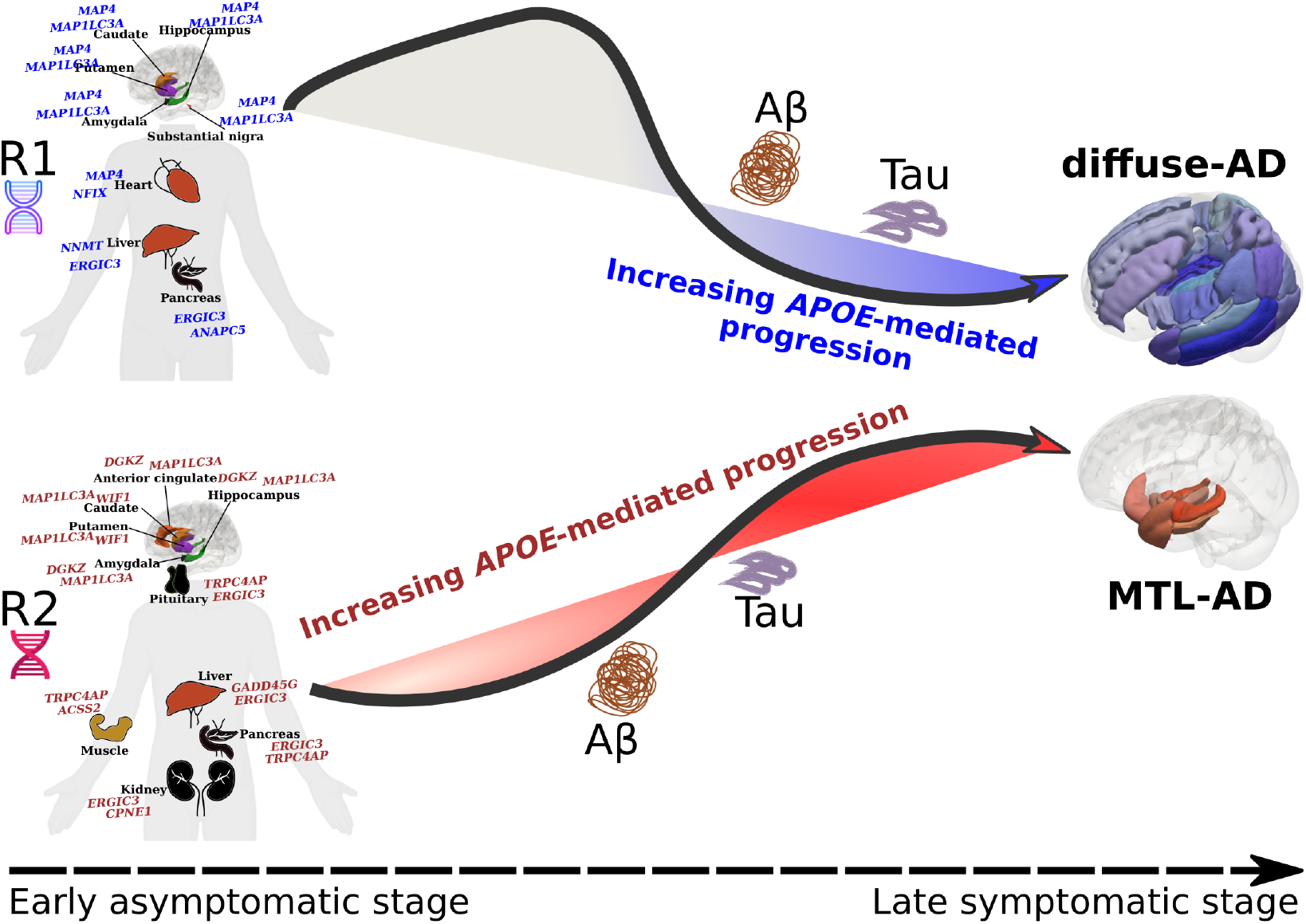
Genes unrelated to *APOE* influence early manifestations of R1 and R2. Genes unrelated to *APOE* and overrepresented in organs beyond the human brain are associated with early manifestations of the R1 (diffuse-AD) and R2 (MTL-AD) dimensions, which capture the heterogeneity of AD-related brain atrophy. For visualization purposes, we display the two genes with the highest expression values in the tissue specificity analyses for each organ/tissue. The black arrow line emulates the longitudinal progression trajectory along these two dimensions. The positions of beta-amyloid, tau, and *APOE* (increasing *APOE*-mediated progression) indicate the time point when they are associated with the two dimensions. The blue/red gradient-color background indicates a higher influence of *APOE*-related genes (left to right; early to late stages). The brain atrophy patterns are presented in the 3D view. In early asymptomatic stages, the R1-related genes are implicated in cardiovascular diseases and inflammation; the R2-related genes are involved in hormone-related dysfunction. Critically, longitudinal progression of the dimension demonstrates an impact of the *APOE* genes in early asymptomatic stages in R2, but this longitudinal effect occurs only in late symptomatic stages in R1. These results suggest that comorbidities (e.g., cardiovascular conditions) or normal aging in R1 may alter or delay the trajectory of neurodegeneration in early asymptomatic stages; *APOE*-related genes may play a pronounced role in the acceleration and progression in late symptomatic stages for both dimensions. Of note, the underlying pathological processes that initiate and drive the progression of the two dimensions are not mutually exclusive. Hence, both R1 and R2 can be co-expressed in the same individual. In addition, the two dimensions can also be affected by other AD hypotheses, such as the mitochondrial hypothesis^66^ and the metabolic hypothesis.^82^ MTL: medial temporal lobe.

Several recent studies^84–88^, as detailed in an insightful overview by Luo et al., collectively provide a comprehensive transcriptomics and epigenomics atlas depicting AD progression at the single-cell level^89^. These studies highlight the involvement of microglia-related inflammation, lipid metabolism, and mitochondrial dysfunction. This substantiates the primary hypothesis in our study: the two dimensions are linked to diverse pathological mechanisms, encompassing cardiovascular diseases, inflammation, and hormonal dysfunction, potentially driven by genes beyond *APOE*.

## Limitations

This study has several limitations. Firstly, there is a need for longitudinal data from the general population, as exemplified by the UK Biobank, to provide further validation for the hypotheses proposed to cover the entire AD spectrum in the same population. Secondly, it is essential to extend the generalization of the current GWAS findings to include underrepresented ethnic groups, going beyond the European ancestry populations. Lastly, substantial methodological advancements are imperative, particularly in the integration of imaging and genetic data, to effectively capture the full spectrum of AD heterogeneity.

## Outlook

In conclusion, the current study used a novel deep semi-supervised representation learning method to establish two AD dimensions. Our findings support that those diverse pathological mechanisms, including cardiovascular diseases, inflammation, hormonal dysfunction, and involving multiple organs,^90–92^ collectively affect AD pathogenesis in asymptomatic stages. These novel biomarkers may serve as instrumental variables to guide future treatments in the early asymptomatic stages of AD, targeting multi-organ dysfunction beyond the brain.

## Data availability

The GWAS summary statistics corresponding to this study are publicly available on the MEDICINE knowledge portal (http://labs.loni.usc.edu/medicine/) and the BRIDGEPORT web portal (https://www.cbica.upenn.edu/bridgeport/).

## Acknowledgements

This research has been conducted using the UK Biobank Resource under Application Number 35148. Data used in the preparation of this article were in part obtained from the Alzheimer’s Disease Neuroimaging Initiative (ADNI) database (adni.loni.usc.edu). As such, the investigators within the ADNI contributed to the design and implementation of ADNI and/or provided data but did not participate in the analysis or writing of this report. A complete listing of ADNI investigators can be found at: http://adni.loni.usc.edu/wpcontent/uploads/how_to_apply/ADNI_Acknowledgement_List.pdf. ADNI is funded by the National Institute on Aging, the National Institute of Biomedical Imaging and Bioengineering, and through generous contributions from the following: AbbVie, Alzheimer’s Association; Alzheimer’s Drug Discovery Foundation; Araclon Biotech; BioClinica, Inc.; Biogen; Bristol-Myers Squibb Company; CereSpir, Inc.; Cogstate; Eisai Inc.; Elan Pharmaceuticals, Inc.; Eli Lilly and Company; EuroImmun; F. Hoffmann-La Roche Ltd and its affiliated company Genentech, Inc.; Fujirebio; GE Healthcare; IXICO Ltd.; Janssen Alzheimer Immunotherapy Research & Development, LLC.; Johnson & Johnson Pharmaceutical Research & Development LLC.; Lumosity; Lundbeck; Merck & Co., Inc.; Meso Scale Diagnostics, LLC.; NeuroRx Research; Neurotrack Technologies; Novartis Pharmaceuticals Corporation; Pfizer Inc.; Piramal Imaging; Servier; Takeda Pharmaceutical Company; and Transition Therapeutics. The Canadian Institutes of Health Research is providing funds to support ADNI clinical sites in Canada. Private sector contributions are facilitated by the Foundation for the National Institutes of Health (www.fnih.org). The grantee organization is the Northern California Institute for Research and Education, and the study is coordinated by the Alzheimer’s Therapeutic Research Institute at the University of Southern California. ADNI data are disseminated by the Laboratory for Neuro Imaging at the University of Southern California.

## Funding

The primary funding support for this present study is from the initial funding package as an Assistant Professor of Neurology for WJ, provided by Stevens Neuroimaging and Informatics Institute, Keck School of Medicine of USC, University of Southern California. The iSTAGING consortium is a multi-institutional effort funded by NIA by RF1 AG054409 for DC. The Baltimore Longitudinal Study of Aging neuroimaging study is funded by the Intramural Research Program, National Institute on Aging, National Institutes of Health and by HHSN271201600059C. Other supporting grants include 5U01AG068057-02, 1U24AG074855-01. SRH is Supported by multiple grants and contracts from NIH.

## Authors’ contributions

Dr. Wen has full access to all the data in the study and takes responsibility for the integrity of the data and the accuracy of the data analysis.

*Study concept and design*: Wen, Davatzikos

*Acquisition, analysis, or interpretation of data*: Wen, Davatzikos

*Drafting of the manuscript*: Wen

*Critical revision of the manuscript for important intellectual content*: all authors

*Statistical and genetic analysis*: Wen

## Competing interests

DAW served as Site PI for studies by Biogen, Merck, and Eli Lilly/Avid. He has received consulting fees from GE Healthcare and Neuronix. He is on the DSMB for a trial sponsored by Functional Neuromodulation. AJS receives support from NIH (P30 AG010133, P30 AG072976, R01 AG019771, R01 AG057739, U01 AG024904, R01 LM013463, R01 AG068193, T32 AG071444, and U01 AG068057 and U01 AG072177). He has also received support from Avid Radiopharmaceuticals, a subsidiary of Eli Lilly (in-kind contribution of PET tracer precursor); Bayer Oncology (Scientific Advisory Board); Eisai (Scientific Advisory Board); Siemens Medical Solutions USA, Inc. (Dementia Advisory Board); Springer-Nature Publishing (Editorial Office Support as Editor-in-Chief, Brain Imaging, and Behavior). ME receives support from multiple NIH grants, the Alzheimer’s Association, and the Alzheimer’s Therapeutic Research Institute. HJG has received travel grants and speaker honoraria from Fresenius Medical Care, Neuraxpharm, Servier, and Janssen Cilag and research funding from Fresenius Medical Care. HJG had personal contracts approved by the university administration for speaker honoraria and one IIT with Fresenius Medical Care.

## Supplementary material

Supplementary material is available at *Brain* online.

## Notes

### Summary of Updates

We had a revision at a journal under peer review. This is the revised version. Thanks

## References

1. Guthrie, H. et al. Safety, Tolerability, and Pharmacokinetics of Crenezumab in Patients with Mild-to-Moderate Alzheimer’s Disease Treated with Escalating Doses for up to 133 Weeks. J Alzheimers Dis 76, 967–979 (2020).

2. Sevigny, J. et al. The antibody aducanumab reduces Aβ plaques in Alzheimer’s disease. Nature 537, 50–56 (2016).

3. Congdon, E. E. & Sigurdsson, E. M. Tau-targeting therapies for Alzheimer disease. Nat Rev Neurol 14, 399–415 (2018).

4. Hardy, J. & Selkoe, D. J. The Amyloid Hypothesis of Alzheimer’s Disease: Progress and Problems on the Road to Therapeutics. Science 297, 353–356 (2002).

5. Brinkmalm, G. & Zetterberg, H. The phosphorylation cascade hypothesis of Alzheimer’s disease. Nat Aging 1, 498–499 (2021).

6. Du, X., Wang, X. & Geng, M. Alzheimer’s disease hypothesis and related therapies. Translational Neurodegeneration 7, 2 (2018).

7. Jack, C. R. et al. Tracking pathophysiological processes in Alzheimer’s disease: an updated hypothetical model of dynamic biomarkers. Lancet Neurol 12, 207–216 (2013).

8. Frisoni, G. B. et al. The probabilistic model of Alzheimer disease: the amyloid hypothesis revised. Nat Rev Neurosci 23, 53–66 (2022).

9. Makin, S. The amyloid hypothesis on trial. Nature 559, S4–S7 (2018).

10. Herrup, K. The case for rejecting the amyloid cascade hypothesis. Nat Neurosci 18, 794– 799 (2015).

11. Barisano, G. et al. Blood–brain barrier link to human cognitive impairment and Alzheimer’s disease. Nat Cardiovasc Res 1, 108–115 (2022).

12. Wyss-Coray, T. Inflammation in Alzheimer disease: driving force, bystander or beneficial response? Nat Med 12, 1005–1015 (2006).

13. Heneka, M. T. et al. Neuroinflammation in Alzheimer’s Disease. Lancet Neurol 14, 388– 405 (2015).

14. Leng, F. & Edison, P. Neuroinflammation and microglial activation in Alzheimer disease: where do we go from here? Nat Rev Neurol 17, 157–172 (2021).

15. Dean, D. W. Neuroendocrine Theory of Aging: Chapter 1. (2012).

16. Schally, A. V. Endocrine approaches to treatment of Alzheimer’s disease and other neurological conditions: Part I: Some recollections of my association with Dr. Abba Kastin: A tale of successful collaboration. Peptides 72, 154–163 (2015).

17. Xiong, J. et al. FSH blockade improves cognition in mice with Alzheimer’s disease. Nature 603, 470–476 (2022).

18. Dubois, B. et al. Preclinical Alzheimer’s disease: Definition, natural history, and diagnostic criteria. Alzheimers Dement 12, 292–323 (2016).

19. Rajpurkar, P., Chen, E., Banerjee, O. & Topol, E. J. AI in health and medicine. Nat Med 28, 31–38 (2022).

20. Wen, J. et al. Convolutional neural networks for classification of Alzheimer’s disease: Overview and reproducible evaluation. Medical Image Analysis 63, 101694 (2020).

21. Kendler, K. & Neale, M. Endophenotype: a conceptual analysis. Mol Psychiatry 15, 789– 797 (2010).

22. Yang, Z. et al. A deep learning framework identifies dimensional representations of Alzheimer’s Disease from brain structure. Nat Commun 12, 7065 (2021).

23. Wen, J. et al. Multi-scale semi-supervised clustering of brain images: Deriving disease subtypes. Med Image Anal 75, 102304 (2021).

24. Young, A. L. et al. Uncovering the heterogeneity and temporal complexity of neurodegenerative diseases with Subtype and Stage Inference. Nat Commun 9, 4273 (2018).

25. Zhang, X. et al. Bayesian model reveals latent atrophy factors with dissociable cognitive trajectories in Alzheimer’s disease. COGNITIVE SCIENCES 10.

26. Vogel, J. W. et al. Four distinct trajectories of tau deposition identified in Alzheimer’s disease. Nat Med 27, 871–881 (2021).

27. Bellenguez, C. et al. New insights into the genetic etiology of Alzheimer’s disease and related dementias. Nat Genet 54, 412–436 (2022).

28. Lambert, J.-C. et al. Meta-analysis of 74,046 individuals identifies 11 new susceptibility loci for Alzheimer’s disease. Nat Genet 45, 1452–1458 (2013).

29. Yang, Z., Wen, J. & Davatzikos, C. Surreal-GAN:Semi-Supervised Representation Learning via GAN for uncovering heterogeneous disease-related imaging patterns. ICLR (2021).

30. Petersen, R. C. et al. Alzheimer’s Disease Neuroimaging Initiative (ADNI): clinical characterization. Neurology 74, 201–209 (2010).

31. Bycroft, C. et al. The UK Biobank resource with deep phenotyping and genomic data. Nature 562, 203–209 (2018).

32. Resnick, S. M. et al. One-year age changes in MRI brain volumes in older adults. Cereb Cortex 10, 464–472 (2000).

33. Breitner, J. C. S., Poirier, J., Etienne, P. E. & Leoutsakos, J. M. Rationale and Structure for a New Center for Studies on Prevention of Alzheimer’s Disease (StoP-AD). J Prev Alzheimers Dis 3, 236–242 (2016).

34. Habes, M. et al. The Brain Chart of Aging: Machine-learning analytics reveals links between brain aging, white matter disease, amyloid burden, and cognition in the iSTAGING consortium of 10,216 harmonized MR scans. Alzheimer’s & Dementia 17, 89–102 (2021).

35. Tustison, N. J. et al. N4ITK: improved N3 bias correction. IEEE Trans. Med. Imaging 29, 1310–1320 (2010).

36. Doshi, J. et al. MUSE: MUlti-atlas region Segmentation utilizing Ensembles of registration algorithms and parameters, and locally optimal atlas selection. Neuroimage 127, 186–195 (2016).

37. Pomponio, R. et al. Harmonization of large MRI datasets for the analysis of brain imaging patterns throughout the lifespan. Neuroimage 208, 116450 (2020).

38. Zhang, X. et al. Bayesian model reveals latent atrophy factors with dissociable cognitive trajectories in Alzheimer’s disease. Proc Natl Acad Sci USA 113, E6535–E6544 (2016).

39. Ferreira, D., Nordberg, A. & Westman, E. Biological subtypes of Alzheimer disease: A systematic review and meta-analysis. Neurology 94, 436–448 (2020).

40. Wen, J. et al. Subtyping Brain Diseases from Imaging Data. in Machine Learning for Brain Disorders (ed. Colliot, O.) 491–510 (Springer US, 2023). doi:10.1007/978-1-0716-3195-9_16.

41. Yang, J., Lee, S. H., Wray, N. R., Goddard, M. E. & Visscher, P. M. GCTA-GREML accounts for linkage disequilibrium when estimating genetic variance from genome-wide SNPs. PNAS 113, E4579–E4580 (2016).

42. Zhao, B. et al. Genome-wide association analysis of 19,629 individuals identifies variants influencing regional brain volumes and refines their genetic co-architecture with cognitive and mental health traits. Nat Genet 51, 1637–1644 (2019).

43. Choi, S. W., Mak, T. S.-H. & O’Reilly, P. F. Tutorial: a guide to performing polygenic risk score analyses. Nat Protoc 15, 2759–2772 (2020).

44. Davatzikos, C., Xu, F., An, Y., Fan, Y. & Resnick, S. M. Longitudinal progression of Alzheimer’s-like patterns of atrophy in normal older adults: the SPARE-AD index. Brain 132, 2026–2035 (2009).

45. Moradi, E., Hallikainen, I., Hänninen, T. & Tohka, J. Rey’s Auditory Verbal Learning Test scores can be predicted from whole brain MRI in Alzheimer’s disease. NeuroImage: Clinical 13, 415–427 (2017).

46. Squire, L. R., Stark, C. E. L. & Clark, R. E. The medial temporal lobe. Annu Rev Neurosci 27, 279–306 (2004).

47. Bashyam, V. M. et al. MRI signatures of brain age and disease over the lifespan based on a deep brain network and 14 468 individuals worldwide. Brain 143, 2312–2324 (2020).

48. Prins, N. D. & Scheltens, P. White matter hyperintensities, cognitive impairment and dementia: an update. Nat Rev Neurol 11, 157–165 (2015).

49. Andreasen, N. et al. Evaluation of CSF-tau and CSF-Aβ42 as Diagnostic Markers for Alzheimer Disease in Clinical Practice. Archives of Neurology 58, 373–379 (2001).

50. Buniello, A. et al. The NHGRI-EBI GWAS Catalog of published genome-wide association studies, targeted arrays and summary statistics 2019. Nucleic Acids Res 47, D1005–D1012 (2019).

51. Bulik-Sullivan, B. K. et al. LD Score regression distinguishes confounding from polygenicity in genome-wide association studies. Nat Genet 47, 291–295 (2015).

52. Jiang, L., Zheng, Z., Fang, H. & Yang, J. A generalized linear mixed model association tool for biobank-scale data. Nat Genet 53, 1616–1621 (2021).

53. Watanabe, K., Taskesen, E., van Bochoven, A. & Posthuma, D. Functional mapping and annotation of genetic associations with FUMA. Nat Commun 8, 1826 (2017).

54. Lanni, C., Masi, M., Racchi, M. & Govoni, S. Cancer and Alzheimer’s disease inverse relationship: an age-associated diverging derailment of shared pathways. Mol Psychiatry 26, 280–295 (2021).

55. Fisher, Y., Nemirovsky, A., Baron, R. & Monsonego, A. Dendritic cells regulate amyloid-β-specific T-cell entry into the brain: the role of perivascular amyloid-β. J Alzheimers Dis 27, 99–111 (2011).

56. Brezovakova, V., Valachova, B., Hanes, J., Novak, M. & Jadhav, S. Dendritic Cells as an Alternate Approach for Treatment of Neurodegenerative Disorders. Cell Mol Neurobiol 38, 1207–1214 (2018).

57. Kong, D. et al. Niacin Promotes Cardiac Healing after Myocardial Infarction through Activation of the Myeloid Prostaglandin D2 Receptor Subtype 1. J Pharmacol Exp Ther 360, 435–444 (2017).

58. Crouse, J. R. New developments in the use of niacin for treatment of hyperlipidemia: new considerations in the use of an old drug. Coron Artery Dis 7, 321–326 (1996).

59. Duggal, J. K. et al. Effect of niacin therapy on cardiovascular outcomes in patients with coronary artery disease. J Cardiovasc Pharmacol Ther 15, 158–166 (2010).

60. Moutinho, M. et al. The niacin receptor HCAR2 modulates microglial response and limits disease progression in a mouse model of Alzheimer’s disease. Sci Transl Med 14, eabl7634 (2022).

61. Wen, J. et al. Novel genomic loci and pathways influence patterns of structural covariance in the human brain. 2022.07.20.22277727 Preprint at 10.1101/2022.07.20.22277727 (2022).

62. Yang, J. et al. Microtubule-associated protein tau is associated with the resistance to docetaxel in prostate cancer cell lines. Res Rep Urol 9, 71–77 (2017).

63. Bernatz, S. et al. Impact of Docetaxel on blood-brain barrier function and formation of breast cancer brain metastases. J Exp Clin Cancer Res 38, 434 (2019).

64. Shin, W. et al. A first-in-human study to investigate the safety, tolerability, pharmacokinetics, and pharmacodynamics of KM-819 (FAS-associated factor 1 inhibitor), a drug for Parkinson’s disease, in healthy volunteers. Drug Des Devel Ther 13, 1011–1022 (2019).

65. Kim, B.-S. et al. Pharmacological Intervention Targeting FAF1 Restores Autophagic Flux for α-Synuclein Degradation in the Brain of a Parkinson’s Disease Mouse Model. ACS Chem Neurosci 13, 806–817 (2022).

66. Swerdlow, R. H., Burns, J. M. & Khan, S. M. The Alzheimer’s disease mitochondrial cascade hypothesis: progress and perspectives. Biochim Biophys Acta 1842, 1219–1231 (2014).

67. Hopkins, A. L. & Groom, C. R. The druggable genome. Nat Rev Drug Discov 1, 727–730 (2002).

68. Pushpakom, S. et al. Drug repurposing: progress, challenges and recommendations. Nat Rev Drug Discov 18, 41–58 (2019).

69. Jiang, C., Li, G., Huang, P., Liu, Z. & Zhao, B. The Gut Microbiota and Alzheimer’s Disease. J Alzheimers Dis 58, 1–15 (2017).

70. Seo, D., Boros, B. D. & Holtzman, D. M. The microbiome: A target for Alzheimer disease? Cell Res 29, 779–780 (2019).

71. Heppner, F. L., Ransohoff, R. M. & Becher, B. Immune attack: the role of inflammation in Alzheimer disease. Nat Rev Neurosci 16, 358–372 (2015).

72. Kinney, J. W. et al. Inflammation as a central mechanism in Alzheimer’s disease. Alzheimers Dement (N Y) 4, 575–590 (2018).

73. Fujiwara, N. & Kobayashi, K. Macrophages in inflammation. Curr Drug Targets Inflamm Allergy 4, 281–286 (2005).

74. Laurin, D., David Curb, J., Masaki, K. H., White, L. R. & Launer, L. J. Midlife C-reactive protein and risk of cognitive decline: a 31-year follow-up. Neurobiol Aging 30, 1724–1727 (2009).

75. Thambisetty, M. et al. Association of Plasma Clusterin Concentration With Severity, Pathology, and Progression in Alzheimer Disease. Archives of General Psychiatry 67, 739–748 (2010).

76. Ruparelia, N., Chai, J. T., Fisher, E. A. & Choudhury, R. P. Inflammatory processes in cardiovascular disease: a route to targeted therapies. Nat Rev Cardiol 14, 133–144 (2017).

77. Vitale, G., Salvioli, S. & Franceschi, C. Oxidative stress and the ageing endocrine system. Nat Rev Endocrinol 9, 228–240 (2013).

78. Strittmatter, W. J. Old Drug, New Hope for Alzheimer’s Disease. Science 335, 1447–1448 (2012).

79. Strittmatter, W. J. Medicine. Old drug, new hope for Alzheimer’s disease. Science 335, 1447–1448 (2012).

80. Clark, I. A. & Atwood, C. S. Is TNF a Link between Aging-Related Reproductive Endocrine Dyscrasia and Alzheimer’s Disease? JAD 27, 691–699 (2011).

81. Clark, I. A., Alleva, L. M. & Vissel, B. The roles of TNF in brain dysfunction and disease. Pharmacol Ther 128, 519–548 (2010).

82. Demetrius, L. A. & Driver, J. Alzheimer’s as a metabolic disease. Biogerontology 14, 641–649 (2013).

83. Jack, C. R. et al. NIA-AA Research Framework: Toward a biological definition of Alzheimer’s disease. Alzheimers Dement 14, 535–562 (2018).

84. Gazestani, V. et al. Early Alzheimer’s disease pathology in human cortex involves transient cell states. Cell 186, 4438–4453.e23 (2023).

85. Xiong, X. et al. Epigenomic dissection of Alzheimer’s disease pinpoints causal variants and reveals epigenome erosion. Cell 186, 4422–4437.e21 (2023).

86. Sun, N. et al. Human microglial state dynamics in Alzheimer’s disease progression. Cell 186, 4386–4403.e29 (2023).

87. Dileep, V. et al. Neuronal DNA double-strand breaks lead to genome structural variations and 3D genome disruption in neurodegeneration. Cell 186, 4404–4421.e20 (2023).

88. Mathys, H. et al. Single-cell atlas reveals correlates of high cognitive function, dementia, and resilience to Alzheimer’s disease pathology. Cell 186, 4365–4385.e27 (2023).

89. Luo, W., Qu, W. & Gan, L. The AD odyssey 2023: Tales of single cell. Cell 186, 4257– 4259 (2023).

90. Tian, Y. E. et al. Biological aging of human body and brain systems. 2022.09.03.22279337 Preprint at 10.1101/2022.09.03.22279337 (2022).

91. Zhao, B. et al. Heart-brain connections: Phenotypic and genetic insights from magnetic resonance images. Science 380, abn6598 (2023).

92. Wen, J. et al. The Genetic Architecture of Biological Age in Nine Human Organ Systems. medRxiv 2023.06.08.23291168 (2023) doi:10.1101/2023.06.08.23291168.

93. Liberzon, A. et al. Molecular signatures database (MSigDB) 3.0. Bioinformatics 27, 1739–1740 (2011).

94. Wishart, D. S. et al. DrugBank 5.0: a major update to the DrugBank database for 2018. Nucleic Acids Res 46, D1074–D1082 (2018).

95. Zhou, Y. et al. Therapeutic target database update 2022: facilitating drug discovery with enriched comparative data of targeted agents. Nucleic Acids Res 50, D1398–D1407 (2022).

