## Supplementary materials for "Genetic, Clinical Underpinnings of Brain Change Along Two Neuroanatomical Dimensions of Clinically-defined Alzheimer’s Disease"

**Running title**: Two AI Dimensions of Alzheimer's Disease
**Keywords:** Alzheimer's disease; dimensions; imaging-genomics; deep learning; semi-supervised representation learning.

**eMethod 1: Surreal-GAN: methodological considerations and advances**
**eMethod 2: Study participants and populations**
**eMethod 3: Image quality check**
**eMethod 4: Genetic data quality check protocol**
**eMethod 5: Annotation of genomic loci and gene mappings**
**eMethod 6: Prioritized gene set enrichment and tissue specificity analysis**
**eMethod 7: Annotation of novel genomic loci and genes related to AD**
**eMethod 8: Polygenic risk score calculation**
**eMethod 9: Target-Drug-disease network for "druggable genes"**
**eText 1: Sensitivity check analysis for the main GWASs of Euroepan ancestry in the general population**
**eFigure 1: The degree of normality for MUSE ROIs before and after statistical harmonization**
**eFigure 2: Distributions of the R1 and R2 dimensions of Surreal-GAN and the P1, P2, P3, and P4 dimensions of Smile-GAN**
**eFigure 3: Pearson's correlation coefficient between the R1 and R2 dimensions of Surreal-GAN and the P1, P2, P3, and P4 dimensions of Smile-GAN**
**eFigure 4: The descriptive statistics of the expression of the R1 and R2 dimensions in the four populations**
**eFigure 5: Longitudinal conversion of MCI to AD and between R1- and R2-dominant groups in the MCI/AD population**
**eFigure 6: PRS plots at baseline for the MCI/AD population**
**eFigure 7: QQ plots for the baseline GWAS for the MCI/AD population**
**eFigure 8: Manhattan and QQ plots for the longitudinal GWAS for the general population**
**eFigure 9: Tissue specificity analyses for the prioritized genes in the two dimensions in the general population**
**eFigure 10: Singe gene expression heat maps for the tissue specificity analyses in the general population**
**eFigure 11: The longitudinal rate of change of R1 and R2 neuroanatomical dimensions of brain atrophy in the MCI/AD population.**
**eTable 1: Brain association studies for the MCI/AD population from ADNI and BLSA**
**eTable 2: Identified genomic loci and mapped genes for baseline GWAS in the MCI/AD population**
**eTable 3: Clinical association studies for the MCI/AD population from ADNI and BLSA**
**eTable 4: Brain association studies for the general population from UKBB**
**eTable 5: Clinical association studies for the general population from UKBB**
**eTable 6: Identified genomic loci and mapped genes for GWAS in the general population**
**eTable 7: Gene set enrichment analysis results in the general population**
**eTable 8: Brain association studies for the cognitively unimpaired (CU) population from ADNI and BLSA**
**eTable 9: Clinical association studies for the cognitively unimpaired (CU) population from ADNI and BLSA**

**eMethod 1: Surreal-GAN: methodological considerations and advances**

Surreal-GAN[1] dissects underlying disease-related heterogeneity via a deep representation learning approach under the principle of semi-supervised clustering.[2,3] The methodological advance of Surreal-GAN is to model neuroanatomical heterogeneity by considering both spatial and temporal (i.e., disease severity) variation using only cross-sectional MRI data. The following schematic figure demonstrates the principles of Smile-GAN[2] (its precursor) and Surreal-GAN in the semi-supervised learning framework to disentangle AD neuroanatomical heterogeneity.

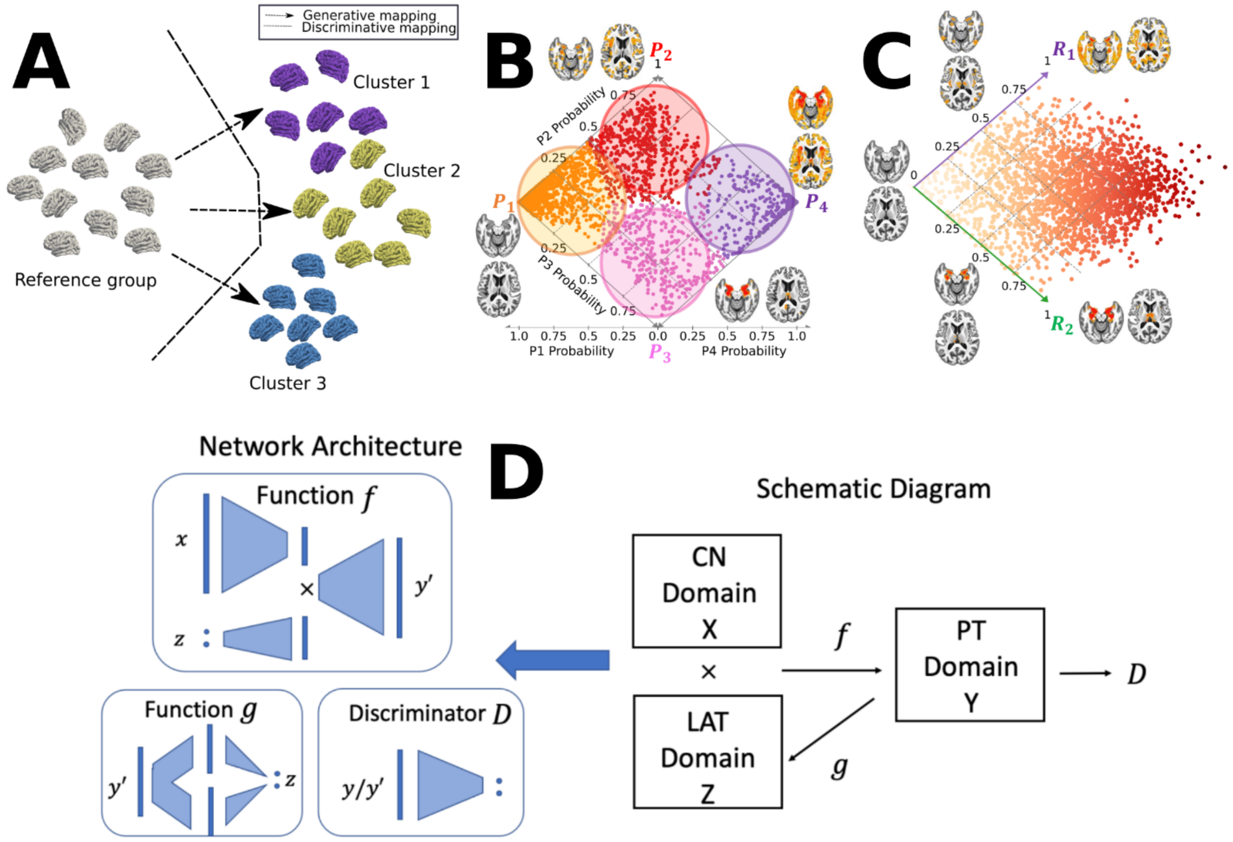


**Fig**. **A**) Schematic figures of semi-supervised clustering methods, including generative[1,2,4] and discriminative approaches.[3,5] Semi-supervised clustering methods dissect the neuroanatomical heterogeneity of brain diseases by seeking the so-called "*1-to-k*" mapping between the healthy control (CN) group as a reference and the patient (PT) group as a target. They sought to tease out clusters that are likely driven by distinct pathological trajectories instead of by global similarity/dissimilarity in data. The figure is adapted from our previous work.[6] **B**) Conceptual illustration of applying Smile-GAN ADNI to cluster MCI/AD participants into four hard-coded subtypes (P1, P2, P3, and P4) and two longitudinal pathways after applying the model to longitudinal data: P1→P2→P4; *ii*) P1→P3→P4. The figure is adapted from our previous work.[2] **C**) Conceptual illustration of applying Surreal-GAN to ADNI to recapitulate the neuroanatomical heterogeneity of MCI/AD participants into two continuous dimensional scores. Methodological advances are to capture disease severity and heterogeneity simultaneously using only cross-sectional data and allow the same individuals to express themselves along multiple dimensions. The figure is adapted from our previous work.[1] **D**) General network architectures (left) and schematic diagram of the semi-supervised learning (right, corresponding to Fig. A). The figure is adapted from our previous work.[1]

Smile-GAN learns one transformation function, *f*, which generates synthesized PT data, $\boldsymbol{y}' = f(\boldsymbol{x}, \boldsymbol{z})$, from real CU data $\boldsymbol{x}$ and the latent variable $\boldsymbol{z}$**.** The categorical variable $\boldsymbol{z}$ is encoded as an *M*-dimensional one-hot vector with value 1 being placed at any position with equal probability (i.e., *1/k*). Thus, $k$ different values of $\boldsymbol{z}$ lead to $k$ different mapping directions and enable *M* subgroups clustering.

Surreal-GAN also learns one transformation function $f$, which transforms CU data $\boldsymbol{x}$ to different synthesized PT data $\boldsymbol{y}' = f(\boldsymbol{x}, \boldsymbol{z})$, with latent variable $\boldsymbol{z}$ specifying distinct mapping directions. However, compared to Smile-GAN, Surreal-GAN considers that disease heterogeneity spatially and temporally (subtype) expands along a continuum (severity), similar to Sustain[7] to a certain extent. Thus, the latent variable $\boldsymbol{z}$ is modeled as a continuous variable, allowing infinite mapping directions from CU to PT data. $\boldsymbol{z} \sim p_{lat}(\boldsymbol{z})$ is sampled from a multivariate uniform distribution $U(0,1)^k$, rather than a categorical distribution as in Smile-GAN. Further, Surreal-GAN aims at disentangling spatial and temporal variations independently so that each dimension of the latent variable $\boldsymbol{z}$ is correlated with the severity of one relatively homogeneous imaging pattern. In contrast, different dimensions can be associated with spatially different imaging patterns.

To achieve the goals above, the objective function of Surreal-GAN consists of one adversarial loss and other regularization terms. The adversarial loss aims at matching the distribution of synthesized PT data, $p_{syn}$, and the distribution of real PT data, $p_{PT}$. Other regularization terms serve the following purposes: 1) encouraging sparse transformations (change loss); 2) reconstructing latent variables from synthesized or real PT data through the decomposer $g_1$ (decomposition loss) and reconstruction function $g_2$ (reconstruction loss); 3) boosting spatial separation of synthesized/captured patterns (orthogonality loss); 4) enforcing positive correlations between components of $\boldsymbol{z}$ and severity of synthesized patterns (monotonicity loss and cn loss).

Specifically, with distributions of CU, real PT, synthesized PT data denoted as $p_{CU}(\boldsymbol{x})$, $p_{PT}(\boldsymbol{y})$ and $p_{syn}(\boldsymbol{y}')$ respectively, the adversarial loss is defined as:

$$L_{GAN}(D, f) = E_{\boldsymbol{y}\sim p_{PT}(\boldsymbol{y})}\left[log\left(D(\boldsymbol{y})\right)\right] + E_{\boldsymbol{z}\sim p_{Lat}(\boldsymbol{z}),\boldsymbol{x}\sim p_{CN}(\boldsymbol{x})}\left[1 - log\left(D\left(f(\boldsymbol{x}, \boldsymbol{z})\right)\right)\right]$$
$$= E_{\boldsymbol{y}\sim p_{PT}(\boldsymbol{y})}\left[log\left(D(\boldsymbol{y})\right)\right] + E_{p_{syn}(\boldsymbol{y}')}\left[1 - log\left(D(\boldsymbol{y}')\right)\right]$$

The transformation function attempts to synthesize PT data $\boldsymbol{y}'$, so that they follow similar distributions as real PT data. The discriminator, $D$, tries to distinguish the synthesized PT data from real PT data. Therefore, the discriminator is updated to maximize the adversarial loss, while the transformation function is optimized to minimize it.

Other regularization terms are introduced to further regularize the transformation function $f$. With the assumption that the disease process will not change brain anatomy dramatically and primarily only affect certain regions throughout most of the disease stages, the change loss is introduced to control sparsity and distance of transformations:

$$L_{change}(f) = E_{\boldsymbol{z}\sim p_{Lat}(\boldsymbol{z}),\boldsymbol{x}\sim p_{CU}(\boldsymbol{x})}[\|f(\boldsymbol{x}, \boldsymbol{z}) - \boldsymbol{x}\|_1]$$

The decomposer $g_1$ serves to reconstruct changes synthesized by each component: $\boldsymbol{q}_i = f(\boldsymbol{x}, \boldsymbol{a}^{\boldsymbol{i}}) - \boldsymbol{x}$, where $\boldsymbol{a}_{\boldsymbol{i}}^{\boldsymbol{i}} = \boldsymbol{z}_{\boldsymbol{i}}$ and $\boldsymbol{a}_{\boldsymbol{j}}^{\boldsymbol{i}} = 0$ for $j \neq i$. With $\widehat{\boldsymbol{q}}_{f(x,z)} = [\boldsymbol{q}_1^T, \boldsymbol{q}_2^T, \cdots, \boldsymbol{q}_M^T]^T$, the decomposition loss is defined as:

$$L_{decom}(f, g_1) = E_{\boldsymbol{z}\sim p_{Lat}(\boldsymbol{z}),\boldsymbol{x}\sim p_{CN}(\boldsymbol{x})}\left[\left\|g_1(f(\boldsymbol{x}, \boldsymbol{z})) - \widehat{\boldsymbol{q}}_{f(\boldsymbol{x},\boldsymbol{z})}\right\|_2\right]$$

The reconstruction function $g_2$ serves to further reconstruct each component of the sampled $\boldsymbol{z}$ variable from $g_1\left(f(\boldsymbol{x}, \boldsymbol{z})\right)$. The function $g$ is defined as a composition of $g_1$ and $g_2$, with $g\left(f(\boldsymbol{x}, \boldsymbol{z})\right) = \left[g_2\left(g_1\left(f(\boldsymbol{x}, \boldsymbol{z})\right)_{0:S}\right), \cdots, g_2\left(g_1\left(f(\boldsymbol{x}, \boldsymbol{z})\right)_{S*(M-1):S*M}\right)\right]^T$ (S = number of ROIs), and the reconstruction loss is defined as

$$L_{recons}(f, g) = L_{recons}(f, g_1, g_2) = E_{\boldsymbol{z}\sim p_{Lat}(\boldsymbol{z}),\boldsymbol{x}\sim p_{CN}(\boldsymbol{x})}\left[\left\|g\left(f(\boldsymbol{x}, \boldsymbol{z})\right) - \boldsymbol{z}\right\|_2\right]$$

The orthogonality loss aims to boost changes led each component, $\boldsymbol{q}_i$, to be relatively orthogonal to each other. For this purpose, a matrix $\boldsymbol{A}_{f(\boldsymbol{x},\boldsymbol{z})}$ is constructed with the $i_{th}$ column $A_{f(\boldsymbol{x},\boldsymbol{z})_{:,i}} = \boldsymbol{q}_i/||\boldsymbol{q}_i||_2$, and the orthogonality loss is defined as:

$$L_{ortho}(f) = E_{\boldsymbol{z}\sim p_{Lat}(z),\boldsymbol{x}\sim p_{CU}(\boldsymbol{x})}[||\boldsymbol{A}_{f(\boldsymbol{x},\boldsymbol{z})}^T\boldsymbol{A}_{f(\boldsymbol{x},\boldsymbol{z})} - I||_F]$$

To encourage a positive correlation between the severity of the synthesized pattern and the value of each component $\boldsymbol{z}_i$, another latent variable $\boldsymbol{z}' \sim p_{sev}(\boldsymbol{z}'|\boldsymbol{z})$ is sampled conditioned on previously sampled $\boldsymbol{z}$ variables, such that $\boldsymbol{z}_i' \geq \boldsymbol{z}_i$ for any $1 \leq i \leq k$. With these two sampled latent variables, the monotonicity loss is defined as:

$$L_{mono}(f) = E_{\boldsymbol{z} \sim p_{Lat}(\boldsymbol{z}), \boldsymbol{z} \sim p_{sev}(\boldsymbol{z}'|\boldsymbol{z}), \boldsymbol{x} \sim p_{CU}(\boldsymbol{x})}[||max\ (|f(\boldsymbol{x}, \boldsymbol{z}) - \boldsymbol{x}| - |f(\boldsymbol{x}, \boldsymbol{z}') - \boldsymbol{x}|, 0)||_2]$$

The cn loss is introduced to further ensure that a small $\boldsymbol{z}$ variable lead to mild patterns. By letting $p_{cn}(\boldsymbol{z}) = U(0, 0.05)^k$ to be a multivariate uniform distribution, the cn loss is defined as:

$$L_{cn}(f) = E_{\boldsymbol{z}^{cn} \sim p_{cn}(\boldsymbol{z}), \boldsymbol{x} \sim p_{CU}(\boldsymbol{x})}[\|f(\boldsymbol{x}, \boldsymbol{z}^{\boldsymbol{cn}}) - \boldsymbol{x}\|_1]$$

With all loss functions introduced above, the full objective of Surreal-GAN can be written as

$$L(D, f, g_1, g_2) = L_{GAN}(D, f) + \gamma L_{change}(f) + \kappa L_{decom}(f, g_1) + \zeta L_{recon}(f, g_1, g_2) + \lambda L_{ortho}(f) + \mu L_{mono}(f) + \eta L_{cn}(f)$$

With $\gamma$, $\kappa$, $\zeta$, $\lambda$, $\mu$, and $\eta$ being hyperparameters that control the relative importance of each loss function during the training process. More details of parameter selections can be found in the Surreal-GAN paper.[1]

Through the training process, parametrized functions $f$ , $g_1$, and $g_2$ are updated to satisfy that:

$$f, g_1, g_2 = arg \min_{f,g} \max_{D} L(D, f, g_1, g_2)$$

More importantly, after the training process, the function g, a composition of $g_1$ and $g_2$, can be applied to unseen PT data to infer the latent variable, which is referred to as the R-indices (i.e., dimensions) of PT data.

By modeling, Surreal-GAN's R1 and R2 dimensions are more appropriate for dimensional analyses as continuous variables. In contrast, the four neuroanatomical subtypes of Smile-GAN are better instruments for case-control analyses as categorical variables. We used ADNI data to show that the R1 and R2 dimensional scores were approximately normally distributed, and the P1, P2, P3, and P4 probability scores are not normally distributed (**Supplementary eFigure 8**).

**eMethod 2: Study participants and populations**

For ADNI, cognitively unimpaired (CU), MCI, and AD participants were recruited from multiple centers in the United States. The primary goal of ADNI was to derive the R1 and R2 dimensions by applying the Surreal-GAN model to T1w MRIs. We included all baseline and longitudinal T1w MRI scans and cognitive data available from ADNI in iSTAGING. In addition, we also included whole-genome sequencing (WGS) data for ADNI in AI4AD.

BLSA is a longitudinal study that aims to study aging and related diseases, such as AD. At baseline recruitment, participants were mostly cognitively normal and had multiple time points of longitudinal follow-ups for MRIs and cognition.

Participants enrolled in PREVENT-AD were cognitively normal older adults with a family history of AD (at least one parent or multiple siblings)[a]. The inclusion criteria are generally similar but more stringent than the proxy-AD diagnosis[8,9] used in UK Biobank (see below), including *i*) being cognitively normal, *ii*) having a family history of AD, *iii*) aging within 15 years from the age of disease onset of their youngest relative, and *iv*) no history of neurological or psychiatric diseases.

For UKBB, we defined asymptomatic participants[b] as those who did not have a diagnosis of Alzheimer's disease (G30 in ICD-10 diagnoses, see below) in our data consolidation. However, these asymptomatic participants might have diagnoses of other illnesses or comorbidities based on ICD-10. Furthermore, we defined proxy-AD[c] in UKBB as long as the participant satisfied one of the following criteria: *i*) illnesses_of_father_f20107 and *ii*) illnesses_of_mother_f20110.

**eMethod 3: Image quality check**

Raw T1-weighted MRIs were first quality checked (QC) for motion, image artifacts, or restricted field-of-view. Another QC was performed as follows: First, the images were examined by manually evaluating for pipeline failures (e.g., poor brain extraction, tissue segmentation, and registration errors). Furthermore, a second step automatically flagged images based on outlying values of quantified metrics (i.e., PSC values); those flagged images were re-evaluated.

**eMethod 4: Genetic data quality check protocol**

For ADNI WGS data, we first convert the VCF files into *plink* binary format. We excluded related individuals (up to 2nd-degree) using the KING software for family relationship inference.[10] Further QC steps are: excluding criteria were: i) individuals with more than 2% of missing genotypes; ii) variants with minor allele frequency (MAF) of less than 0.1%; iii) variants with larger than 5% missing genotyping rate; iv) variants that failed the Hardy-Weinberg test at $1x10^{-5}$. We then removed duplicated variants from all 22 autosomal chromosomes. We also excluded individuals for whom either imaging or genetic data were not available. To adjust for population stratification,[11] we derived the first 40 genetic principal components (PC) using the SmartPCA software[12].

For UKBB, the genetic pipeline was previously described elsewhere.[13,14] All QC steps were documented in our BRIDGEPORT web portal: https://www.cbica.upenn.edu/bridgeport/data/pdf/BIGS_genetic_protocol.pdf. First, we excluded related individuals (up to 2nd-degree) from the complete UKBB sample using the KING software for family relationship inference[10]. We then removed duplicated variants from all 22 autosomal chromosomes. Individuals whose genetically identified sex did not match their self-acknowledged sex were removed. Other excluding criteria were: i) individuals with more than 3% of missing genotypes; ii) variants with minor allele frequency (MAF) of less than 1%; iii) variants with larger than 3% missing genotyping rate; iv) variants that failed the Hardy-Weinberg test at $1x10^{-10}$. To adjust for population stratification[11], we derived the first 40 genetic principle components (PC) using the FlashPCA software[15]. Details of the genetic quality check protocol are described elsewhere[13,16].

**eMethod 5: Annotation of genomic loci and gene mappings**

The annotation of genomic loci and gene mappings was performed on the online platform of FUMA (*SNP2GENE*, https://fuma.ctglab.nl/, version: v1.3.8). For the annotation of genomic loci, default parameters were set in FUMA. First, lead SNPs (correlation $r^2 \leq 0.1$, distance < 250 kilobases) are assigned to a genomic locus (non-overlapping). The SNP with the lowest P-value represents each genomic locus. For gene mappings, three different strategies were used to map the SNPs to genes. First, positional mapping maps SNPs to genes if the SNPs are physically located inside a gene (a 10 kb window by default). Second, expression quantitative trait loci (eQTL) mapping maps SNPs to genes showing a significant eQTL association. Lastly, chromatin interaction mapping maps SNPs to genes when there is a significant chromatin interaction between the disease-associated regions and nearby or distant genes.[17]

**eMethod 6: Prioritized gene set enrichment and tissue specificity analysis**

FUMA provides the functionality *GENE2FUNC* to study the expression of prioritized genes and test for enrichment of the set of genes in pre-defined pathways. We used the mapped genes as prioritized gene inputs. The background genes were specified as all genes in FUMA, and default values were defined for all other parameters. *GENE2FUNC* outputs a single gene-level expression heat map that quantifies the expression values (average expression per label or average of normalized expression per label) in different tissues, including the GTEx v8[18] 54 tissue types and 30 general tissue types. For tissue specificity analysis, differentially expressed gene sets (DEG) were pre-calculated by performing a two-sided t-test for any one label of tissue against all others. Input genes were tested against the pre-defined DEG sets using the hypergeometric test. The tissue specificity plot highlights significant enrichment at the Bonferroni corrected P-value < 0.05.

**eMethod 7: Annotation of novel genomic loci and genes related to AD**

A two-step procedure was performed to determine if a genomic locus or gene was associated with AD-related clinical traits. First, we manually queried the identified genomic loci, mapped genes, checked if any AD-related traits were previously reported in GWAS Catalog, and downloaded these associations. In addition, we checked if any input genes overlap with the gene set pathways (defined in GWAS catalog reported genes) related to AD. We defined a genomic locus or a gene as a novel association if the variant was not associated with any clinical traits in GWAS Catalog. For these clinical traits reported in GWAS Catalog, we mapped them into several different categories.

**eMethod 8: Polygenic risk score calculation**

We calculated the PRS[19] using both ADNI and UKBB genetic data. The weights of the PRS were defined based on independent base data,[20] ensuring that the base population does not overlap with the target population in ADNI and UKBB (European ancestry). The QC steps for the base data are as follows: *i*) SNP-based heritability estimate ($h^2 > 0.05$) using LDSC[21] to avoid spurious SNP data; *ii*) the genome reference consortium human build of the base data is on GRCh37[22]; *iii*) removal of duplicated and ambiguous SNPs. The QC steps for the target data are as follows: *i*) using LiftOver[23] to convert the ADNI WGS from GRCh38 to GRCh37; *ii*) standard GWAS QC (low minor allele frequency, low genotyping rate, etc.); *iii*) pruning to remove highly correlated SNPs; *iv*) removal of high heterozygosity samples; *v*) removal of duplicated, mismatching and ambiguous SNPs. After rigorous QC, we used PLINK to generate PRS for ADNI and UKBB by adopting the classic C+T method (clumping + thresholding: C+T). To approximate the "best-fit" PRS, we performed a logistic regression using the PRS calculated at different P-value thresholds, controlling for age, sex, and the first five genetic PCs. We chose the PRS that explains the highest phenotypic variance (AD vs. CU in ADNI).

**eMethod 9: Target-Drug-disease network for “druggable genes”**

We constructed a target-drug-disease network from the DrugBank database (v.5.1.9)[24] and the Therapeutic Target Database (updated by September 29th, 2021).[25] Specifically, we constrained the target to human organisms and included all drugs with different highest statuses (e.g., patented, approved, etc.). We further mapped the drug to the Anatomical Therapeutic Chemical (ATC) classification system for the Drugbank database and the International Classification of Diseases (ICD-11) for the Therapeutic Target Database. These resulted in 5927 drugs (including drug-like molecules), 2775 target genes (i.e., druggable genome), and 1770 diseases classified by the ATC code (5th level) for the DrugBank database, and 11,343 drugs, 1785 target genes, and 1101 diseases defined by the ICD-11 code. We defined a gene as a “druggable gene” if the target gene expresses proteins to bind drug-like molecules and the drug is at any clinical trial stage.

**eText 1: Sensitivity check analysis for the main GWASs of Euroepan ancestry in the general population**

**Split-sample GWAS**

In the GWAS from the first split, we found 74 and 409 R-SNP associations for R1 and R2 (P-value < $5x10^{-8}$). Of these, 74 (71) and 379 (267) associations were replicated in the second split GWAS using both the nominal P-value (<0.05, 100% and 92.66% concordance rates for R1 and R2) and the Bonferroni-corrected threshold (<0.05/*N*, 95.94% and 65.28% concordance rates for R1 and R2). Detailed results are presented in **Supplementary eFile 2**.

**sex-stratified GWAS**

For the female-specific GWAS, we observed 52 and 234 R-SNP associations for R1 and R2. Among these, 52 (42) and 195 (195) associations were replicated in the male-specific GWAS using both the nominal P-value (<0.05, 100% and 83.3% concordance rates for R1 and R2) and the Bonferroni-corrected threshold (<0.05/*N*, 80.76% and 83.3% concordance rates for R1 and R2). While we observe fewer significant loci (P-value < $5x10^{-8}$) in the sensitivity analyses than the results obtained from the full sample sizes, which may be due to reduced sample sizes, the signal peaks of the P-values remain consistent, as well as the effect directions. Detailed results are presented in **Supplementary eFile 3**.

**fastGWA for linear mixed model**

In the GWAS using PLINK linear models, we found 437, and 735 R-SNP associations for R1 and R2 (P-value < $5x10^{-8}$). All these associations were replicated in the fastGWA GWAS using both the nominal P-value (<0.05) and the Bonferroni-corrected threshold (<0.05/*N*). Detailed results are presented in **Supplementary eFile 4**.

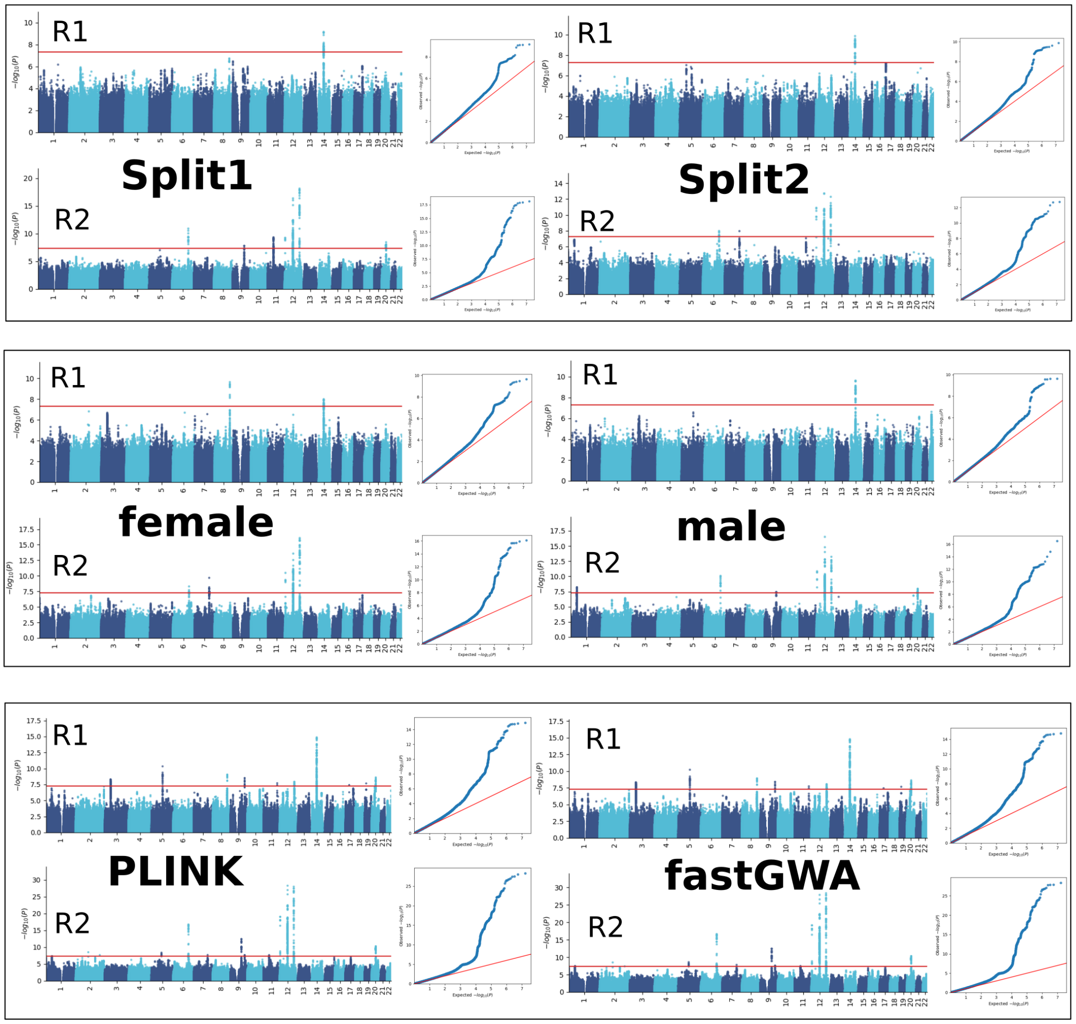


Split-sample, sex-stratified, and fastGWA GWASs for the sensitivity check analyses. Manhattan and QQ plots are shown.

**eFigure 1: The degree of normality for MUSE ROIs before and after statistical harmonization**

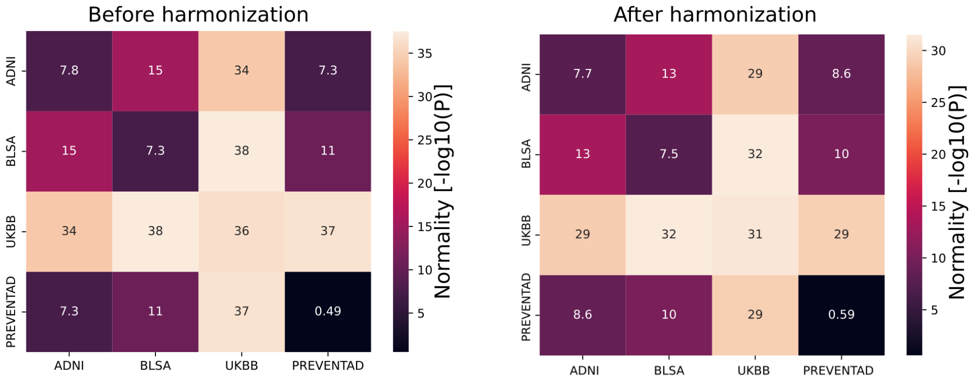


We showcased the degree of normality of one MUSE GM ROI (right accumbens area) from each pair of sites using the Shapiro-Wilk test (*scipy.stats.shapiro function*) before (left) harmonization and after harmonization (right). A higher -log10(P) indicates the data are less likely to be normally distributed. As a general trend, our statistical harmonization techniques demonstrated a slight improvement in the normality of the data. Additionally, we consistently applied normality transformations to all statistical analyses, including GWAS, to mitigate any additional non-normality.

**eFigure 2: Distributions of the R1 and R2 dimensions of Surreal-GAN and the P1, P2, P3, and P4 dimensions of Smile-GAN**

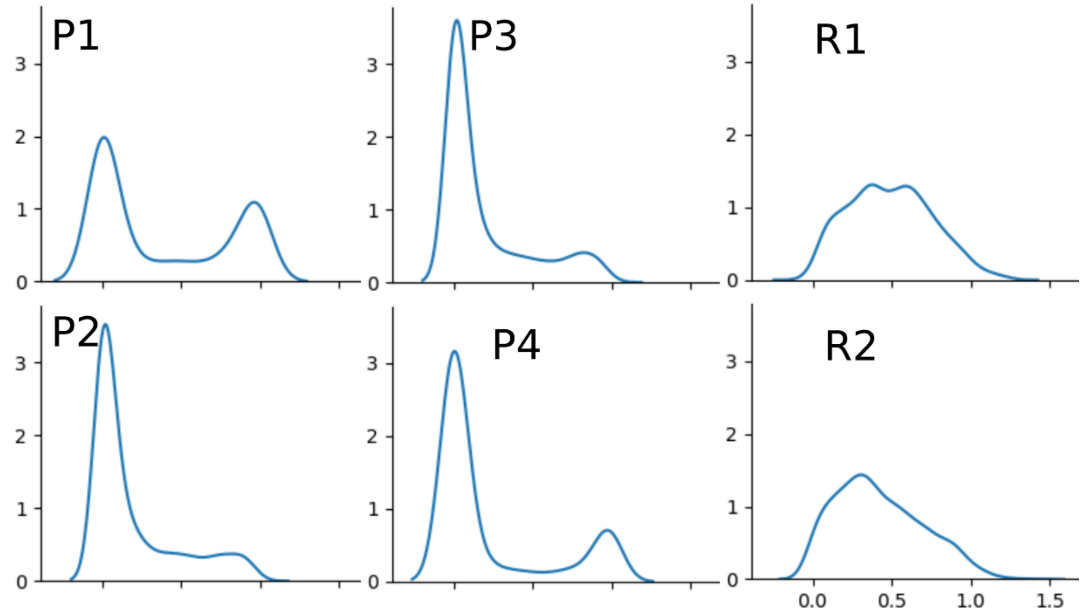


By modeling, Surreal-GAN's R1 and R2 dimensions are more appropriate and statistically powerful for continuous trait-based GWAS analyses as continuous dimensional scores. In contrast, the four subtype probability scores of Smile-GAN are better instruments for case-control GWAS as categorical variables. We used ADNI data to show that the R1 and R2 dimension scores were approximately normally distributed, and the P1, P2, P3, and P4 dimension scores followed bimodal distributions. Of note, we performed an inverse-normal transformation[31] to enable the phenotype/trait of interest to be normally distributed.

**eFigure 3: Pearson's correlation coefficient between the R1 and R2 dimensions of Surreal-GAN and the P1, P2, P3, and P4 dimensions of Smile-GAN**

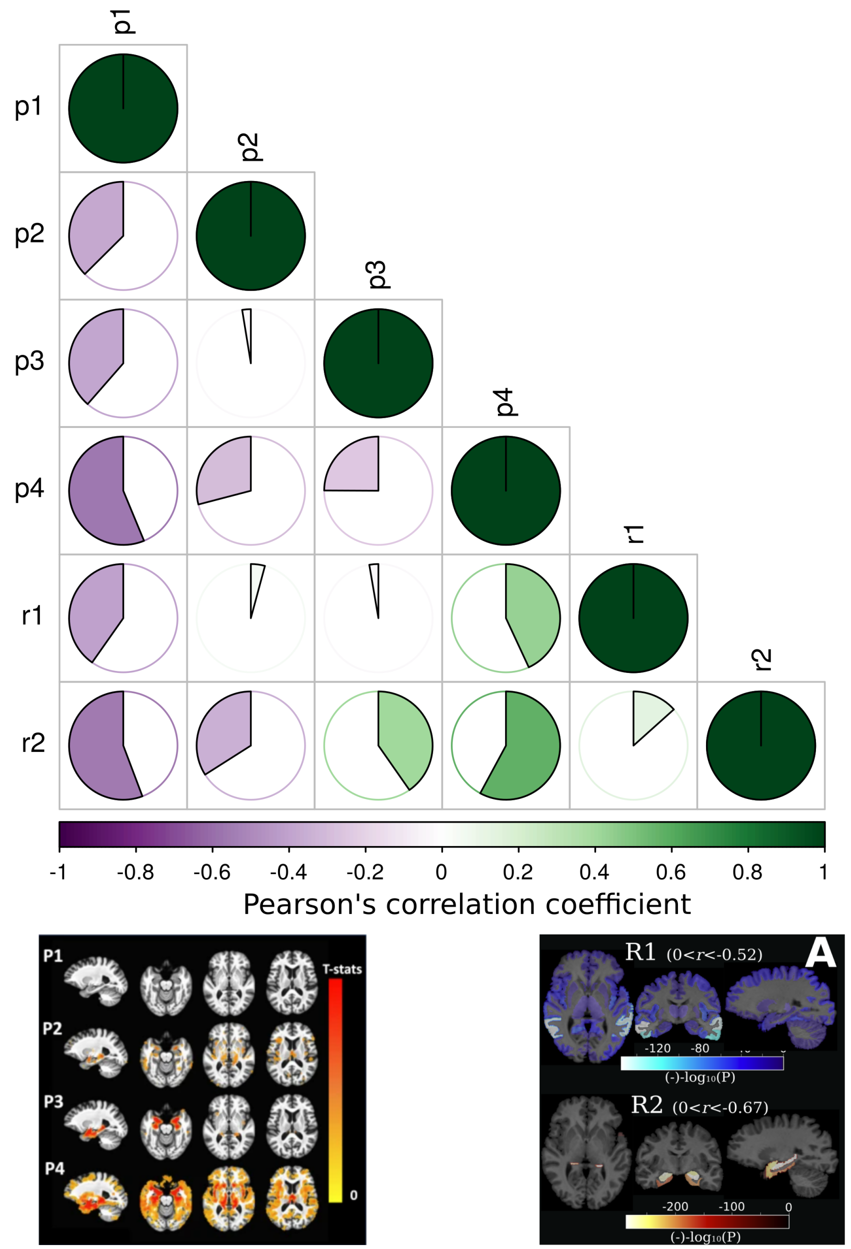


We calculated Pearson's correlation coefficient in the ADNI dataset. We also show here the imaging patterns of each dimension. The figure for the P1-P4 from the Smile-GAN model is originally from our previous work (Yang et al., 2019[2]).

**eFigure 4: The descriptive statistics of the expression of the R1 and R2 dimensions in the four populations**

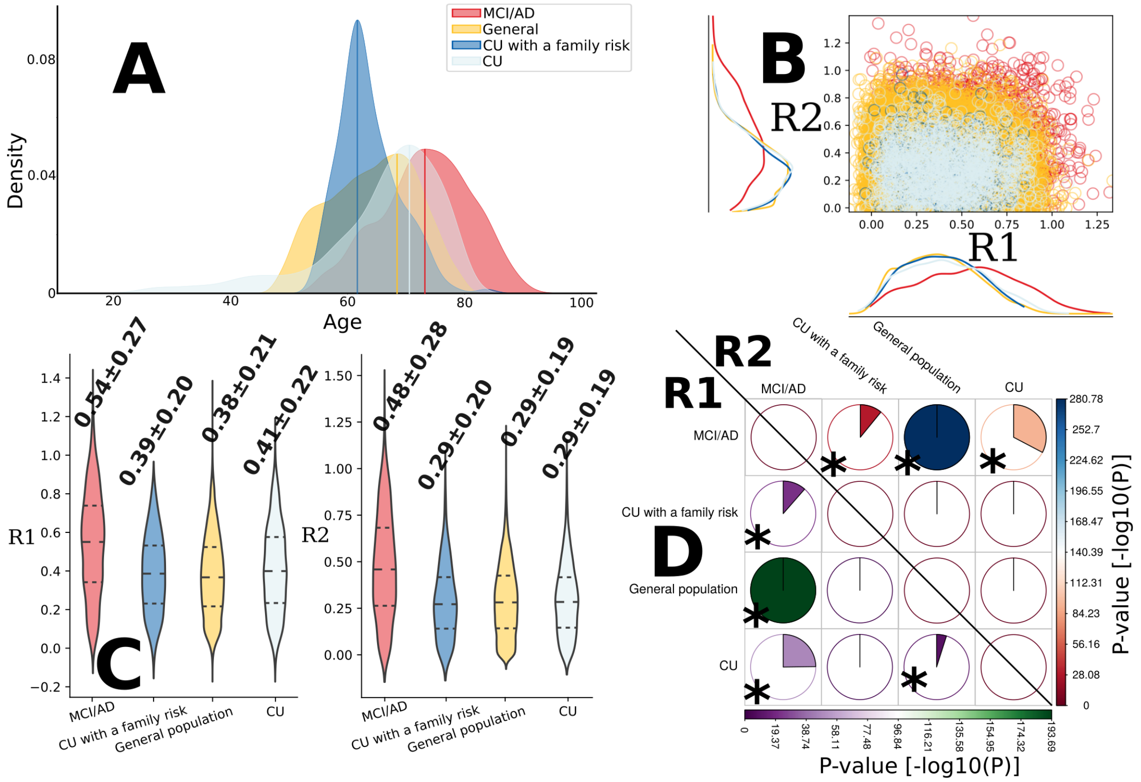


**A**) Four different populations were used in this study: *i*) the *MCI/AD population* from ADNI[26] and BLSA,[27] which represented the symptomatic population (age distribution: 73.45±7.69); *ii*) the *general population* from UKBB,[28] which included participants with high AD risks based on family history of AD (i.e., proxy-AD, *N*=10,189 out of 39,575) and many other disease diagnoses (age distribution: 64.12±7.54); *iii*) the *cognitively unimpaired (CU) population* from ADNI[26] and BLSA,[27] which endorsed the normative aging elders without any cognitive decline at baseline (age distribution: 65.75±10.90); *iv*) the *CU with a family risk population*, which recruited 343 CU participants that had a first-relative family history of AD diagnosis (age distribution: 63.62±5.05). Refer to Method 1 in the main manuscript for more details. A kernel density estimate plot visualizes the age distribution of each population. **B**) The scatter plot for the R1 and R2 dimensions in the four populations is presented, together with the kernel density estimate plot for each population. The two dimensions are highly expressed in the MCI/AD population and gradually less manifested in the three asymptomatic populations: the general population, the CU population, and the CU with a family history population. Of note, some participants showed negative values for the two neuroanatomical dimensions. These participants did not express much for this dimension and can be interpreted as super normal individuals compared to the MCI/AD patients from ADNI in the training sample. **C**) The violin plots show the distribution of the R1 and R2 dimensions of Surreal-GAN in the four populations. **D**) The pie plots show the statistically significant difference (denoted with *) of the R1 and R2 dimensions in each pair of populations after Bonferroni correction [>(-$\log_{10}$(0.05/2/4)=2.20). The left-lower panel shows the results for R1, and the right-upper panel displays the results for R2.

**eFigure 5: Longitudinal conversion of MCI to AD and between R1- and R2-dominant groups in the MCI/AD population**

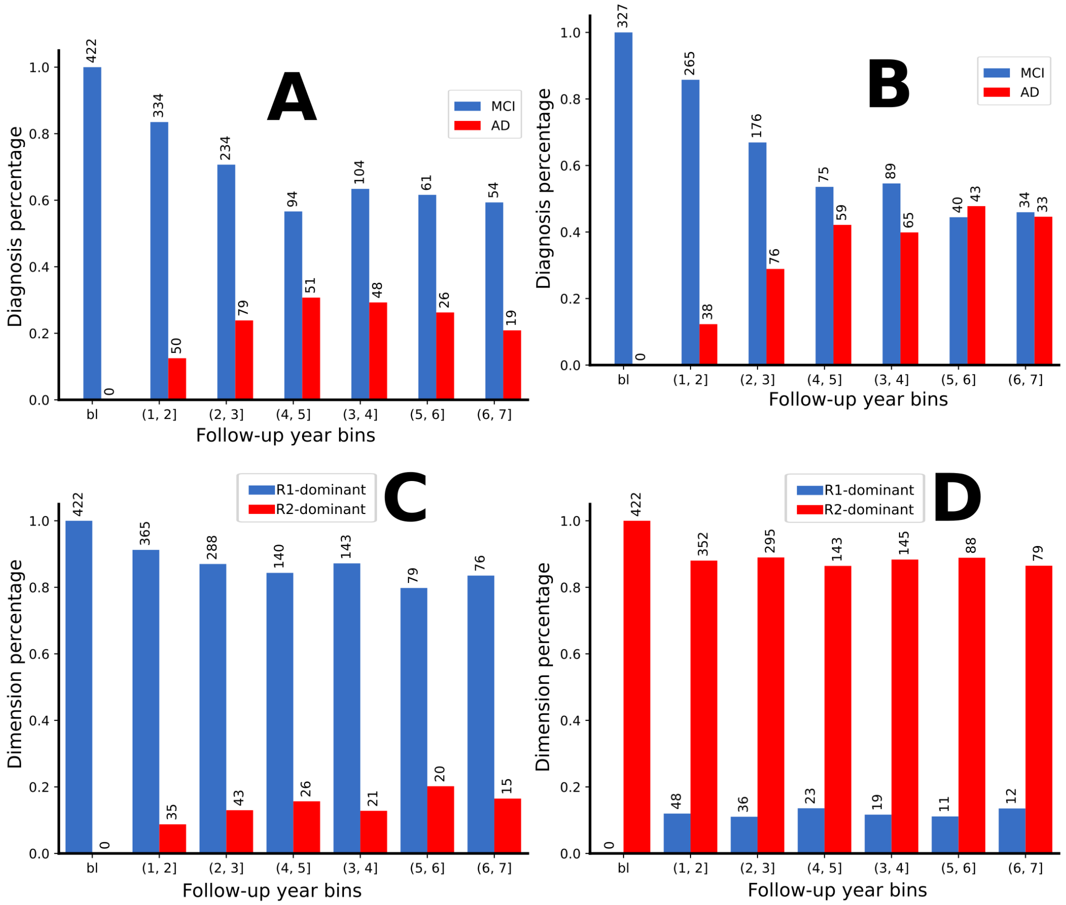


At baseline, MCI participants transition to AD within a 6-year follow-up period, as assessed using all available longitudinal scans (>6 years resulted in a small number of participants) for both the R1-dominant group (**A**) and the R2-dominant group (**B**). Likewise, MCI participants at baseline transition from the R1-dominant group to the R2-dominant group (**C**) and in the reverse direction (**D**). The percentage of each diagnosis and dimension group is shown on the y-axis; the number of participants in each diagnosis and dimension group is also shown.

**eFigure 6: PRS plots at baseline for the MCI/AD population**

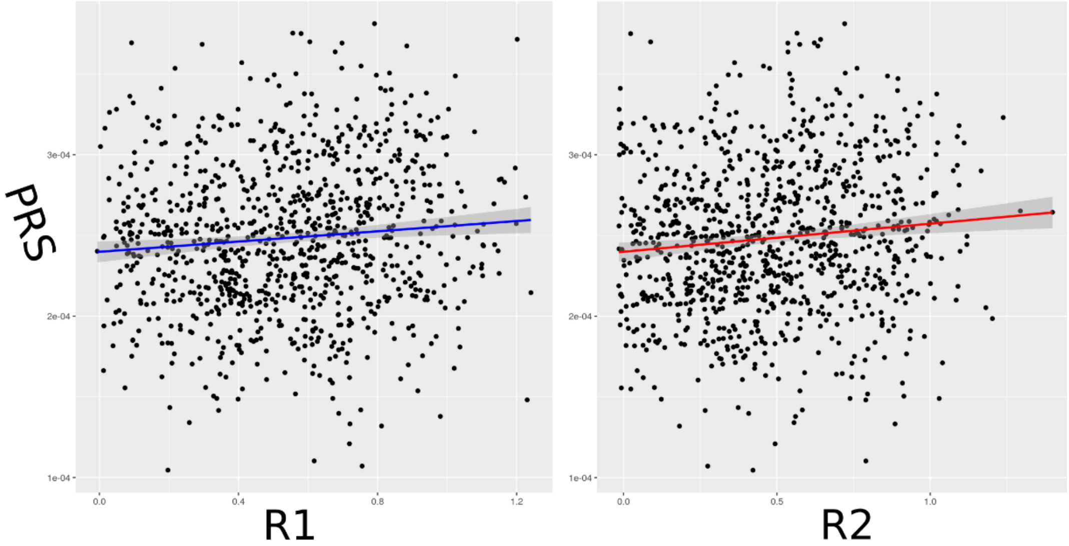


We fit a linear regression model to associate the PRS with the two dimensions (**Method 6F** in the main manuscript). The results showed a slightly stronger positive association with the R2 dimension [$r$=0.11, -$\log_{10}$(P-value)=3.14] than with the R1 dimension [$r$=0.09, -$\log_{10}$(P-value)=2.31].

**eFigure 7: QQ plots for the baseline GWAS for the MCI/AD population**

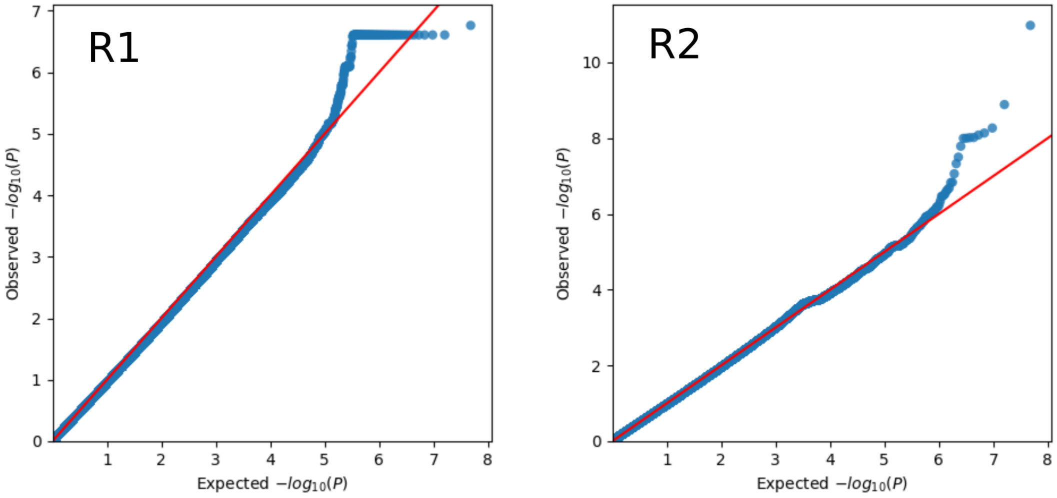


The two GWASs are overall under-powered in the MCI/AD population.

**eFigure 8: Manhattan and QQ plots for the longitudinal GWAS for the general population**

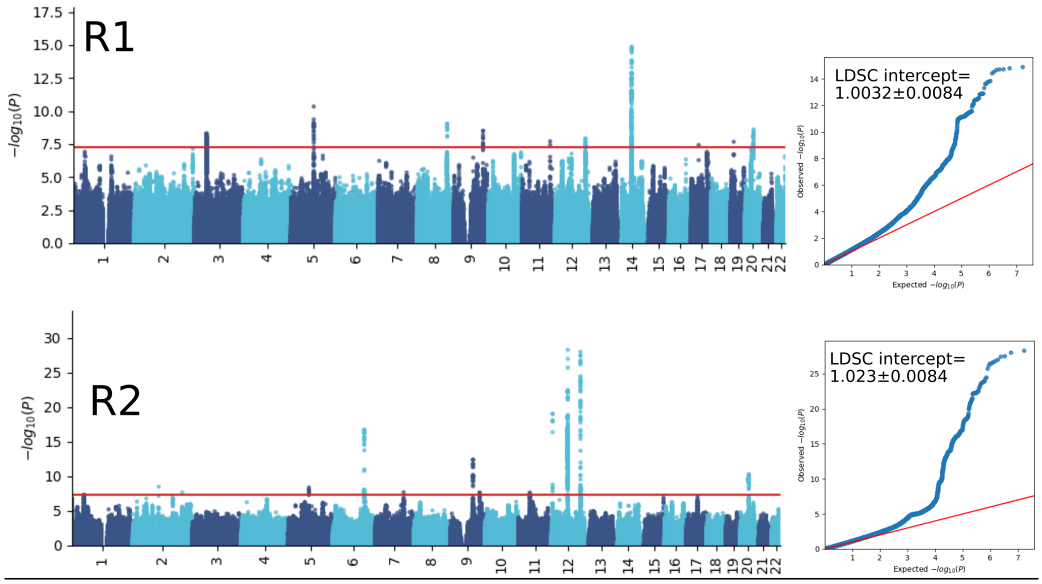


We present the LDSC intercepts for the two GWASs.

**eFigure 9: Tissue specificity analyses for the prioritized genes in the two dimensions in the general population**

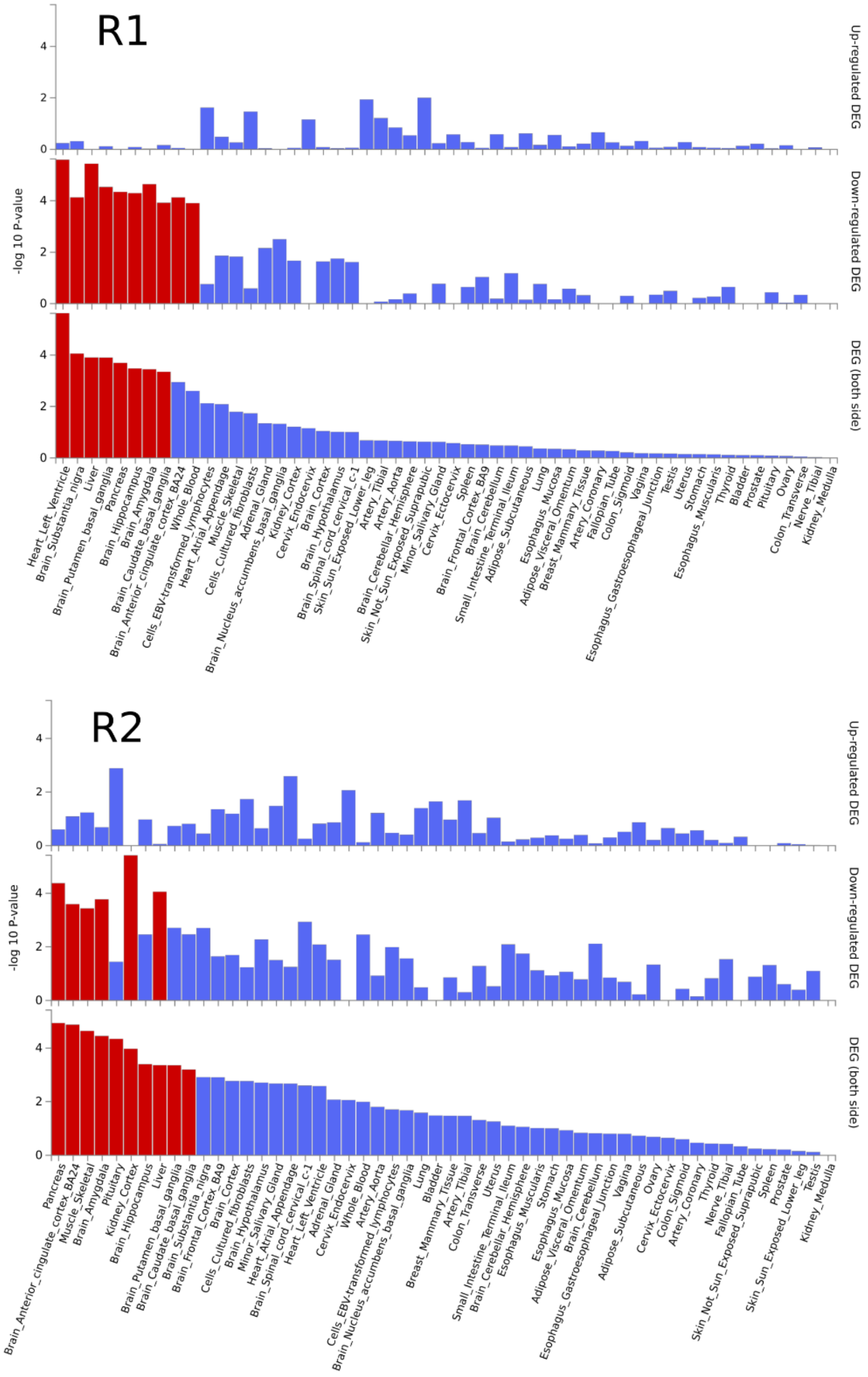


We performed a prioritized tissue specificity analysis using the mapped/annotated genes. Significantly enriched differentially expressed gene sets (DEG) are highlighted in red. The *GENE2FUNC*[17] pipeline from FUMA was performed to examine the overrepresentation of prioritized genes in pre-defined DEGs (up-regulated, down-regulated, and both-side DEGs) from different gene expression data. The input genes (Fig. 2C) were tested against each DEG using the hypergeometric test.

**eFigure 10: Singe gene expression heat maps for the tissue specificity analyses in the general population**

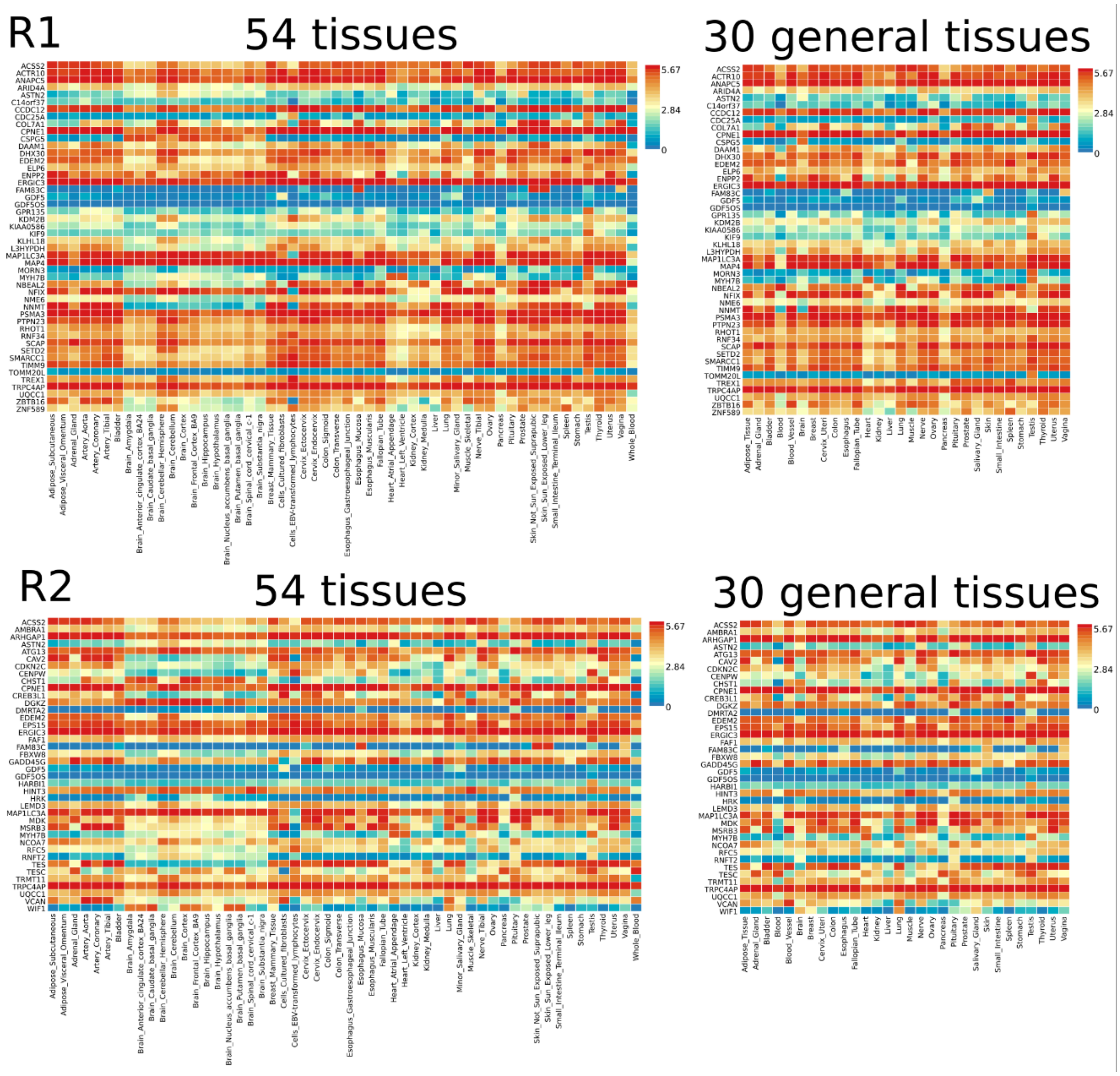


The gene-level heatmap displays the average expression per label ($log_2$ transformed) using data from GTEx v8 54 tissue types (left) and 30 general tissue types (right). Genes and tissue/organs are ordered in alphabetical order.

**eFigure 11: The longitudinal rate of change of R1 and R2 neuroanatomical dimensions of brain atrophy in the MCI/AD population.**

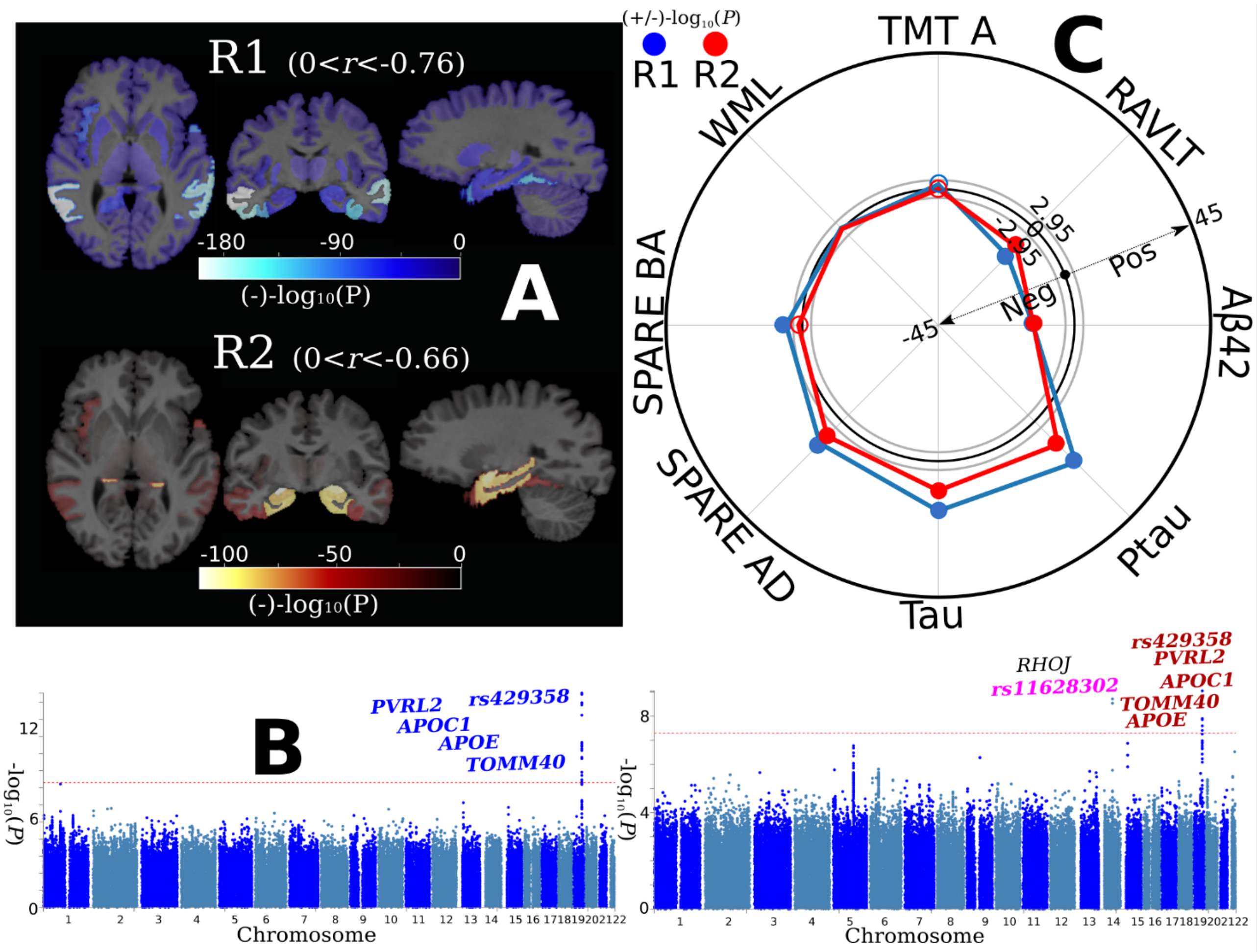


**A)** Longitudinal brain association studies show that the R1 dimension exhibits widespread brain atrophy, whereas the R2 dimension displays focal medial temporal lobe atrophy. We first derived the rate of change of the 119 GM ROIs and the R1 and R2 dimensions using a linear mixed effect model; a linear regression model was then fit to the rate of change of the ROIs, R1, and R2 to derive the beta value of each ROI. A negative value denotes longitudinal brain atrophy with a negative coefficient of the rate of change in the linear regression model. The range of $r$ for each dimension is also shown. Of note, the sample size ($N$) for R1 and R2 is the same for each ROI. **B)** Genome-wide association studies demonstrate that the R1 and R2 dimensions are both associated with variants related to *APOE* genes (genome-wide P-value threshold with the red line: $-\log_{10}$(P-value) > 7.30). We associated each common variant with R1 and R2 using the whole-genome sequencing data from ADNI. Gene annotations were performed via positional, expression quantitative trait loci, and chromatin interaction mappings using FUMA.[17] We then manually queried whether they were previously associated with AD-related traits in the GWAS Catalog.[29] Blue and Red-colored loci/genes indicate variants associated with AD-related traits in previous literature for R1 and R2, respectively. **C)** Clinical association studies show that the R1 and R2 dimensions are associated with SPARE-AD[30], the CSF level of tau, ptau, and amyloid.

**eTable 1: Brain association studies for the MCI/AD population from ADNI and BLSA.** Each ROI was fit with a linear regression model by controlling covariates. $-\log_{10}$(P-value) and Pearson's correlation (*r*) are reported. For the P-values, we also added the sign of the ROI's beta coefficient values to indicate the effect's direction. The threshold for Bonferroni corrected $-\log_{10}$(P-value) is > 3.38. For the *r* values, the ROIs that did not survive the multiple comparisons or showed a positive *r* (negative *r* means brain atrophy) are assigned a value of 0 for visualization purposes.

| ROI | R1: sign[$-\log_{10}$(P-value)] | R2: sign[$-\log_{10}$(P-value)] | R1: *r* | R2: *r* |
|---|---|---|---|---|
| Right Accumbens Area | -7.50571 | -18.5683 | -0.13385 | -0.22157 |
| Left Accumbens Area | -4.57493 | -16.6058 | -0.10096 | -0.21176 |
| Right Amygdala | -12.1396 | -281.515 | -0.17085 | -0.65027 |
| Left Amygdala | -13.421 | -262.174 | -0.17854 | -0.63577 |
| Right Caudate | 4.47843 | 0.06662 | 0.03963 | 0 |
| Left Caudate | 3.80988 | 0.01493 | 0.0358 | 0 |
| Right Cerebellum Exterior | -1.42046 | -1.43982 | 0 | 0 |
| Left Cerebellum Exterior | -0.89444 | -1.43875 | 0 | 0 |
| Right Hippocampus | -7.10095 | -273.254 | -0.12845 | -0.65086 |
| Left Hippocampus | -4.85907 | -286.073 | -0.10114 | -0.66999 |
| Right Pallidum | 0.45787 | -0.44775 | 0 | 0 |
| Left Pallidum | 0.90213 | -0.62152 | 0 | 0 |
| Right Putamen | -1.32989 | -1.3305 | 0 | 0 |
| Left Putamen | -1.08232 | -0.87145 | 0 | 0 |
| Right Thalamus Proper | -5.56006 | -14.8387 | -0.1137 | -0.16236 |
| Left Thalamus Proper | -4.7225 | -11.6842 | -0.10534 | -0.14483 |
| Cerebellar Vermal Lobules I-V | -2.35113 | -0.08627 | 0 | 0 |
| Cerebellar Vermal Lobules VI-VII | 1.2309 | 0.19721 | 0 | 0 |
| Cerebellar Vermal Lobules VIII-X | 1.01513 | -1.28001 | 0 | 0 |
| Left Basal Forebrain | -3.13748 | -31.1551 | 0 | -0.30623 |
| Right Basal Forebrain | -0.07585 | -18.4305 | 0 | -0.24177 |
| Right ACgG anterior cingulate gyrus | -2.78765 | -5.35387 | 0 | -0.12467 |
| Left ACgG anterior cingulate gyrus | -8.48968 | -2.56191 | -0.15256 | 0 |
| Right AIns anterior insula | -26.1772 | -16.5433 | -0.23749 | -0.18867 |
| Left AIns anterior insula | -29.959 | -21.3573 | -0.25152 | -0.21728 |
| Right AOrG anterior orbital gyrus | -4.39974 | 0.14837 | -0.12046 | 0 |
| Left AOrG anterior orbital gyrus | -4.6851 | -0.18991 | -0.1193 | 0 |
| Right AnG angular gyrus | -93.8132 | -0.65332 | -0.46814 | 0 |
| Left AnG angular gyrus | -72.644 | -1.2325 | -0.42036 | 0 |
| Right Calc calcarine cortex | 0.13368 | 0.53457 | 0 | 0 |
| Left Calc calcarine cortex | 0.14284 | 1.57312 | 0 | 0 |
| Right CO central operculum | -40.5874 | -0.15535 | -0.28393 | 0 |
| Left CO central operculum | -53.6712 | -0.88026 | -0.32856 | 0 |
| Right Cun cuneus | -2.36823 | -0.23906 | 0 | 0 |
| Left Cun cuneus | -1.68046 | -0.22262 | 0 | 0 |
| Right Ent entorhinal area | -11.3956 | -212.355 | -0.16588 | -0.57772 |
| Left Ent entorhinal area | -14.2329 | -234.601 | -0.17832 | -0.63567 |

| | | | | |
|---|---|---|---|---|
| Right FO frontal operculum | -25.7413 | -0.60814 | -0.24631 | 0 |
| Left FO frontal operculum | -21.6485 | -1.82348 | -0.21898 | 0 |
| Right FRP frontal pole | -3.30212 | 1.53437 | 0 | 0 |
| Left FRP frontal pole | -8.63073 | -0.61843 | -0.16264 | 0 |
| Right FuG fusiform gyrus | -47.9404 | -26.9014 | -0.30709 | -0.21163 |
| Left FuG fusiform gyrus | -50.7759 | -26.4785 | -0.31803 | -0.22676 |
| Right GRe gyrus rectus | -2.45628 | -3.16739 | 0 | 0 |
| Left GRe gyrus rectus | 0.0541 | -2.01584 | 0 | 0 |
| Right IOG inferior occipital gyrus | -22.9084 | 0.04341 | -0.24389 | 0 |
| Left IOG inferior occipital gyrus | -28.3487 | 0.1735 | -0.26282 | 0 |
| Right ITG inferior temporal gyrus | -128.732 | -18.3864 | -0.46735 | -0.16197 |
| Left ITG inferior temporal gyrus | -120.071 | -20.2061 | -0.46359 | -0.17083 |
| Right LiG lingual gyrus | -6.49459 | -1.78258 | -0.14713 | 0 |
| Left LiG lingual gyrus | -2.5011 | -0.47708 | 0 | 0 |
| Right LOrG lateral orbital gyrus | 0.47646 | 3.80038 | 0 | 0.08938 |
| Left LOrG lateral orbital gyrus | -0.16276 | 0.35955 | 0 | 0 |
| Right MCgG middle cingulate gyrus | -4.18551 | -0.6196 | -0.12514 | 0 |
| Left MCgG middle cingulate gyrus | -3.06066 | -0.00227 | 0 | 0 |
| Right MFC medial frontal cortex | -4.06739 | -6.21943 | -0.0963 | -0.1428 |
| Left MFC medial frontal cortex | -0.70819 | -2.32881 | 0 | 0 |
| Right MFG middle frontal gyrus | -23.0666 | 6.25053 | -0.23259 | 0.09546 |
| Left MFG middle frontal gyrus | -28.2634 | 4.22507 | -0.25529 | 0.07449 |
| Right MOG middle occipital gyrus | -29.2134 | -0.6177 | -0.2844 | 0 |
| Left MOG middle occipital gyrus | -36.8055 | -1.00391 | -0.31201 | 0 |
| Right MOrG medial orbital gyrus | -6.72452 | -0.67406 | -0.15059 | 0 |
| Left MOrG medial orbital gyrus | -0.78826 | 0.0603 | 0 | 0 |
| Right MPoG postcentral gyrus medial segment | -0.22712 | 1.52575 | 0 | 0 |
| Left MPoG postcentral gyrus medial segment | 0.07641 | 1.87733 | 0 | 0 |
| Right MPrG precentral gyrus medial segment | -0.12598 | 1.43652 | 0 | 0 |
| Left MPrG precentral gyrus medial segment | -0.70782 | 4.12148 | 0 | 0.08423 |
| Right MSFG superior frontal gyrus medial segment | -11.9297 | 0.46343 | -0.17345 | 0 |
| Left MSFG superior frontal gyrus medial segment | -13.114 | -0.31839 | -0.17525 | 0 |
| Right MTG middle temporal gyrus | -155.837 | -15.2718 | -0.51863 | -0.16543 |
| Left MTG middle temporal gyrus | -149.597 | -15.5284 | -0.5215 | -0.17027 |
| Right OCP occipital pole | 4.31229 | 0.64664 | 0.036 | 0 |
| Left OCP occipital pole | 0.93506 | 0.10407 | 0 | 0 |
| Right OFuG occipital fusiform gyrus | -8.06137 | 0.24056 | -0.15547 | 0 |

| | | | | |
|---|---|---|---|---|
| Left OFuG occipital fusiform gyrus | -27.0683 | -3.09207 | -0.2645 | 0 |
| Right OpIFG opercular part of the inferior frontal gyrus | -17.399 | -0.07107 | -0.20488 | 0 |
| Left OpIFG opercular part of the inferior frontal gyrus | -30.847 | 0.40158 | -0.26572 | 0 |
| Right OrIFG orbital part of the inferior frontal gyrus | -5.11808 | 1.5731 | -0.11731 | 0 |
| Left OrIFG orbital part of the inferior frontal gyrus | -3.7895 | 2.03442 | -0.09809 | 0 |
| Right PCgG posterior cingulate gyrus | -32.6246 | -2.30486 | -0.28065 | 0 |
| Left PCgG posterior cingulate gyrus | -37.6512 | -4.94491 | -0.29181 | -0.07176 |
| Right PCu precuneus | -62.8395 | 0.22134 | -0.36339 | 0 |
| Left PCu precuneus | -47.6593 | -0.36123 | -0.32947 | 0 |
| Right PHG parahippocampal gyrus | -19.1909 | -189.939 | -0.20697 | -0.5469 |
| Left PHG parahippocampal gyrus | -19.7229 | -208.439 | -0.20401 | -0.57164 |
| Right PIns posterior insula | -49.5689 | -9.65035 | -0.3065 | -0.15683 |
| Left PIns posterior insula | -60.8608 | -6.47558 | -0.33883 | -0.14154 |
| Right PO parietal operculum | -18.1403 | 0.86838 | -0.2199 | 0 |
| Left PO parietal operculum | -21.4814 | 0.2996 | -0.23754 | 0 |
| Right PoG postcentral gyrus | -16.0393 | 2.99274 | -0.2013 | 0 |
| Left PoG postcentral gyrus | -16.6346 | 2.15175 | -0.20579 | 0 |
| Right POrG posterior orbital gyrus | -0.91914 | -2.81108 | 0 | 0 |
| Left POrG posterior orbital gyrus | -0.14594 | -2.28165 | 0 | 0 |
| Right PP planum polare | -29.4866 | -8.91216 | -0.24271 | -0.15817 |
| Left PP planum polare | -16.8675 | -7.19882 | -0.18734 | -0.15264 |
| Right PrG precentral gyrus | -12.9721 | 9.52306 | -0.17726 | 0.1078 |
| Left PrG precentral gyrus | -15.5279 | 5.81795 | -0.19317 | 0.08159 |
| Right PT planum temporale | -23.4547 | -0.65338 | -0.23324 | 0 |
| Left PT planum temporale | -28.7539 | -0.77124 | -0.26384 | 0 |
| Right SCA subcallosal area | -0.05839 | -3.30599 | 0 | 0 |
| Left SCA subcallosal area | -0.04662 | -1.18346 | 0 | 0 |
| Right SFG superior frontal gyrus | -20.3875 | 1.36576 | -0.22634 | 0 |
| Left SFG superior frontal gyrus | -21.7389 | 1.42391 | -0.23471 | 0 |
| Right SMC supplementary motor cortex | -3.43682 | -0.00728 | -0.09812 | 0 |
| Left SMC supplementary motor cortex | -1.81719 | -0.50428 | 0 | 0 |
| Right SMG supramarginal gyrus | -89.0207 | -0.09531 | -0.45635 | 0 |
| Left SMG supramarginal gyrus | -93.6886 | 1.18687 | -0.47475 | 0 |
| Right SOG superior occipital gyrus | -13.1905 | 0.37282 | -0.18002 | 0 |
| Left SOG superior occipital gyrus | -7.89322 | 0.20519 | -0.13827 | 0 |

| | | | | |
|---|---|---|---|---|
| Right SPL superior parietal lobule | -35.647 | 0.72415 | -0.29015 | 0 |
| Left SPL superior parietal lobule | -36.7243 | 1.40962 | -0.2972 | 0 |
| Right STG superior temporal gyrus | -50.1012 | -4.34522 | -0.33617 | -0.10192 |
| Left STG superior temporal gyrus | -58.679 | -6.32759 | -0.36681 | -0.12603 |
| Right TMP temporal pole | -42.0432 | -87.8416 | -0.30518 | -0.40385 |
| Left TMP temporal pole | -44.3756 | -74.3941 | -0.31169 | -0.38748 |
| Right TrIFG triangular part of the inferior frontal gyrus | -0.63125 | -0.26408 | 0 | 0 |
| Left TrIFG triangular part of the inferior frontal gyrus | -4.62617 | 0.29648 | -0.11686 | 0 |
| Right TTG transverse temporal gyrus | -1.92176 | -0.36857 | 0 | 0 |
| Left TTG transverse temporal gyrus | -4.67912 | -0.00132 | -0.11116 | 0 |

**eTable 2: Identified genomic loci and mapped genes for baseline GWAS in the MCI/AD population**

| Locus (lead SNP) | $-\log_{10}$(P-value) | Chromosome | Mapped genes |
|---|---|---|---|
| rs429358 | 10.98 | 19 | *APOE, PVRL2, APOC1, TOMM40* |

**eTable 3: Clinical association studies for the MCI/AD population from ADNI and BLSA.** Each clinical variable/biomarker was fit with a linear regression model by controlling covariates. $-\log_{10}$(P-value) and the coefficient of each variable/biomarker (beta) are reported. For the P-values, we added the sign of the variable's beta coefficient values to indicate the effect's direction. The threshold for Bonferroni corrected $-\log_{10}$(P-value) is > 2.95.

| Variable | R1: sign[$-\log_{10}$(P-value)] | R2: sign[$-\log_{10}$(P-value)] |
|---|---|---|
| FAQ_Total | 27.553 | 28.8057 |
| CDR_Global | 19.4197 | 14.7913 |
| CDR_SOB | 25.1713 | 25.8645 |
| NPIQ_Total | 1.98106 | 0.82936 |
| DSST | -9.43994 | -0.04787 |
| TMT_A | 41.8293 | 0.27449 |
| TMT_B | 43.8428 | 2.95569 |
| MMSE | -27.793 | -29.9245 |
| Digit_Span_Backward | -5.19651 | -0.29222 |
| Digit_Span_Forward | -1.232 | -0.87773 |
| MOCA | -19.1169 | -12.0298 |
| BNT | -14.9887 | -16.0565 |
| RAVLT | -24.1567 | -38.206 |
| RAVLT_IM | -27.8745 | -30.1843 |
| RAVLT_Short | -13.0604 | -38.1494 |
| RAVLT_Long | -13.2972 | -37.6006 |
| LM_DEL_A | -21.6429 | -60.621 |
| LM_IM_A | -28.0603 | -31.5233 |
| ANI_Fluency | -27.8458 | -10.4616 |
| VEG_Fluency | -12.1705 | -4.17084 |
| TS_RATIO | 0.08455 | 0.82137 |
| TS_RATIO_ADJ | 0.09716 | 0.64491 |
| BNT_percent | -14.9887 | -16.0565 |
| Cholesterol | 0.10488 | 0.30897 |
| Diastole | 0.53221 | -0.23115 |
| Glucose | 0.02649 | -0.56589 |
| Systole | 4.03676 | -0.40777 |
| Triglycerides | -0.70672 | 0.10056 |
| ABETA42_ADNI | -16.6712 | -9.64753 |
| PTAU | 5.69056 | 6.12353 |
| TAU | 4.7303 | 7.25391 |
| Diabetes | -0.00052 | 1.41286 |
| Hypertension | 2.00269 | 0.027 |
| Diagnosis_Depression | -0.44864 | 0.047 |
| Alcoholic | 0.89498 | 0.32819 |
| Family_History_Dementia | -1.84702 | -0.25455 |
| Ethnicity | 0.80007 | 0.04149 |
| Height | -0.57955 | 0.70593 |
| BMI | -0.97997 | -7.32202 |
| Weight | -1.57751 | -5.66086 |
| SPARE_AD | 109.59 | 201.144 |
| SPARE_BA | 54.6991 | 8.33545 |
| AV45_PET_ADNI | 8.33865 | 4.19236 |
| FDG | -52.5638 | -10.3312 |
| WMLS_Volume_701 | 6.1794 | 1.29141 |

**eTable 4: Brain association studies for the general population from UKBB.** Each ROI was fit with a linear regression model by controlling covariates. $-\log_{10}$(P-value) and Pearson's correlation ($r$) are reported. For the P-values, we also added the sign of the ROI's beta coefficient values to indicate the effect's direction. The threshold for Bonferroni corrected $-\log_{10}$(P-value) is > 3.38. For the $r$ values, the ROIs that did not survive the multiple comparisons or showed a positive $r$ (negative $r$ means brain atrophy) are assigned a value of 0 for visualization purposes.

| ROI | R1: sign[$-\log_{10}$(P-value)] | R2: sign[$-\log_{10}$(P-value)] | R1: $r$ | R2: $r$ |
|---|---|---|---|---|
| Right Accumbens Area | 1.41659 | -182.266 | 0 | -0.0492 |
| Left Accumbens Area | 1.61428 | -167.936 | 0 | -0.03835 |
| Right Amygdala | -98.726 | -330 | -0.15118 | -0.33719 |
| Left Amygdala | -56.4488 | -330 | -0.12555 | -0.31386 |
| Right Caudate | 300 | -44.3821 | 0.11226 | -0.02326 |
| Left Caudate | 300 | -37.3245 | 0.10616 | -0.00926 |
| Right Cerebellum Exterior | -9.33216 | -151.136 | -0.08723 | 0.00104 |
| Left Cerebellum Exterior | -12.3741 | -116.615 | -0.09151 | 0.0084 |
| Right Hippocampus | 0.12934 | -330 | 0 | -0.36498 |
| Left Hippocampus | 1.61488 | -330 | 0 | -0.34206 |
| Right Pallidum | 22.3496 | -27.02 | -0.038 | 0.0343 |
| Left Pallidum | 21.293 | -47.475 | -0.02817 | 0.01584 |
| Right Putamen | -0.82029 | -69.6543 | 0 | 0.00366 |
| Left Putamen | -0.3018 | -59.1727 | 0 | 0.00934 |
| Right Thalamus Proper | -28.9526 | -87.2289 | -0.12938 | 0.04023 |
| Left Thalamus Proper | -33.4427 | -92.0801 | -0.13339 | 0.04002 |
| Cerebellar Vermal Lobules I-V | -29.5876 | -7.76566 | -0.10368 | 0.03997 |
| Cerebellar Vermal Lobules VI-VII | 0.80143 | -21.3033 | 0 | -0.00421 |
| Cerebellar Vermal Lobules VIII-X | -3.851 | -87.7558 | -0.05945 | -0.02211 |
| Left Basal Forebrain | 137.211 | -64.3165 | 0.06201 | -0.03614 |
| Right Basal Forebrain | 104.834 | -110.189 | 0.04623 | -0.05927 |
| Right ACgG anterior cingulate gyrus | -0.21265 | -5.72052 | 0 | 0.03231 |
| Left ACgG anterior cingulate gyrus | -10.4907 | -5.89634 | -0.1117 | 0.04079 |
| Right AIns anterior insula | -90.4988 | -112.325 | -0.15479 | -0.00513 |
| Left AIns anterior insula | -87.4626 | -84.7404 | -0.15327 | 0.00519 |
| Right AOrG anterior orbital gyrus | -3.62996 | 142.979 | -0.08145 | 0.17176 |
| Left AOrG anterior orbital gyrus | -5.75963 | 70.2007 | -0.06233 | 0.13786 |
| Right AnG angular gyrus | -300 | 13.7095 | -0.37557 | 0.10415 |
| Left AnG angular gyrus | -300 | 32.058 | -0.3504 | 0.11536 |
| Right Calc calcarine cortex | 14.0495 | 84.0712 | -0.02481 | 0.14709 |
| Left Calc calcarine cortex | 22.6608 | 92.9419 | -0.00831 | 0.14631 |
| Right CO central operculum | -300 | 66.8329 | -0.25532 | 0.14862 |
| Left CO central operculum | -300 | 15.5315 | -0.28549 | 0.11619 |
| Right Cun cuneus | -1.40722 | 25.3579 | 0 | 0.11906 |
| Left Cun cuneus | 0.01067 | 35.5432 | 0 | 0.12482 |
| Right Ent entorhinal area | -15.7387 | -330 | -0.07315 | -0.33158 |
| Left Ent entorhinal area | -37.3154 | -330 | -0.09629 | -0.38061 |
| Right FO frontal operculum | -283.5 | 2.60188 | -0.20745 | 0 |

| | | | | |
|---|---|---|---|---|
| Left FO frontal operculum | -174.006 | 67.8112 | -0.18019 | 0.14426 |
| Right FRP frontal pole | -12.934 | 178.43 | -0.11962 | 0.1829 |
| Left FRP frontal pole | -43.9357 | 73.5255 | -0.13305 | 0.1402 |
| Right FuG fusiform gyrus | -198.547 | -36.8795 | -0.17519 | 0.03427 |
| Left FuG fusiform gyrus | -300 | -18.9938 | -0.21721 | 0.04813 |
| Right GRe gyrus rectus | -3.66473 | 9.10609 | -0.08331 | 0.09133 |
| Left GRe gyrus rectus | 0.7205 | 15.698 | 0 | 0.09976 |
| Right IOG inferior occipital gyrus | -15.7166 | 5.87919 | -0.10042 | 0.08228 |
| Left IOG inferior occipital gyrus | -7.20403 | 113.876 | -0.10013 | 0.14714 |
| Right ITG inferior temporal gyrus | -300 | 0.85369 | -0.32453 | 0 |
| Left ITG inferior temporal gyrus | -300 | 15.0401 | -0.3341 | 0.104 |
| Right LiG lingual gyrus | -53.927 | -20.8439 | -0.13147 | 0.04954 |
| Left LiG lingual gyrus | -6.76542 | 26.077 | -0.10231 | 0.11848 |
| Right LOrG lateral orbital gyrus | 18.2409 | 132.453 | -0.0165 | 0.1532 |
| Left LOrG lateral orbital gyrus | 36.4951 | 210.898 | -0.00863 | 0.19322 |
| Right MCgG middle cingulate gyrus | -3.90798 | 25.1491 | -0.09126 | 0.10476 |
| Left MCgG middle cingulate gyrus | -8.4155 | 15.4716 | -0.0963 | 0.09492 |
| Right MFC medial frontal cortex | 1.78702 | -3.97811 | 0 | 0.04703 |
| Left MFC medial frontal cortex | 21.2943 | 2.90145 | -0.02022 | 0 |
| Right MFG middle frontal gyrus | -129.233 | 279.61 | -0.17922 | 0.21173 |
| Left MFG middle frontal gyrus | -109.494 | 330 | -0.17173 | 0.22885 |
| Right MOG middle occipital gyrus | -149.532 | 88.1747 | -0.1873 | 0.14692 |
| Left MOG middle occipital gyrus | -300 | 18.7585 | -0.26727 | 0.10437 |
| Right MOrG medial orbital gyrus | 15.4308 | 9.37965 | -0.06111 | 0.09879 |
| Left MOrG medial orbital gyrus | 106.518 | 15.6631 | 0.00145 | 0.0948 |
| Right MPoG postcentral gyrus medial segment | 18.4511 | 106.131 | -0.00525 | 0.14145 |
| Left MPoG postcentral gyrus medial segment | 0.94506 | 79.6395 | 0 | 0.13066 |
| Right MPrG precentral gyrus medial segment | 18.3772 | 193.817 | -0.01673 | 0.19332 |
| Left MPrG precentral gyrus medial segment | 69.2107 | 147.742 | 0.03316 | 0.16704 |
| Right MSFG superior frontal gyrus medial segment | -76.67 | 244.578 | -0.15288 | 0.20867 |
| Left MSFG superior frontal gyrus medial segment | -91.3879 | 168.41 | -0.15268 | 0.19107 |
| Right MTG middle temporal gyrus | -300 | -20.9451 | -0.37466 | 0.05852 |
| Left MTG middle temporal gyrus | -300 | -21.69 | -0.39366 | 0.0523 |
| Right OCP occipital pole | 197.625 | 84.9317 | 0.0602 | 0.13398 |
| Left OCP occipital pole | 255.843 | 211.639 | 0.06741 | 0.1804 |
| Right OFuG occipital fusiform gyrus | -12.0126 | 94.2742 | -0.0894 | 0.14724 |
| Left OFuG occipital fusiform gyrus | -101.671 | -3.81391 | -0.14763 | 0.05335 |

| | | | | |
|---|---|---|---|---|
| Right OpIFG opercular part of the inferior frontal gyrus | -273.101 | 91.9958 | -0.20386 | 0.15424 |
| Left OpIFG opercular part of the inferior frontal gyrus | -243.767 | 180.399 | -0.20055 | 0.19079 |
| Right OrIFG orbital part of the inferior frontal gyrus | -2.27412 | 22.8358 | 0 | 0.09156 |
| Left OrIFG orbital part of the inferior frontal gyrus | -0.24746 | 58.0824 | 0 | 0.12043 |
| Right PCgG posterior cingulate gyrus | -209.592 | -11.9412 | -0.19378 | 0.0483 |
| Left PCgG posterior cingulate gyrus | -262.46 | -34.3738 | -0.20553 | 0.03161 |
| Right PCu precuneus | -300 | 54.9645 | -0.29166 | 0.14355 |
| Left PCu precuneus | -300 | 40.902 | -0.27177 | 0.13637 |
| Right PHG parahippocampal gyrus | -173.097 | -330 | -0.16496 | -0.34984 |
| Left PHG parahippocampal gyrus | -198.601 | -330 | -0.17802 | -0.37505 |
| Right PIns posterior insula | -266.472 | -23.7998 | -0.21771 | 0.04153 |
| Left PIns posterior insula | -300 | -89.6578 | -0.23383 | -0.00766 |
| Right PO parietal operculum | -66.7351 | 66.8369 | -0.14711 | 0.13074 |
| Left PO parietal operculum | -102.229 | 85.3908 | -0.1649 | 0.15125 |
| Right PoG postcentral gyrus | -282.817 | 120.309 | -0.2271 | 0.16294 |
| Left PoG postcentral gyrus | -270.006 | 78.0722 | -0.22169 | 0.14354 |
| Right POrG posterior orbital gyrus | 78.6402 | 1.4835 | 0.00852 | 0 |
| Left POrG posterior orbital gyrus | 45.1416 | 1.80514 | -0.00667 | 0 |
| Right PP planum polare | -71.6443 | -43.1669 | -0.14793 | 0.00847 |
| Left PP planum polare | -34.4771 | -11.488 | -0.12538 | 0.03474 |
| Right PrG precentral gyrus | -46.192 | 330 | -0.13865 | 0.2993 |
| Left PrG precentral gyrus | -45.3155 | 330 | -0.13484 | 0.28293 |
| Right PT planum temporale | -120.698 | 115.443 | -0.16067 | 0.16964 |
| Left PT planum temporale | -273.535 | 9.16214 | -0.20004 | 0.10329 |
| Right SCA subcallosal area | 65.4707 | -8.55722 | -0.00285 | 0.02647 |
| Left SCA subcallosal area | 71.5098 | -12.2986 | 0.00515 | 0.0165 |
| Right SFG superior frontal gyrus | -36.5317 | 274.089 | -0.13275 | 0.21365 |
| Left SFG superior frontal gyrus | -53.0535 | 189.783 | -0.13843 | 0.19185 |
| Right SMC supplementary motor cortex | -0.03372 | 9.90833 | 0 | 0.09648 |
| Left SMC supplementary motor cortex | 2.78577 | 46.7257 | 0 | 0.13199 |
| Right SMG supramarginal gyrus | -300 | 48.1282 | -0.3503 | 0.12322 |
| Left SMG supramarginal gyrus | -300 | 27.7533 | -0.39415 | 0.11116 |
| Right SOG superior occipital gyrus | -47.779 | 51.9886 | -0.1022 | 0.12563 |
| Left SOG superior occipital gyrus | -20.1313 | 186.27 | -0.08898 | 0.18213 |
| Right SPL superior parietal lobule | -283.018 | 19.7144 | -0.21697 | 0.10935 |

| | | | | |
|---|---|---|---|---|
| Left SPL superior parietal lobule | -228.822 | 42.5037 | -0.20604 | 0.12639 |
| Right STG superior temporal gyrus | -285.959 | 59.4549 | -0.22136 | 0.14333 |
| Left STG superior temporal gyrus | -232.898 | 45.5607 | -0.20047 | 0.13356 |
| Right TMP temporal pole | -300 | -330 | -0.2528 | -0.08413 |
| Left TMP temporal pole | -300 | -263.576 | -0.24616 | -0.05492 |
| Right TrIFG triangular part of the inferior frontal gyrus | 6.09269 | 7.36744 | -0.02755 | 0.08494 |
| Left TrIFG triangular part of the inferior frontal gyrus | 0.42304 | 21.9329 | 0 | 0.10336 |
| Right TTG transverse temporal gyrus | -5.83701 | 11.5266 | -0.08222 | 0.09567 |
| Left TTG transverse temporal gyrus | 0.38757 | 1.90911 | 0 | 0 |

**eTable 5: Clinical association studies for the general population from UKBB.** Each clinical variable/biomarker was fit with a linear regression model by controlling covariates. $-\log_{10}$(P-value) and the coefficient of each variable/biomarker (beta) are reported. For the P-values, we added the sign of the variable's beta coefficient values to indicate the effect's direction. The threshold for Bonferroni corrected $-\log_{10}$(P-value) is > 3.08.

| Variable | R1: sign[$-\log_{10}$(P-value)] | R2: sign[$-\log_{10}$(P-value)] |
|---|---|---|
| Cholesterol | -1.24939 | -0.38153 |
| Lipoprotein A | -0.53029 | 0.04826 |
| HDL cholesterol | -3.9835 | -1.84715 |
| Triglycerides | 13.1313 | 0.0063 |
| Apolipoprotein A | -0.80123 | -1.81085 |
| Apolipoprotein B | -0.20008 | 0.1776 |
| C-reactive protein | 3.58939 | 0.14067 |
| LDL direct | -1.6734 | -0.00117 |
| Glucose | 7.08853 | -0.86482 |
| Glycated haemoglobin (HbA1c) | 14.2612 | -1.67051 |
| Maternal smoking | 0.78205 | -0.77154 |
| Household income | 0.07764 | 2.15911 |
| Education | 3.18428 | 2.89039 |
| Ethnicity | 0.63381 | 0.29638 |
| Townsend Deprivation Index tertiles | 2.92812 | 1.18548 |
| Sleep duration | 0.70548 | 0.29459 |
| Easy to get up in the morning | -1.88294 | 3.93415 |
| Morning or evening person | 1.17031 | -1.66819 |
| Nap during the day | 0.66029 | -0.6407 |
| Insomnia | -0.05737 | -2.81902 |
| Snoring | 0.05859 | -0.11376 |
| Daytime dozing sleeping | -0.5191 | 0.15494 |
| Sleep too much | 2.24252 | 0.2305 |
| Sleep trouble start | -0.14304 | -0.0328 |
| Sleep trouble end | -2.39029 | -0.09417 |
| Sleep any problem | 0.55325 | -0.31399 |
| Ever smoked | -0.18363 | -0.28497 |
| Smoking status | 0.74339 | -0.20541 |
| Current tobacco smoking | 2.64765 | 0.10738 |
| Past tobacco smoking | 1.72746 | -0.26343 |
| Light smokers at least 100 smokes in lifetime | -0.66022 | 1.70368 |
| Age started smoking in former smokers | -1.40981 | 0.05561 |
| Age stopped smoking | 9.72342 | -0.73592 |

| | | |
|---|---|---|
| Smoking smokers in household | 0.30144 | 0.29766 |
| Exposure to tobacco smoke at home | 0.20145 | -0.10873 |
| Exposure to tobacco smoke outside home | -0.09736 | 0.22288 |
| Coffee intake | -0.60155 | -2.47628 |
| Ever vascular heart problem diagnosed | -5.01173 | -0.09841 |
| Age high blood pressure diagnosed | -1.32468 | -0.7335 |
| Diabetes diagnosed by doctor | 21.3642 | -2.04208 |
| Diastolic blood pressure (automated reading) | 10.5304 | 1.43241 |
| Pulse rate (automated reading) | 1.26898 | 0.13024 |
| Systolic blood pressure (automated reading) | 1.83095 | 0.16135 |
| Pair matching_1 | 6.51554 | 0.63956 |
| Pair matching_2 | 7.76305 | 1.38813 |
| Reaction time_1 | 2.32465 | 5.09847 |
| Reaction time_2 | 0.43728 | 0.68777 |
| VNR | -17.0222 | -0.78106 |
| Prospective memory_1 | 1.46794 | -0.65534 |
| Prospective memory_2 | 2.34823 | 8.31264 |
| Pair_matching_2 (online) | 0.9509 | 0.25269 |
| Pair_matching_3 (online) | 0.7003 | 0.92109 |
| Symbolic digital substitution_1 (online) | -7.16303 | -0.63596 |
| Symbolic digital substitution_2 (online) | 2.12657 | 3.19173 |
| VNR (online) | -7.91013 | -1.46754 |
| TMT-A | 3.02215 | -0.08899 |
| TMT-B | 8.47316 | -2.29301 |
| AD_by_proxy | -2.24382 | 0.12213 |
| SPARE_AD | 136 | 250 |
| SPARE_BA | 236.859 | -46.3563 |
| WMLS_Volume_701 | 120.241 | 2.06141 |

**eTable 6: Identified genomic loci and mapped genes for GWAS in the general population**

**R1:**

| Locus | P-value | Chromosome | Mapped genes |
|---|---|---|---|
| rs13079098 | 4.91E-09 | 3 | *MAP4, PTPN23, KIF9, COL7A1, CSPG5, KLHL18, SCAP, DHX30, NBEAL2, CCDC12, ELP6, CDC25A, ZNF589,NME6, SMARCC1, SETD2,TREX1* |
| rs17669337 | 4.51E-11 | 5 | -- |
| rs2028849 | 8.76E-10 | 8 | *ENPP2* |
| rs11794318 | 3.02E-09 | 9 | *ASTN2* |
| rs3825045 | 1.98E-08 | 11 | *ZBTB16, NNMT* |
| rs9795600 | 1.12E-08 | 12 | *ANAPC5, KDM2B, MORN3, RNF34* |
| rs11158204, rs5809016 | 1.31E-15, 1.30E-15 | 14 | *GPR135, TOMM20L, AL132989.1, C14orf37, L3HYPDH, ACTR10, TIMM9, KIAA0586, DAAM1, ARID4A, PSMA3* |
| rs34395183 | 3.70E-08 | 17 | *RHOT1* |
| rs68175985 | 2.11E-08 | 19 | *NFIX* |
| rs201662360 | 2.41E-09 | 20 | *MYH7B, EDEM2, TRPC4AP, UQCC1, MAP1LC3A, GDF5, ERGIC3, FAM83C, ACSS2, GDF5OS, CPNE1* |

**R2:**

| Locus | P-value | Chromosome | Mapped genes |
|---|---|---|---|
| rs76606504 | 4.48E-08 | 1 | *EPS15, CDKN2C, DMRTA2, FAF1* |
| rs3860446, rs10184573 | 3.02E-09, 2.09E-08 | 2 | -- |
| rs309587 | 4.06E-09 | 5 | *VCAN* |
| rs117892760 | 1.68E-17 | 6 | *CENPW, HINT3, NCOA7, TRMT11* |
| rs55885771 | 1.88E-08 | 7 | *CAV2, TES* |
| rs1475536, rs2416560 | 3.82E-13, 2.16E-08 | 9 | *GADD45G* |
| rs58163343 | 2.10E-08 | 11 | *MDK, AMBRA1, DGKZ,CREB3L1, ARHGAP1, ATG13, CHST1, HARBI1* |
| rs10774183, rs17178006, rs77956314 | 9.22E-20, 5.30E-29, 9.98E-29 | 12 | *TESC, RFC5, HRK, RNFT2, WIF1, MSRB3, LEMD3, FBXW8* |
| rs143383 | 5.46E-11 | 20 | *CPNE1, GDF5OS, ACSS2, GDF5, ERGIC3, ERGIC3, MAP1LC3A, UQCC1, TRPC4AP, EDEM2, MYH7B* |

#### eTable 7: Gene set enrichment analysis results in the general population

## R1:

| Category | Gene Set | P-value | Overlapped genes |
|---|---|---|---|
| GWAS Catalog | Waist-to-hip ratio | 2.23E-11 | *ACSS2:MYH7B:TRPC4AP:EDEM2: FAM83C:UQCC1:GDF5:ERGIC3:CPNE1* |
| GWAS Catalog | Huntington's disease | 5.63E-09 | *CCDC12:NBEAL2:SETD2:KIF9:KLHL18* |
| GWAS Catalog | Prothrombin time | 8.07E-07 | *MYH7B:TRPC4AP:EDEM2* |
| GWAS Catalog | QRS | 4.87E-05 | *MYH7B:EDEM2* |
| GWAS Catalog | Spine bone | 7.82E-05 | *UQCC1:GDF5* |
| GWAS Catalog | Height | 1.79E-04 | *FAM83C:UQCC1:GDF5OS:GDF5:ERGIC3: CPNE1:ENPP2* |
| GWAS Catalog | Blood protein | 1.99E-04 | *MYH7B:TRPC4AP:EDEM2:UQCC1:GDF5OS: GDF5:CPNE1:CDC25A:ZNF589:COL7A1* |
| Chemical and Genetic perturbation | NIKOLSKY BREAST CANCER 20Q11 AMPLICON | 2.84E-22 | *MAP1LC3A:ACSS2:MYH7B:TRPC4AP: EDEM2:FAM83C:UQCC1:GDF5:ERGIC3:CPNE1* |
| Chemical and Genetic perturbation | LASTOWSKA NEUROBLASTOMA COPY NUMBER DN | 9.36E-07 | *CCDC12:SETD2:SCAP:ELP6:SMARCC1: DHX30:CDC25A:ZNF589:NME6* |
| Chemical and Genetic perturbation | PURBEY TARGETS OF CTBP1 AND SATB1 UP | 3.54E-06 | *ARID4A:GPR135:CPNE1:COL7A1* |
| Chemical and Genetic perturbation | STREICHER LSM1 TARGETS UP | 3.05E-05 | *ARID4A:DAAM1:CDC25A* |
| Immunologic signatures | GSE25123 CTRL VS IL4 AND ROSIGLITAZONE STIM PPARG KO MACROPHAGE DN | 7.17E-06 | *ZBTB16:ANAPC5:NFIX:ACSS2:KLHL18* |
| Curated gene sets | NIKOLSKY BREAST CANCER 20Q11 AMPLICON | 2.84E-22 | *MAP1LC3A:ACSS2:MYH7B:TRPC4AP:EDEM2: FAM83C:UQCC1:GDF5:ERGIC3:CPNE1* |
| Curated gene sets | LASTOWSKA NEUROBLASTOMA COPY NUMBER DN | 9.36E-07 | *CCDC12:SETD2:SCAP:ELP6:SMARCC1: DHX30:CDC25A:ZNF589:NME6* |
| Curated gene sets | PURBEY TARGETS OF CTBP1 AND SATB1 UP | 3.53E-06 | *ARID4A:GPR135:CPNE1:COL7A1* |
| Curated gene sets | REACTOME CHROMATIN ORGANIZATION | 2.93E-05 | *KDM2B:ARID4A:SETD2:ELP6:SMARCC1* |
| Curated gene sets | STREICHER LSM1 TARGETS UP | 3.05E-05 | *ARID4A:DAAM1:CDC25A* |
| Reactome | REACTOME CHROMATIN ORGANIZATION | 2.93E-05 | *KDM2B:ARID4A:SETD2:ELP6:SMARCC1* |

## R2:

| Category | Gene Set | P-value | Overlapped genes |
|---|---|---|---|
| GWAS Catalog | Hippocampus | 2.19E-22 | *WIF1:LEMD3:MSRB3:RNFT2:HRK:FBXW8:TESC:ASTN2* |
| GWAS Catalog | Dentate gyrus | 2.19E-22 | *WIF1:LEMD3:MSRB3:RNFT2:HRK:FBXW8:TESC:ASTN2* |
| GWAS Catalog | Alzheimer's disease | 1.36E-13 | *CREB3L1:DGKZ:MDK:AMBRA1:HARBI1:ATG13:ARHGAP1* |
| GWAS Catalog | Waist-to-hip ratio | 4.66E-12 | *ACSS2:MYH7B:TRPC4AP:EDEM2:FAM83C:UQCC1: GDF5:ERGIC3:CPNE1* |
| GWAS Catalog | Immunoglobulin A | 2.95E-11 | *DGKZ:MDK:AMBRA1:HARBI1:ATG13* |
| GWAS Catalog | Baldness | 4.19E-07 | *DMRTA2:FAF1:CDKN2C:HINT3:TRMT11:CENPW* |
| GWAS Catalog | Prothrombin time | 4.92E-07 | *MYH7B:TRPC4AP:EDEM2* |
| GWAS Catalog | Schizophrenia | 3.59E-07 | *CREB3L1:DGKZ:MDK:AMBRA1:HARBI1:ATG13:ARHGAP1:ERGIC3* |
| GWAS Catalog | Autism spectrum disorder | 1.74E-05 | *CREB3L1:DGKZ:MDK:AMBRA1:HARBI1:ATG13:ARHGAP1* |
| GWAS Catalog | Reaction time | 2.42E-05 | *DGKZ:ATG13:ARHGAP1* |
| GWAS Catalog | QRS | 3.52E-05 | *MYH7B:EDEM2* |
| GWAS Catalog | FEV1 | 4.72E-05 | *FAF1:MSRB3:UQCC1:ASTN2* |
| GWAS Catalog | Spine bone | 5.65E-05 | *UQCC1:GDF5* |
| GWAS Catalog | Height | 6.20E-05 | *FAM83C:UQCC1:GDF5OS:GDF5:ERGIC3:CPNE1:CENPW* |
| GWAS Catalog | Ischemic stroke | 1.13E-04 | *FAF1:CDKN2C* |
| GWAS Catalog | Alzheimer's disease (onset) | 1.69E-04 | *RNFT2:HRK* |
| GWAS Catalog | Neuroticism | 1.98E-04 | *ARHGAP1:RFC5:CENPW* |
| Chemical and Genetic perturbation | NIKOLSKY BREAST CANCER 20Q11 AMPLICON | 4.67E-23 | *MAP1LC3A:ACSS2:MYH7B:TRPC4AP:EDEM2 :FAM83C:UQCC1:GDF5:ERGIC3:CPNE1* |
| Chemical and Genetic perturbation | ROVERSI GLIOMA_COPY_NUMBER_UP | 9.75E-08 | *CHST1:CREB3L1:DGKZ:MDK:ARHGAP1* |
| Chemical and Genetic perturbation | CHICAS RB1 TARGETS CONFLUENT | 3.04E-06 | *CREB3L1:MDK:ARHGAP1:MSRB3:GDF5:TES:CAV2* |
| Chemical and Genetic perturbation | WANG CLIM2 TARGETS UP | 9.95E-06 | *DGKZ:ATG13:UQCC1:CAV2:ASTN2* |
| Chemical and Genetic perturbation | HADDAD B LYMPHOCYTE PROGENITOR | 1.62E-05 | *CDKN2C:RNFT2:HRK:RFC5:GADD45G* |
| Immunologic signatures | GSE7509 UNSTIM VS TNFA IL1B IL6 PGE STIM DC UP | 7.62E-05 | *MAP1LC3A:TRPC4AP:EDEM2:HINT3* |
| Immunologic signatures | GSE17721 CTRL VS LPS 6H BMDC UP | 7.92E-05 | *FAF1:CDKN2C:EPS15:GADD45G* |
| Immunologic signatures | GSE17721 CTRL VS CPG 1H BMDC UP | 7.92E-05 | *CDKN2C:CREB3L1:DGKZ:GADD45G* |
| Immunologic signatures | GSE5960 TH1 VS ANERGIC TH1 DN | 7.92E-05 | *CREB3L1:MDK:TRPC4AP:TES* |
| Curated gene sets | NIKOLSKY BREAST CANCER 20Q11 AMPLICON | 4.67E-23 | *MAP1LC3A:ACSS2:MYH7B:TRPC4AP:EDEM2: FAM83C:UQCC1:GDF5:ERGIC3:CPNE1* |
| Curated gene sets | CHICAS RB1 TARGETS CONFLUENT | 3.04E-06 | *CREB3L1:MDK:ARHGAP1:MSRB3:GDF5:TES:CAV2* |
| Curated gene sets | WANG CLIM2 TARGETS UP | 9.96E-06 | *DGKZ:ATG13:UQCC1:CAV2:ASTN2* |
| Curated gene sets | HADDAD B LYMPHOCYTE PROGENITOR | 1.62E-05 | *CDKN2C:RNFT2:HRK:RFC5:GADD45G* |

**eTable 8: Brain association studies for the cognitively unimpaired (CU) population from ADNI and BLSA.** Each ROI was fit with a linear regression model by controlling covariates. –$\log_{10}$(P-value) and Pearson's correlation (*r*) are reported. For the P-values, we also added the sign of the ROI's beta coefficient values to indicate the effect's direction. The threshold for Bonferroni corrected –$\log_{10}$(P-value) is > 3.38. For the *r* values, the ROIs that did not survive the multiple comparisons or showed a positive *r* (negative *r* means brain atrophy) are assigned a value of 0 for visualization purposes.

| ROI | R1: sign[–$\log_{10}$(P-value)] | R2: sign[–$\log_{10}$(P-value)] | R1: *r* | R2: *r* |
|---|---|---|---|---|
| Right Accumbens Area | 0.02422 | -9.97664 | 0 | -0.01843 |
| Left Accumbens Area | -0.74491 | -5.85925 | 0 | 0.02215 |
| Right Amygdala | -2.48324 | -116.367 | 0 | -0.32023 |
| Left Amygdala | -0.49706 | -128.307 | 0 | -0.35099 |
| Right Caudate | 13.1087 | 0.31537 | 0.11551 | 0 |
| Left Caudate | 13.4701 | -0.09992 | 0.12082 | 0 |
| Right Cerebellum Exterior | -1.22265 | -3.19868 | 0 | 0 |
| Left Cerebellum Exterior | -2.79468 | -3.67638 | 0 | -0.02158 |
| Right Hippocampus | -3.06027 | -170.382 | 0 | -0.41714 |
| Left Hippocampus | -1.10676 | -155.177 | 0 | -0.37296 |
| Right Pallidum | 0.58246 | 0.34191 | 0 | 0 |
| Left Pallidum | 0.24224 | -0.00182 | 0 | 0 |
| Right Putamen | -1.31769 | -1.29282 | 0 | 0 |
| Left Putamen | -0.65712 | -1.74109 | 0 | 0 |
| Right Thalamus Proper | -0.79193 | -3.88437 | 0 | 0.02803 |
| Left Thalamus Proper | -1.17706 | -3.62831 | 0 | 0.03503 |
| Cerebellar Vermal Lobules I-V | -4.13432 | -0.07438 | -0.08298 | 0 |
| Cerebellar Vermal Lobules VI-VII | 0.66074 | 1.69841 | 0 | 0 |
| Cerebellar Vermal Lobules VIII-X | -0.13531 | -3.29106 | 0 | 0 |
| Left Basal Forebrain | 1.52331 | -4.46827 | 0 | 0.01873 |
| Right Basal Forebrain | 1.44434 | -2.36463 | 0 | 0 |
| Right ACgG anterior cingulate gyrus | -0.14346 | 0.11605 | 0 | 0 |
| Left ACgG anterior cingulate gyrus | -0.47404 | 2.92022 | 0 | 0 |
| Right AIns anterior insula | -6.19075 | -0.90888 | -0.09814 | 0 |
| Left AIns anterior insula | -3.9446 | -1.54477 | -0.08014 | 0 |
| Right AOrG anterior orbital gyrus | -0.25551 | 0.10997 | 0 | 0 |
| Left AOrG anterior orbital gyrus | -0.0104 | 0.51573 | 0 | 0 |
| Right AnG angular gyrus | -44.3189 | 2.29514 | -0.25141 | 0 |
| Left AnG angular gyrus | -34.0593 | 2.69873 | -0.23458 | 0 |
| Right Calc calcarine cortex | 0.95427 | 1.3806 | 0 | 0 |
| Left Calc calcarine cortex | 1.60106 | 2.3469 | 0 | 0 |
| Right CO central operculum | -21.4518 | 3.68169 | -0.1368 | 0.17626 |
| Left CO central operculum | -30.7951 | 2.13474 | -0.21934 | 0 |
| Right Cun cuneus | -1.30454 | 0.15463 | 0 | 0 |
| Left Cun cuneus | -0.85633 | 0.089 | 0 | 0 |
| Right Ent entorhinal area | -0.48193 | -94.2836 | 0 | -0.31755 |
| Left Ent entorhinal area | -1.47259 | -112.902 | 0 | -0.36522 |

| | | | | |
|---|---|---|---|---|
| Right FO frontal operculum | -4.71612 | 3.50542 | -0.08757 | 0.14387 |
| Left FO frontal operculum | -7.80919 | 4.71573 | -0.06409 | 0.18985 |
| Right FRP frontal pole | -1.78447 | 7.39516 | 0 | 0.1951 |
| Left FRP frontal pole | -6.25577 | 2.96574 | -0.0413 | 0 |
| Right FuG fusiform gyrus | -11.3483 | -3.99863 | -0.16477 | -0.02522 |
| Left FuG fusiform gyrus | -20.5362 | -4.72936 | -0.20459 | -0.0245 |
| Right GRe gyrus rectus | -0.99644 | 0.36732 | 0 | 0 |
| Left GRe gyrus rectus | 0.08139 | -0.71079 | 0 | 0 |
| Right IOG inferior occipital gyrus | -0.56007 | 0.76815 | 0 | 0 |
| Left IOG inferior occipital gyrus | 0.10238 | 3.30977 | 0 | 0 |
| Right ITG inferior temporal gyrus | -79.5925 | -0.43182 | -0.36824 | 0 |
| Left ITG inferior temporal gyrus | -54.6705 | -2.45957 | -0.33551 | 0 |
| Right LiG lingual gyrus | -1.10594 | -1.03965 | 0 | 0 |
| Left LiG lingual gyrus | 0.21369 | -0.22744 | 0 | 0 |
| Right LOrG lateral orbital gyrus | 5.96812 | 7.65209 | 0.03279 | 0.11501 |
| Left LOrG lateral orbital gyrus | 1.44208 | 8.75206 | 0 | 0.19311 |
| Right MCgG middle cingulate gyrus | 0.07973 | 3.16878 | 0 | 0 |
| Left MCgG middle cingulate gyrus | -0.05795 | 4.14204 | 0 | 0.13171 |
| Right MFC medial frontal cortex | 1.08139 | -1.51896 | 0 | 0 |
| Left MFC medial frontal cortex | 3.58839 | -0.78213 | 0.05774 | 0 |
| Right MFG middle frontal gyrus | -2.90496 | 15.8988 | 0 | 0.18026 |
| Left MFG middle frontal gyrus | -3.27888 | 15.1054 | 0 | 0.18331 |
| Right MOG middle occipital gyrus | -13.2238 | 2.98902 | -0.18857 | 0 |
| Left MOG middle occipital gyrus | -22.4568 | 0.45539 | -0.21777 | 0 |
| Right MOrG medial orbital gyrus | 2.26776 | 3.89384 | 0 | 0.11455 |
| Left MOrG medial orbital gyrus | 7.10968 | 3.86849 | 0.00986 | 0.07279 |
| Right MPoG postcentral gyrus medial segment | 4.26336 | 2.28323 | 0.08468 | 0 |
| Left MPoG postcentral gyrus medial segment | -0.976 | 0.98263 | 0 | 0 |
| Right MPrG precentral gyrus medial segment | 1.7376 | 2.1092 | 0 | 0 |
| Left MPrG precentral gyrus medial segment | 3.86669 | 3.14416 | 0.03619 | 0 |
| Right MSFG superior frontal gyrus medial segment | -1.9857 | 3.71251 | 0 | 0.15299 |
| Left MSFG superior frontal gyrus medial segment | -3.80664 | 1.50272 | -0.03031 | 0 |
| Right MTG middle temporal gyrus | -78.7893 | -1.42619 | -0.32799 | 0 |
| Left MTG middle temporal gyrus | -88.6047 | -3.09453 | -0.35737 | 0 |
| Right OCP occipital pole | 9.10347 | 1.57333 | 0.08553 | 0 |
| Left OCP occipital pole | 11.9957 | 7.11525 | 0.12456 | 0.1673 |
| Right OFuG occipital fusiform gyrus | -0.44305 | 1.93927 | 0 | 0 |

| | | | | |
|---|---|---|---|---|
| Left OFuG occipital fusiform gyrus | -7.02536 | -2.44769 | -0.14617 | 0 |
| Right OpIFG opercular part of the inferior frontal gyrus | -11.9213 | 4.95626 | -0.13442 | 0.16631 |
| Left OpIFG opercular part of the inferior frontal gyrus | -11.1503 | 2.94214 | -0.10786 | 0 |
| Right OrIFG orbital part of the inferior frontal gyrus | 0.70391 | 2.36786 | 0 | 0 |
| Left OrIFG orbital part of the inferior frontal gyrus | 1.81898 | 2.94668 | 0 | 0 |
| Right PCgG posterior cingulate gyrus | -15.3029 | -1.77989 | -0.19969 | 0 |
| Left PCgG posterior cingulate gyrus | -11.5197 | -0.61498 | -0.17758 | 0 |
| Right PCu precuneus | -29.9026 | 2.54008 | -0.2271 | 0 |
| Left PCu precuneus | -19.8806 | 1.80214 | -0.17966 | 0 |
| Right PHG parahippocampal gyrus | -16.582 | -109.089 | -0.16942 | -0.32044 |
| Left PHG parahippocampal gyrus | -11.9967 | -127.379 | -0.16249 | -0.35343 |
| Right PIns posterior insula | -12.4309 | -0.60413 | -0.14632 | 0 |
| Left PIns posterior insula | -19.8889 | -2.07986 | -0.1697 | 0 |
| Right PO parietal operculum | -1.49742 | 3.12846 | 0 | 0 |
| Left PO parietal operculum | -2.05314 | 5.69654 | 0 | 0.14372 |
| Right PoG postcentral gyrus | -11.147 | 15.9636 | -0.14717 | 0.22742 |
| Left PoG postcentral gyrus | -7.26167 | 6.01063 | -0.13704 | 0.15154 |
| Right POrG posterior orbital gyrus | 1.78048 | 0.45129 | 0 | 0 |
| Left POrG posterior orbital gyrus | 3.11751 | 0.99777 | 0 | 0 |
| Right PP planum polare | -10.2705 | 0.14541 | -0.06999 | 0 |
| Left PP planum polare | -2.33648 | 1.43902 | 0 | 0 |
| Right PrG precentral gyrus | -1.58787 | 19.0602 | 0 | 0.23317 |
| Left PrG precentral gyrus | -1.27245 | 23.2786 | 0 | 0.23342 |
| Right PT planum temporale | -7.95868 | 5.77427 | -0.08165 | 0.18999 |
| Left PT planum temporale | -13.6171 | 0.3216 | -0.16147 | 0 |
| Right SCA subcallosal area | 12.1687 | 3.36894 | 0.02896 | 0 |
| Left SCA subcallosal area | 11.271 | 1.70392 | 0.01615 | 0 |
| Right SFG superior frontal gyrus | -0.74848 | 11.4176 | 0 | 0.19607 |
| Left SFG superior frontal gyrus | -1.90879 | 11.9413 | 0 | 0.17828 |
| Right SMC supplementary motor cortex | 1.36407 | 2.92348 | 0 | 0 |
| Left SMC supplementary motor cortex | 0.28531 | 1.30349 | 0 | 0 |
| Right SMG supramarginal gyrus | -47.9769 | 3.07503 | -0.33785 | 0 |
| Left SMG supramarginal gyrus | -60.9529 | 3.41068 | -0.37277 | 0.10429 |
| Right SOG superior occipital gyrus | -2.9757 | 0.45344 | 0 | 0 |
| Left SOG superior occipital gyrus | -0.70766 | 3.84876 | 0 | 0.12067 |

| | | | | |
|---|---|---|---|---|
| Right SPL superior parietal lobule | -7.51241 | 1.52569 | -0.17257 | 0 |
| Left SPL superior parietal lobule | -12.5112 | 1.79292 | -0.19693 | 0 |
| Right STG superior temporal gyrus | -19.5533 | 0.52986 | -0.18466 | 0 |
| Left STG superior temporal gyrus | -14.7256 | 0.67456 | -0.18506 | 0 |
| Right TMP temporal pole | -47.7076 | -21.6918 | -0.28679 | -0.12042 |
| Left TMP temporal pole | -38.7424 | -21.3961 | -0.26189 | -0.12065 |
| Right TrIFG triangular part of the inferior frontal gyrus | 0.86865 | 1.19197 | 0 | 0 |
| Left TrIFG triangular part of the inferior frontal gyrus | 0.33749 | 0.66059 | 0 | 0 |
| Right TTG transverse temporal gyrus | -0.73044 | 0.83119 | 0 | 0 |
| Left TTG transverse temporal gyrus | 0.26298 | 0.05257 | 0 | 0 |

**eTable 9: Clinical association studies for the cognitively unimpaired (CU) population from ADNI and BLSA.** Each clinical variable/biomarker was fit with a linear regression model by controlling covariates. $-\log_{10}$(P-value) and the coefficient of each variable/biomarker (beta) are reported. For the P-values, we added the sign of the variable's beta coefficient values to indicate the effect's direction. The threshold for Bonferroni corrected $-\log_{10}$(P-value) is > 2.95..

| Variable | R1: sign[$-\log_{10}$(P-value)] | R2: sign[$-\log_{10}$(P-value)] |
|---|---|---|
| FAQ_Total | 0.03592 | 0.1324 |
| CDR_Global | 0.2549 | 0.68368 |
| CDR_SOB | 0.46167 | 0.38305 |
| NPIQ_Total | -0.02945 | -1.85116 |
| DSST | 0.79779 | -0.17017 |
| TMT_A | 1.04091 | 0.26962 |
| TMT_B | 0.38981 | 1.39854 |
| MMSE | -0.42715 | -0.69585 |
| Digit_Span_Backward | -1.11892 | -0.4529 |
| Digit_Span_Forward | -0.5489 | 0.15144 |
| MOCA | -0.1209 | 0.046 |
| BNT | 0.18498 | -0.7789 |
| RAVLT | 0.75931 | -0.16779 |
| RAVLT_IM | 1.02925 | 0.03534 |
| RAVLT_Short | 0.41472 | -0.42162 |
| RAVLT_Long | 0.35987 | -0.51004 |
| LM_DEL_A | 0.05756 | 0.17738 |
| LM_IM_A | -0.26376 | 0.45558 |
| ANI_Fluency | 0.07071 | 0.20993 |
| VEG_Fluency | -0.02664 | 0.40168 |
| TS_RATIO | 1.86587 | -0.00366 |
| TS_RATIO_ADJ | 1.97264 | -0.12439 |
| BNT_percent | 0.18498 | -0.7789 |
| Cholesterol | -1.56606 | -0.18187 |
| Diastole | 0.3944 | -0.05256 |
| Glucose | 0.67578 | -0.66702 |
| Systole | 0.84884 | -0.22012 |
| Triglycerides | -0.23883 | -0.35205 |
| ABETA42_ADNI | -0.15777 | -0.07168 |
| PTAU | -0.30024 | 1.00901 |
| TAU | -0.29778 | 0.82113 |
| Diabetes | 1.08237 | 0.02205 |
| Hypertension | -0.28435 | -0.28563 |
| Diagnosis_Depression | -0.59092 | 0.04985 |
| Alcoholic | -0.22664 | 2.69173 |
| Family_History_Dementia | -0.58766 | 0.17379 |
| Ethnicity | 1.44783 | 0.02901 |
| Height | -0.01913 | -0.29661 |
| BMI | 0.01531 | -0.98463 |
| Weight | 0.01957 | -0.95236 |
| SPARE_AD | 75.0432 | 138.158 |
| SPARE_BA | 14.9898 | -3.16218 |
| AV45_PET_ADNI | 0.42998 | 1.17676 |
| FDG | 0.58558 | -1.02347 |
| WMLS_Volume_701 | 6.92099 | 1.98594 |

### References


1. Yang, Z., Wen, J. & Davatzikos, C. Surreal-GAN:Semi-Supervised Representation Learning via GAN for uncovering heterogeneous disease-related imaging patterns. *ICLR* (2021).
2. Yang, Z. *et al.* A deep learning framework identifies dimensional representations of Alzheimer's Disease from brain structure. *Nat Commun* **12**, 7065 (2021).
3. Wen, J. *et al.* Multi-scale semi-supervised clustering of brain images: Deriving disease subtypes. *Med Image Anal* **75**, 102304 (2021).
4. Dong, A., Honnorat, N., Gaonkar, B. & Davatzikos, C. CHIMERA: Clustering of Heterogeneous Disease Effects via Distribution Matching of Imaging Patterns. *IEEE Trans. Med. Imaging* **35**, 612–621 (2016).
5. Varol, E., Sotiras, A. & Davatzikos, C. HYDRA: Revealing heterogeneity of imaging and genetic patterns through a multiple max-margin discriminative analysis framework. *NeuroImage* **145**, 346–364 (2017).
6. Wen, J. *et al.* Subtyping brain diseases from imaging data. Preprint at https://doi.org/10.48550/arXiv.2202.10945 (2022).
7. Young, A. L. *et al.* Uncovering the heterogeneity and temporal complexity of neurodegenerative diseases with Subtype and Stage Inference. *Nat Commun* **9**, 4273 (2018).
8. Liu, J. Z., Erlich, Y. & Pickrell, J. K. Case-control association mapping by proxy using family history of disease. *Nat Genet* **49**, 325–331 (2017).
9. Jansen, I. E. *et al.* Genome-wide meta-analysis identifies new loci and functional pathways influencing Alzheimer's disease risk. *Nat Genet* **51**, 404–413 (2019).

10. Manichaikul, A. *et al.* Robust relationship inference in genome-wide association studies. *Bioinformatics* **26**, 2867–2873 (2010).

11. Price, A. L., Zaitlen, N. A., Reich, D. & Patterson, N. New approaches to population stratification in genome-wide association studies. *Nat Rev Genet* **11**, 459–463 (2010).

12. Price, A. L. *et al.* Principal components analysis corrects for stratification in genome-wide association studies. *Nat Genet* **38**, 904–909 (2006).

13. Wen, J. *et al.* Characterizing Heterogeneity in Neuroimaging, Cognition, Clinical Symptoms, and Genetics Among Patients With Late-Life Depression. *JAMA Psychiatry* (2022) doi:10.1001/jamapsychiatry.2022.0020.

14. Wen, J. *et al.* Novel genomic loci and pathways influence patterns of structural covariance in the human brain. 2022.07.20.22277727 Preprint at https://doi.org/10.1101/2022.07.20.22277727 (2022).

15. Abraham, G., Qiu, Y. & Inouye, M. FlashPCA2: principal component analysis of Biobank-scale genotype datasets. *Bioinformatics* **33**, 2776–2778 (2017).

16. Wen, J. *et al.* The Genetic Architecture of Biological Age in Nine Human Organ Systems. *medRxiv* 2023.06.08.23291168 (2023) doi:10.1101/2023.06.08.23291168.

17. Watanabe, K., Taskesen, E., van Bochoven, A. & Posthuma, D. Functional mapping and annotation of genetic associations with FUMA. *Nat Commun* **8**, 1826 (2017).

18. The GTEx Consortium. The Genotype-Tissue Expression (GTEx) project. *Nat Genet* **45**, 580–585 (2013).

19. Choi, S. W., Mak, T. S.-H. & O'Reilly, P. F. Tutorial: a guide to performing polygenic risk score analyses. *Nat Protoc* **15**, 2759–2772 (2020).

20. Lambert, J.-C. *et al.* Meta-analysis of 74,046 individuals identifies 11 new susceptibility loci for Alzheimer's disease. *Nat Genet* **45**, 1452–1458 (2013).

21. Bulik-Sullivan, B. K. *et al.* LD Score regression distinguishes confounding from polygenicity in genome-wide association studies. *Nat Genet* **47**, 291–295 (2015).

22. International Human Genome Sequencing Consortium. Finishing the euchromatic sequence of the human genome. *Nature* **431**, 931–945 (2004).

23. Karolchik, D. *et al.* The UCSC Genome Browser Database. *Nucleic Acids Res* **31**, 51–54 (2003).

24. Wishart, D. S. *et al.* DrugBank 5.0: a major update to the DrugBank database for 2018. *Nucleic Acids Res* **46**, D1074–D1082 (2018).

25. Zhou, Y. *et al.* Therapeutic target database update 2022: facilitating drug discovery with enriched comparative data of targeted agents. *Nucleic Acids Res* **50**, D1398–D1407 (2022).

26. Petersen, R. C. *et al.* Alzheimer's Disease Neuroimaging Initiative (ADNI): clinical characterization. *Neurology* **74**, 201–209 (2010).

27. Resnick, S. M. *et al.* One-year age changes in MRI brain volumes in older adults. *Cereb Cortex* **10**, 464–472 (2000).

28. Miller, K. L. *et al.* Multimodal population brain imaging in the UK Biobank prospective epidemiological study. *Nat Neurosci* **19**, 1523–1536 (2016).

29. Buniello, A. *et al.* The NHGRI-EBI GWAS Catalog of published genome-wide association studies, targeted arrays and summary statistics 2019. *Nucleic Acids Res* **47**, D1005–D1012 (2019).

30. Davatzikos, C., Xu, F., An, Y., Fan, Y. & Resnick, S. M. Longitudinal progression of Alzheimer's-like patterns of atrophy in normal older adults: the SPARE-AD index. *Brain* **132**, 2026–2035 (2009).

31. Yang, J. *et al.* FTO genotype is associated with phenotypic variability of body mass index. *Nature* **490**, 267–272 (2012).
